# Dynamics of bacterial recombination in the human gut microbiome

**DOI:** 10.1101/2022.08.24.505183

**Authors:** Zhiru Liu, Benjamin H Good

**Affiliations:** Department of Applied Physics, Stanford University, Stanford, CA 94305; Department of Biology, Stanford University, Stanford, CA 94305; Chan Zuckerberg Biohub – San Francisco, San Francisco, CA 94158

## Abstract

Horizontal gene transfer is a ubiquitous force in microbial evolution. Previous studies have shown that the human gut is a hot spot for gene transfer between species, but the more subtle exchange of variation within species – collectively known as recombination – remains poorly characterized in this ecosystem. Here, we show how the genetic structure of the human gut microbiome provides a unique opportunity to measure individual recombination events from sequenced fecal samples, enabling quantitative comparisons of recombination across a diverse range of species that inhabit a common environment. By analyzing a large collection of recent recombination events in the core genomes of 29 gut commensals, we uncovered systematic heterogeneities in the rates and lengths of transferred segments, which are difficult to explain by existing models of ecological isolation or homology-dependent recombination rates. We also find that natural selection plays a role in facilitating the spread of genetic variants onto different strain backgrounds, both within individual hosts and across the broader global population. These results shed light on the dynamics of *in situ* recombination, which place important constraints on the adaptability of gut microbial communities.

## Introduction

The horizontal exchange of genetic material – also known as horizontal gene transfer (HGT) – is a pervasive force in microbial ecology and evolution [1]. HGT is particularly important within the human gut microbiota, where hundreds of species coexist with each other in close physical proximity [2–4]. HGT is often associated with the acquisition of new genes or pathways, which can confer resistance to antibiotics [3–8] or enable novel metabolic capabilities [3, 9–14]. Genetic material can also be transferred between more closely related strains, where it can overwrite existing regions of the genome via homologous recombination [15, 16]. This more subtle form of horizontal exchange acts to reshuffle genetic variants within species, similar to meiotic recombination in sexual organisms. Homologous recombination plays a crucial role in microbial evolution, from the emergence of new bacterial species [17–20] to the transition between clonal and quasi-sexual evolution [21–23]. Homologous recombination can also serve as a scaffold for the incorporation of novel genetic material, which can facilitate the spread of accessory genes across different strain backgrounds [24]. However, while numerous studies have established the pervasiveness of bacterial recombination [21, 25–28], the evolutionary dynamics of this process are still poorly understood in natural populations like the gut microbiota.

Multiple methods have been developed for inferring *in situ* recombination from the fine-scale diversity of natural bacterial isolates [26, 27, 29–33]. The key challenge lies in disentangling the effects of recombination from the other evolutionary forces (e.g. mutation, selection, and genetic drift) that shape genetic diversity over the same timescales. Existing studies often address this problem using an inverse approach, by fitting the observed data to simple parametric models from microbial population genetics. Examples range from simple summary statistics like linkage disequilibrium [26, 27, 34, 35] and related metrics [21, 28, 32, 33, 36–40] to complete probabilistic reconstructions of the genealogies of the sampled genomes [29–31]. Previous applications of these methods have provided extensive evidence for ongoing recombination within the core genomes of many bacterial species [25, 41] – including many species of human gut bacteria [27].

However, many of these existing methods rely on simplified evolutionary scenarios, which ignore the effects of natural selection, and make additional restrictive assumptions about the demographic structure of the population. Recent work has shown that these simplified models often fail to capture key features of microbial genetic diversity [26–28, 42], which can strongly bias estimates of the underlying recombination parameters. Our limited understanding of these effects makes it difficult to answer key questions about the role of recombination in natural populations like the gut microbiota: is recombination fast enough to allow local adaptations to persist within a host, e.g. during fecal microbiota transplants [43] or sudden dietary shifts [44]? Does natural selection tend to promote or hinder the spread of genetic variants across different strain backgrounds? And can the rates and lengths of transferred fragments shed light on the underlying mechanisms of recombination *in situ*?

Here we show that the genetic structure of the human gut microbiome provides a unique opportunity to address these questions. Using strain-resolved metagenomics, we show that the large sample sizes and host colonization structure of this ecosystem enable systematic comparisons of strains across a broad range of distance- and time-scales, from the scale of individual hosts to the diversity of the broader global population. We show that some of these strains are closely related enough that one can resolve homologous recombination events directly, without requiring restrictive modeling assumptions or explicit phylogenetic inference. We use these observations to develop a non-parametric approach for identifying large numbers of recent recombination events within 29 prevalent species of human gut bacteria. This comparative dataset allows us to systematically explore the landscape of homologous recombination in this host-associated ecosystem.

Our results reveal extensive heterogeneity in rates and lengths of transferred fragments – both among different species and between different strains of the same species – which are difficult to explain by ecological isolation or reduced efficiencies of recombination. We also find that natural selection can play an important role in facilitating the spread of transferred fragments into different strain backgrounds. Our results suggest that *in situ* recombination events are shaped by a combination of evolutionary processes, which may strongly depend on the ecological context of their host community.

## Results

### Partially recombined genomes underlie the broad range of genetic diversity in many species of gut bacteria

To quantify the dynamics of homologous recombination across different timescales, we analyzed shotgun metagenomic data from a collection of healthy human gut microbiomes that we collated in a previous study [27]. This dataset comprises 932 fecal samples from 693 subjects from North America, Europe, and China (Table S1). We used a reference-based approach to identify single nucleotide variants (SNVs) in the core genome of each species in each sample (S1 Text 1). These metagenomic variants reflect a complex mixture of the global genetic diversity within a given species, as well as the specific combination of lineages that are present within a given host. While it is difficult to resolve the underlying lineages in the most general case, we previously showed [27] that the lineage structure in many human gut metagenomes is simple enough that the core genome of the dominant strain can be inferred with a high degree of confidence (Figs. 1A & S1). Using this approach, we obtained a total of 5416 “quasi-phased” genomes from 43 different species in 541 unique hosts. The genetic differences between these strains provide a window into the long-term evolutionary forces that operate in these species over multiple host colonization cycles.

**Fig 1.**
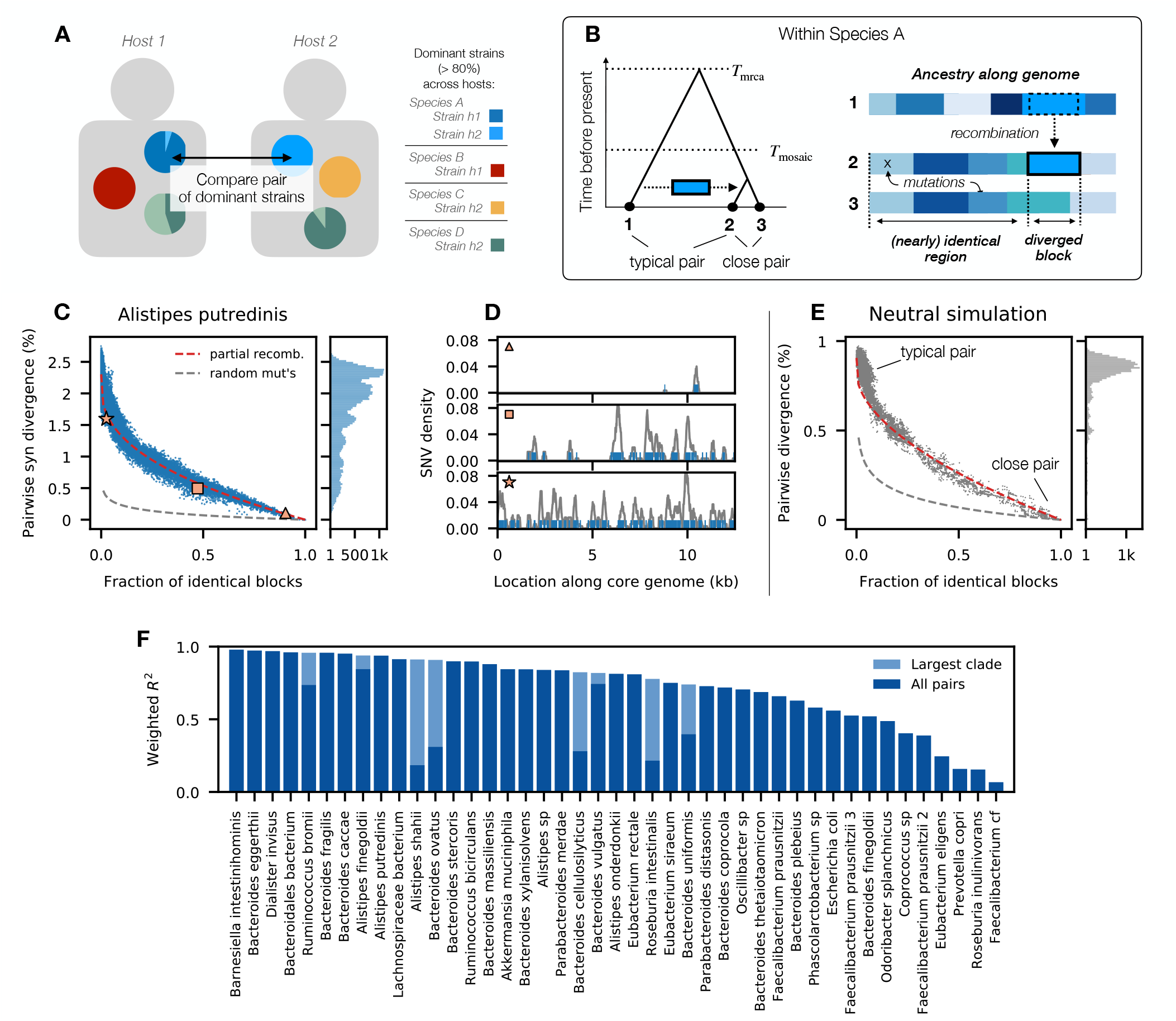
Partially recombined genomes underlie the broad range of genetic divergence in many species of gut bacteria. (A) Genetic differences between the core genomes of the dominant strain of a given species (>80% within-host frequency) are inferred from pairwise comparisons of metagenomes from unrelated hosts (S1 Text 1). (B) Timescales of recombination in a quasi-sexual bacterial population: most strains share a common ancestor ≫ *T*_mosaic_ generations ago, so their present-day genomes are completely overwritten by recombination; in large samples, some pairs of strains will share a common ancestor ≪ *T*_mosaic_ generations ago, and recombination events will be visible as blocks of local divergence against a shared clonal background. (C) Average synonymous divergence vs fraction of identical blocks for pairs of *A. putredinis* strains from unrelated hosts (S1 Text 2). Points denote individual pairs, while the marginal distribution is shown on the right; red line shows the expectation from a simple model of accumulated transfers (Eq. S5, S1 Text 2), while grey line shows the expectation when mutations are randomly distributed across the genome. (D) Spatial distribution of synonymous SNVs for three example pairs from panel C (symbols); only a portion of the core genome is shown. Points denote individual SNVs, while lines show the local divergence in sliding 300bp windows. (E) Analogous version of C for neutral simulations (S1 Text 5.3, Fig. S2). (F) Fraction of genome-wide divergence explained by the partial recombination model in different species. Bars show *R*^2^ values obtained by fitting the red line in panel C, weighted by the local density of points (S1 Text 2). Dark bars are computed using all pairs of strains, while light bars are restricted to the largest clade in species with strong population structure.

Previous work has shown that the genetic diversity within many species of gut bacteria spans a broad range of timescales [27, 45]. For example, in *Alistipes putredinis* (a prominent gut commensal) the synonymous divergence between a typical pair of strains is *d*≈2%, but some pairs of strains are separated by just a handful of SNVs (Fig. 1C). Similar pairs of closely related strains have been observed in other bacterial species, where they are often associated with clinical outbreaks or other local transmission processes [21, 36, 46]. In our case, the breadth of sampling of the human gut microbiome allows us to rule out many of these microepidemic factors, since closely related strains are frequently observed in unrelated hosts from different countries [27].

Population genetic theory predicts that similar patterns can also arise due to the local nature of bacterial recombination [28, 39]. Since the length of a typical recombined segment (ℓ_*r*_) is usually much shorter than the total genome length (*L*), there is a broad range of timescales between the first recombination event (*T*_*r*_) and the time required for the genome to be completely overwritten by imported fragments (*T*_mosaic_ ∼ *T*_*r*_ · *L*/ℓ_*r*_ ; Fig. 1B).

In quasi-sexual bacterial populations, most pairs of strains will share a common ancestor *T*_mrca_ ≫ *T*_mosaic_ generations ago, so that their present-day genomes comprise a mosaic of overlapping recombination events. However, in a large enough sample, some pairs of strains will inevitably share a common ancestor on timescales much shorter than *T*_mosaic_ (Fig. 1A). Among these “closely related” strains, recombination will not have had enough time to completely cover the ancestral genome with DNA from other, more typically diverged strains. Rather, individual recombination events will be visible as “blocks” of typical genetic divergence against a backdrop of nearly identical DNA sequence [28, 39]. These partially recombined genomes have previously been observed in other bacterial species – most notably in *E. coli* [28, 39] and other bacterial pathogens [42, 47, 48]. Fig. 1D shows that similar examples can be observed within the

*A. putredinis* population as well.

To test whether this pattern holds more broadly, we divided the core genome of each pair of strains into blocks of 1000 synonymous sites, and calculated the fraction of blocks with zero SNV differences within them. In a pair of partially recombined genomes, we would expect to see a negative correlation between the fraction of identical blocks (a proxy for the fraction of clonal ancestry) and the overall genetic divergence across the genome (Figs. 1E & S3). One can observe such a trend in *A. putredinis* (Fig. 1C) — well beyond that expected from the randomness of individual mutations. Instead, we find that a simple model of accumulated transfers (red line; S1 Text 2) can account for a large fraction of the spread in genome-wide divergence in *A. putredinis*, consistent with the partial recombination model in Fig. 1B.

Similar patterns can be observed in many other gut commensals (Figs S3 & S4). Some species exhibit some variation in the divergence of the most distantly related strains (e.g. *Bacteroides vulgatus* and *Eubacterium rectale*), consistent with the presence of sub-species or other forms of population structure [18, 27, 49]. Yet even in these cases, we find that partially recombined genomes can still account for much of the variation among more closely related strains. Across species, we find that our simple model of accumulated transfers can explain more than 50% of the weighted variation in pairwise divergence within 36 of the 43 species we examined (Fig. 1F), suggesting that it is a common phenomenon in the human gut.

Despite the generality of this trend, the total number of closely related strains can vary substantially between species (Table S2). For example, many *Alistipes* and *Bacteroides* species contain hundreds of closely related pairs, while other species like *Prevotella copri* have only a handful. While the causes of these differences are currently unclear, the simplified patterns of recombination among these strains suggest that we can use them to directly resolve individual recombination events within a range of different species.

### Measuring individual recombination events that accumulate between closely related strains in different hosts

To identify individual recombination events across a diverse range of human gut species, we turned to an automated approach for analyzing the spatial distribution of genetic differences along the core genomes of closely related pairs of strains. We chose to focus on the core genome to limit the impact of plasmids and other mobile genetic elements, which can be horizontally transmitted at much higher rates than normal chromosomal DNA [50–52]. By restricting our attention to core genes, we sought to infer the baseline rates of recombination that shape the evolution of the larger genome, which involve the permanent replacement of existing sequences in addition to successful transfers.

Our pairwise model assumes that the genetic differences along the core genome arise through a mixture of two processes: (i) point mutations (which alter individual sites) and (ii) homologous recombination events (which replace longer stretches of DNA with a corresponding fragment sampled from another strain in the populations). For sufficiently close pairs, the mutation and recombination processes have a negligible chance of overlapping, which means that they can be captured by a simple hidden Markov model (HMM) that transitions between clonal and recombined regions at different locations along the genome (Figs. 2B & S5; S1 Text 3.1). The corresponding transition rates between these states will vary between different pairs of strains, due to the differences in their time-aggregated rates of recombination. Since the genealogies of close pairs are particularly simple, these pairwise estimates can implicitly capture various forms of selection, non-equilibrium demography, and other deviations from the simplest neutral null models, even when there is insufficient data for a complete phylogenetic reconstruction.

**Fig 2.**
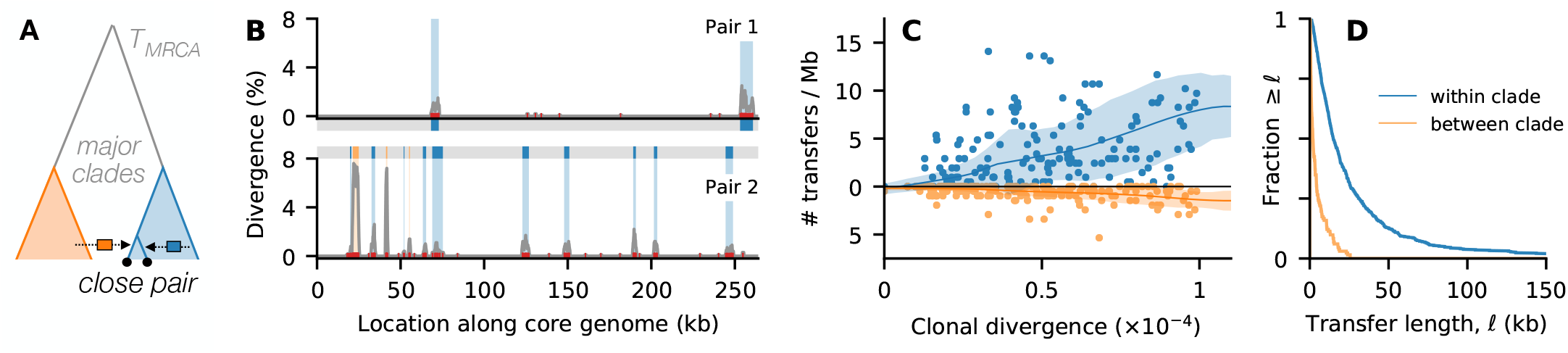
Measuring individual recombination events that accumulate between closely related strains. (A) Schematic illustration for *B. vulgatus*, which has a strong population structure with two major clades. (B) Our pairwise hidden Markov model (CP-HMM) classifies the core enome of each pair of closely related strains into clonal regions (grey) and recombined regions (blue=within-clade, orange=between-clade) based on their local synonymous divergence; points denote individual SNVs, while lines show the local divergence in sliding 1000bp windows. Data from two example pairs are shown. (C) The observed number of recombination events in all pairs of closely related *B. vulgatus* strains as a function of the synonymous divergence in their inferred clonal regions (S1 Text 3.1). These events are further partitioned into within-clade and between-clade transfers (top and bottom). Lines indicate the average trend computed using a local regression technique, while shaded regions indicate the local spread (S1 Text 3.3). (D) Distribution of the estimated transfer lengths for each of the recombination events in panel C. These data show that the rates and lengths of successful transfers strongly depend on the divergence of the imported fragments.

In contrast to previous approaches [28–31, 53], we used the empirical distribution of local divergence to model the number of SNVs imported by each recombined fragment (S1 Text 3.1). This allows us to capture the broad variation observed in different transfers (Fig. S6) in a way that is directly informed by the available data. We validated the performance of our algorithm (CP-HMM) through simulations and found that it can reliably identify individual recombination events across a range of genetic divergence scales (Figs. S7-S10; S1 Text 3.2).

Fig. 2 shows an example of this approach applied to *Bacteroides vulgatus*, one of the most abundant and prevalent species in the human gut. *B. vulgatus* was previously shown to have a strong population structure containing two major clades (Fig. 2A) [27], whose within-clade divergence is ∼10-fold smaller than the divergence between clades. We exploited this structure to further resolve the recombination events into within- and between-clade transfers based on their local sequence divergence (Fig. 2B, S1 Text 3.1). By applying our HMM algorithm to the 210 pairs of closely related *B. vulgatus* strains in our cohort, we identified a total of ≈1700 recombined regions with a mean length of ≈20kb (Figs S11 & S12; Table S3). We also applied our algorithm to a separate collection of *B. vulgatus* isolate genomes (Fig. S13; S1 Text 3.10) to verify that our conclusions were robust to the quasi-phasing approach employed in Fig. 2.

We observed an overall trend toward larger numbers of recombination events in strains with higher clonal divergence (Fig. 2C), consistent with the gradual accumulation of successful transfers over time. However, the larger sample reveals that this is not a simple linear relationship: some strains have anomalously large numbers of transfers even at low clonal divergence, while others have anomalously few transfers even at high clonal divergence (Figs. 2C). Similar results are also observed when considering the cumulative length of the recombined genome for each pair (Figs S11 & S12), which confirms that this variation is not an artifact of the event detection algorithm. Instead, these data suggest that successful transfers in *B. vulgatus* do not accumulate at a fixed recombination rate, as assumed under the simplest models of neutral evolution.

We also found that recombination between the major *B. vulgatus* clades occurred much less frequently than recombination within clades, with a ∼5-fold reduction in the total number of detected transfers as a function of their genetic divergence (Figs. 2C and S8). This genetic isolation could arise from several factors, ranging from reduced opportunities for recombination (e.g. due to ecological isolation [2] or fewer homologous flanking regions for initiating strand invasion [54, 55]) to greater downstream incompatibilities in the acquired fragments (e.g. epistatic interactions [56, 57] or mismatch-repair-mediated proofreading [58, 59]). In this case, the larger ensemble of detected transfers allows us to further distinguish between these scenarios. Beyond the reduction in the number of detected recombination events, we also observed a systematic difference in the lengths of the individual transfers, with a ∼7-fold reduction in the median transfer length between clades (Figs. 2D and S8). These differences indicate that the greater genetic isolation of the *B. vulgatus* clades cannot be captured by a simple rescaling of the recombination rate, and that additional factors like epistasis or mismatch-repair-mediated proofreading are necessary to explain the data.

### Variation of recombination rates within and across gut species

To understand how these results for *B. vulgatus* extend to other members of the gut microbiome, we applied the same approach to the other species in our dataset with a sufficient number of closely related strains. This pairwise analysis yielded a total of 228,078 recombined regions in 7,383 closely related pairs from 29 different species. These data revealed systematic variations in the rates and lengths of transferred fragments across many prevalent gut species (Figs 3, S14-S16), similar to *E. coli* and other bacterial pathogens [16, 60–62].

**Fig 3.**
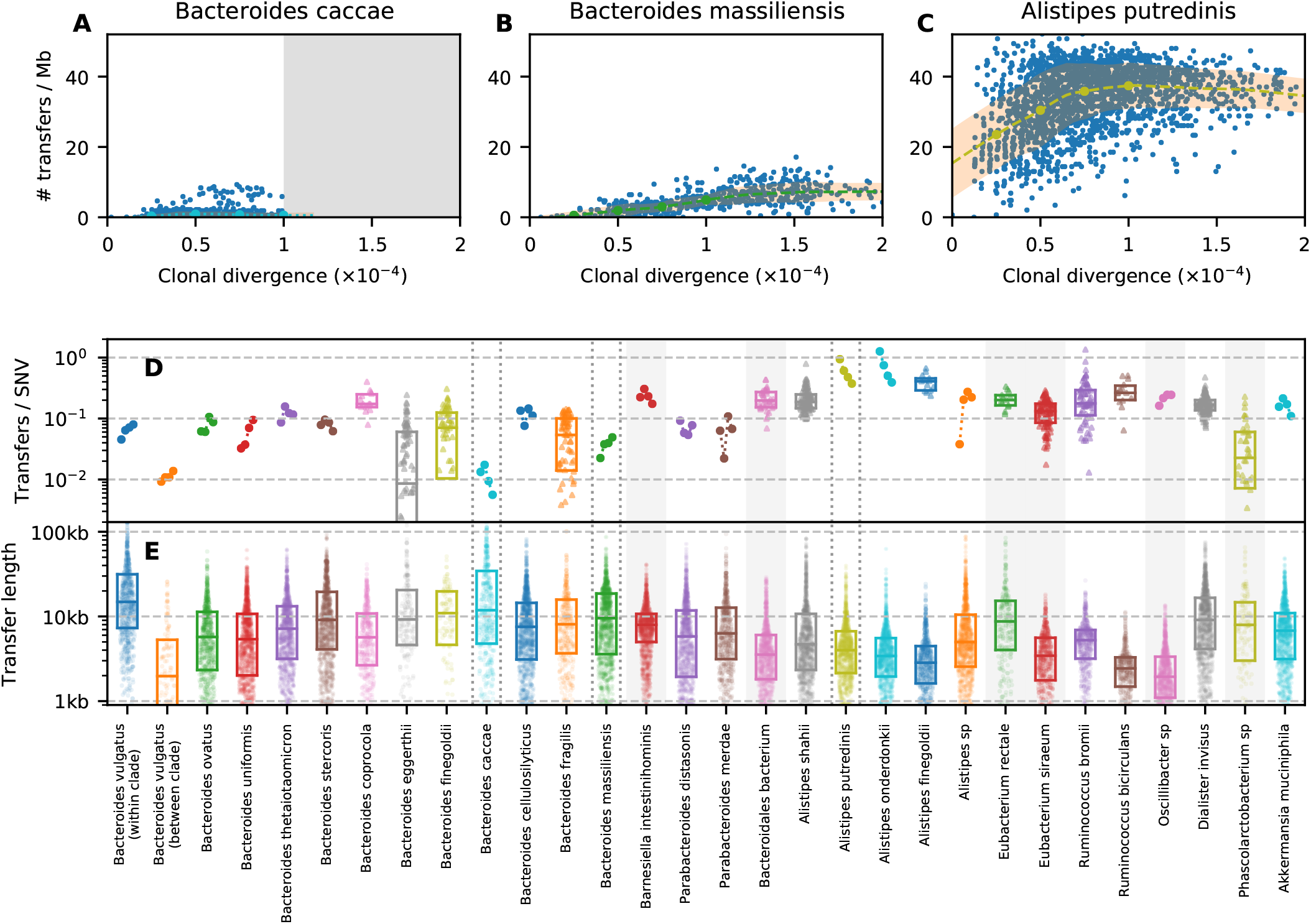
Heterogeneous recombination rates within and between prevalent gut species. (A-C) Analogous versions of Fig. 2C for three example species, which were chosen to illustrate a range of characteristic behaviors. Grey regions denote the points that were excluded by our filtering steps (S1 Text 3.3). (D) Apparent recombination rates (number of transfers / clonal divergence / core genome length) for all species with a sufficient number of closely related strains (S1 Text 3.3). For species with >100 close pairs, we plot the average recombination rate at four characteristic divergence times (*d*_*c*_ = 2.5, 5, 7.5, 10 × 10^−5^, highlighted as points along the trend lines in panels A-C) using the trend lines in panels A-C; estimates are connected by lines to aid visualization. For species with <100 close pairs, we plot the distribution of apparent recombination rates for all individual pairs; box plots indicate the median and inter-quartile range. (E) Lengths of recombined fragments for each of the species in panel D. Symbols show the lengths of all detected transfer events across all pairs of closely related strains; box plots indicate the median and inter-quartile range.

We found that some of these trends were consistent with the phylogenetic relationships between species. For example, species in the *Alistipes* genus tended to have relatively frequent and short transfers, while *Bacteroides* species tended to have lower rates and longer transfers. However, we also observed large differences within individual genera. For instance, *Bacteroides massiliensis* has a relatively linear accumulation of transfers over time (Fig. 3B), while most pairs of *Bacteroides caccae* strains have few detected recombination events (Fig. 3A). The typical transfer length varies among *Bacteroides* species as well (6-35kb), spanning a larger range than *Alistipes* (3-6kb).

Zooming in further, we also observed considerable variation within individual species. Some of these differences could be attributed to the presence of strong population structure (similar to *B. vulgatus*), with a reduction in both the rates and lengths of successful transfers between highly diverged clades (e.g. *Alistipes Shahii*; Fig. S17). However, we also observed substantial variation even in the absence of population structure. For example, *A. putredinis* contains many closely related strains with an anomalously large number of transfers, as well as an excess of more diverged strains with few recombined segments (Figs. 3C & S9). Other species (e.g. *B. caccae*; Fig. 3A) exhibited bimodal distributions of transferred fragments. None of these behaviors can be captured by a single underlying recombination rate.

Interestingly, apart from the handful of species with strong population structure, we observed no systematic trend between the frequency of recombination and the divergence of the transferred fragments (Figs 4 & S18), as expected under certain models of homologous recombination [19, 63]. This observation, in combination with the large number of species in our dataset, helps shed further light on the mechanisms that could be responsible for the lower recombination rates we observe between clades. For example, *B. thetaiotaomicron* and *B. stercoris* both maintain high recombination rates at synonymous divergences comparable to the genetically isolated clades observed in *B. vulgatus* and *B. finegoldii* (Figs 4 & S18). This suggests that the genetic isolation of these clades is not a product of their underlying recombination machinery (which should be similar in different *Bacteroides* species) but rather by genetic incompatibilities that have accumulated between the two clades, or related scenarios like incompatible restriction-modification systems [64–67]. Understanding the ecological and evolutionary forces that caused these incompatibilities to emerge within some *Bacteroides* species but not others is an interesting avenue for future work.

**Fig 4.**
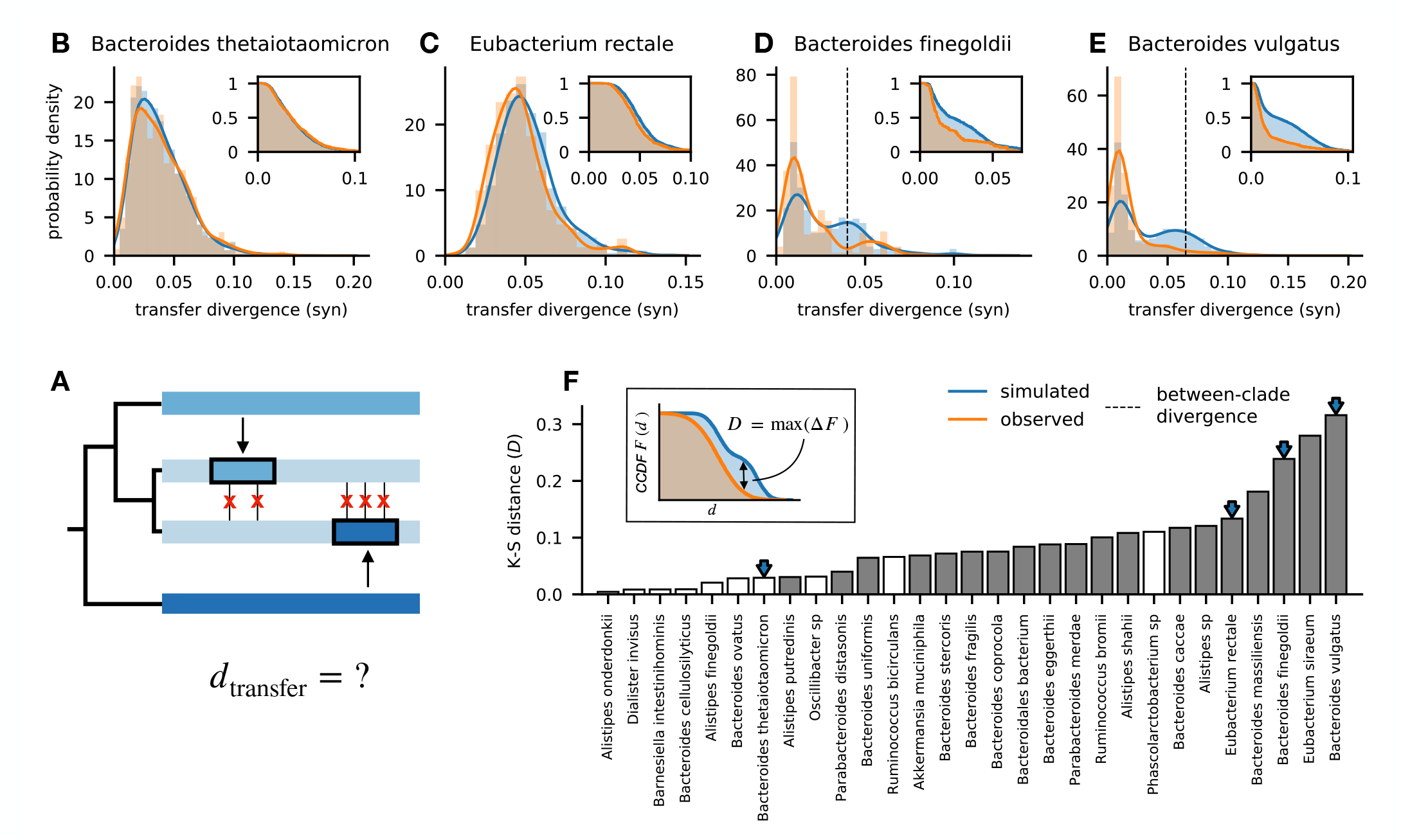
Quantifying the frequency of recombination as a function of the genetic divergence between donor and recipient DNA sequences. (A) Schematic illustration showing the genetic divergence of two recombined fragments relative to the focal pair of genomes. The synonymous divergence of each detected transfer is computed and aggregated across all closely related pairs within a species. (B-E) Distribution of donor-recipient divergence for all detected transfers in four example species. Orange lines show the observed data, while the blue lines show a null expectation obtained by randomly drawing segments from the observed collection of genomes (S1 Text 3.6). Insets show the corresponding complementary cumulative distribution functions (CCDF). For species with a strong clade structure (D,E), the average between-clade divergence is indicated by dashed vertical lines. (F) Differences between the observed and simulated divergence distributions for all of the species in Fig. 3, summarized by the Kolmogorov–Smirnov (K-S) distance (inset). Solid bars indicate statistically significant differences (*P* < 10^−3^; one-sided K-S test). Arrows indicate the example species in panels B-E. These data show that many species exhibit only small differences between their observed and expected divergence distributions (K-S distance ≲0.1), even when their overall divergence is comparable to counterexamples like D & E.

### Signatures of within-host recombination in co-colonized hosts

Our preceding analysis focused on the successful transfers that have accumulated between closely related strains in unrelated hosts. How do these long-term dynamics – which aggregate over multiple host colonization cycles – emerge from the local processes of competition and colonization within individual hosts?

Some of this recombination could occur when multiple strains of the same species are present within the same host [68]. While examples of co-colonization are less common in the human gut [27, 45], we can still identify many individual hosts in our larger cohort in which two diverged strains were present at intermediate frequencies, based on the frequencies of SNVs within their corresponding metagenomes (Fig. S1). Recombination between these strains will generate hybrid genomes that contain a short fragment from their donor (Fig. 5A). Each of these hybrid strains will originate as a single cell, and will not be visible in a mixed sample unless they later rise to appreciable frequencies. Such a shift could occur through a single-cell bottleneck, e.g. if the hybrid strain is lucky enough to found a new population in naive host. Alternatively, if the transferred fragment provides a fitness benefit to the recipient strain, it can rapidly increase in frequency within its host and eventually displace its parent. These “gene-specific sweeps” will lead to a characteristic depletion of SNVs within the donated region in a mixed population sample, while preserving the remaining genetic variation elsewhere along the genome (Fig. 5A). The higher frequencies of the resulting hybrids will make them substantially more likely to seed future colonization events in other hosts, suggesting that they could play an important role in generating the recombination events we observed in Figs. 2 & 3.

**Fig 5.**
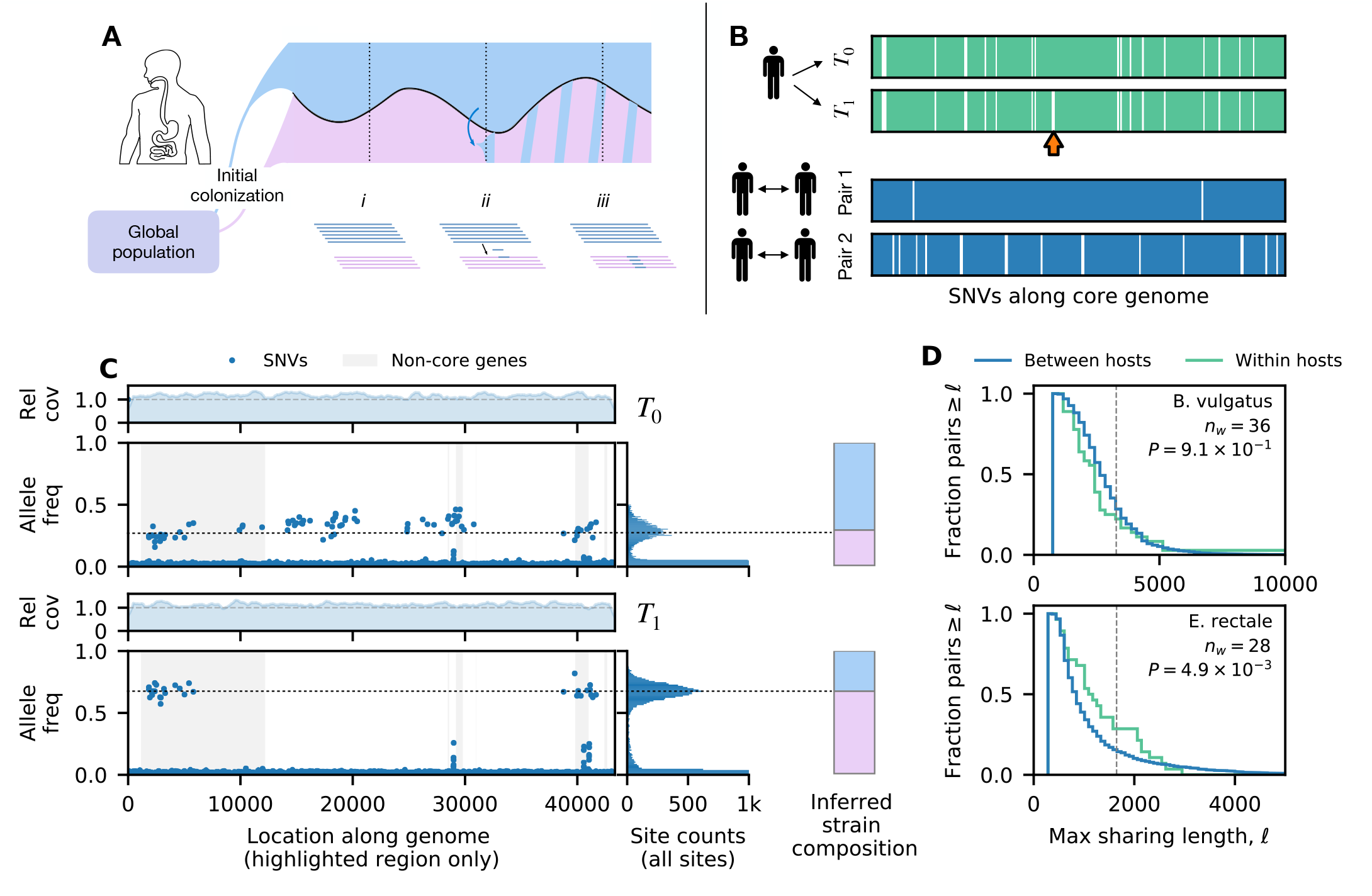
Signatures of within-host recombination among co-colonizing strains. (A) Schematic illustration of a potential recombination scenario: (i) a single host is colonized by a pair of diverged strains; (ii) recombination generates hybrid strains that initially reside at low frequencies; (iii) if a hybrid replaces its parent (e.g. due to a selective sweep), it will lead to a depletion of genetic diversity within the transferred region. (B) An example hybrid sweep in a *B. vulgatus* population. Top two panels show the spatial distribution of within-host SNVs (green vertical lines) from a single host sampled at two timepoints (Δ*t*∼6mo); orange arrow highlights a sudden depletion of SNVs within a ∼ 20kb region. For comparison, the bottom two panels show analogous distributions computed for pairs of strains from different hosts. (C) Detailed view of the putative recombination event in panel B. Top and bottom parts show metagenomic data collected from the same host at timepoints *T*_0_ (top) and *T*_1_ (bottom). In the bottom panel of each timepoint, symbols denote the frequencies of within-host SNVs in the highlighted region of panel B (arrow), which are polarized according to the major allele at *T*_0_; for comparison, the genome-wide distribution of SNV frequencies is shown on the right, illustrating the coexistence between two dominant strains at both timepoints (black dotted lines, bar plots). In the top panel of each timepoint, solid lines denote the local relative coverage, estimated from a moving average of the local read depth divided by the median read depth across the genome. The consistent relative coverage around 1 (grey dashed lines) at both timepoints indicates that the depletion of SNVs in the highlighted regions is not caused by large deletion in one of the coexisting strains. (D) Distribution of the longest sharing tract in each co-colonized host for two example species (S1 Text 4.3). Grey dashed lines indicate the mean transfer length inferred in Fig. 3E. The total number of co-colonized samples and the *P*-value under the one-sided Kolmogorov-Smirnov test are shown. The *B. vulgatus* distribution is indistinguishable from its between-host counterpart, while *E. rectale* possesses a significantly higher rate of within-host sharing (*P* < 5 × 10^−3^).

Figure 5 shows an example of this scenario in a longitudinally sampled host who was co-colonized by a pair of typically diverged *B. vulgatus* strains (*d*≈1%). We observed a sudden depletion of within-host SNVs within a ∼20kb region during the ∼6-month interval between samples (Fig. 5B,C), while the SNV patterns across the rest of the genome were largely preserved. This local depletion of diversity cannot be explained by a large deletion event in one of the two strains, since the estimated copy number of the recombined region remained close to one at both timepoints (Figs. 5C & S19). This region spanned a total of 25 core and accessory genes on the reference genome, including a resistance-nodulation-division (RND) family efflux pump (Table S5 and Fig. S25); at present, it is not clear which of these genes was responsible for driving the sweep, or if the recombined fragment was simply hitchhiking alongside a different causative mutation.

With limited longitudinal data from co-colonized hosts, it is difficult to find many contemporaneous examples like the one illustrated above. However, we reasoned that the remnants of these gene-specific sweeps would still be visible even in metagenomic data from a single timepoint. Previous work suggests that conspecific strains can coexist within their hosts for years at a time [27, 69, 70]. Any gene-specific sweeps that occur during this interval will produce an extended run of zero SNVs against the backdrop of an otherwise diverse metagenome. We identified many such runs of shared ancestry among the co-colonized hosts in our cohort (S1 Text 4.2), including several other examples in the *B. vulgatus* population above (Figs. 5B). These runs can extend for thousands of base pairs, and are significantly longer than we would expect if the mutations were randomly scattered across the genome (*P* < 10^−10^, Fig. S20). This suggests that they could be candidates for previous gene-specific sweeps that occurred within the host’s lifetime.

However, it is important to distinguish this scenario from older recombination events that were inherited by the strains before they colonized their current host (Fig. 5A). Estimates suggest that a 10kb fragment will require hundreds of years on average to accumulate its first mutation [71], which implies that any given run could be consistent with a broad range of possible ages. Consistent with this expectation, we also observed many long runs of shared ancestry when comparing strains from unrelated hosts – some of which extended for as long as the within-host examples above (Figs. 5B & S20).

This suggests that the true signal of within-host recombination must be distinguished from this baseline level of sharing. We reasoned that if within-host recombination was prevalent, we should still expect to see longer runs of shared ancestry in co-colonizing strains compared to random pairs of strains obtained from unrelated hosts. To test this idea, we used the length of the longest run as a test statistic, and asked how the distribution of this quantity differed between co-colonizing strains of the same species and random pairs of strains selected from unrelated hosts.

We observed a strong enrichment of long runs in co-colonizing strains of *E. rectale* (Fig. 5D), which suggests that they were likely caused by previous within-host recombination events similar to the *B. vulgatus* example above. Similar results were obtained when we examined the total length of runs that exceeded a given length threshold (Fig. S21). In contrast, we found that some of the other species with high rates of recombination across hosts (e.g. *A. putredinis*; Fig. 3C) did not show any enrichment in within-host sharing (Fig. S22). This negative result could imply that co-colonizing strains recombine less frequently in these species, or that fewer hybrid strains manage to sweep to high frequencies. It could also occur if the background levels of between-host sharing are sufficiently frequent that they overwhelm any signature of within-host sweeps. This scenario could be particularly relevant for species like *B. vulgatus* (Fig. 5), in which nearly half of all random strain pairs share identical sequences longer than the typical transfer length in Fig. 2. These results show how understanding the population genetic patterns between hosts can be important for resolving the evolutionary forces within individual host communities.

### Distribution of shared DNA segments across hosts reveals selection on recent transfers

The high levels of between-host sharing in species like *B. vulgatus* raise a natural question: why do random pairs of strains share so many stretches of identical DNA within their core genomes? Population genetic theory predicts that such tracts of shared ancestry can emerge even in simple neutral scenarios due to the joint action of recombination, mutation, and genetic drift [72]. For a random pair of strains, the ex<u>p</u>ected number of shared fragments longer than ℓ scales as ∼*L*/*d*ℓ^2^(1 + *r*/*μ*)^2^, where *d* is the average divergence between typical pairs of strains (Fig. S23; S1 Text 5.1). The slow decay with ℓ and *r* implies that this number will often be larger than one, even for tracts as long as ℓ∼10kb. This suggests that the presence of shared segments alone is not surprising.

However, this simple neutral scenario makes strong predictions about how often a given region is shared across multiple pairs of strains. To test whether this scenario could recapitulate our data, we scanned across the genome of each species, an<u>d</u> calculated the probability that each position was involved in a long shared segment (ℓ · *d* > 15, Fig. 6A, S1 Text 5.2). This analysis revealed a systematic variation in the probability of shared segments at different genomic locations (Fig. 6B-D).

**Fig 6.**
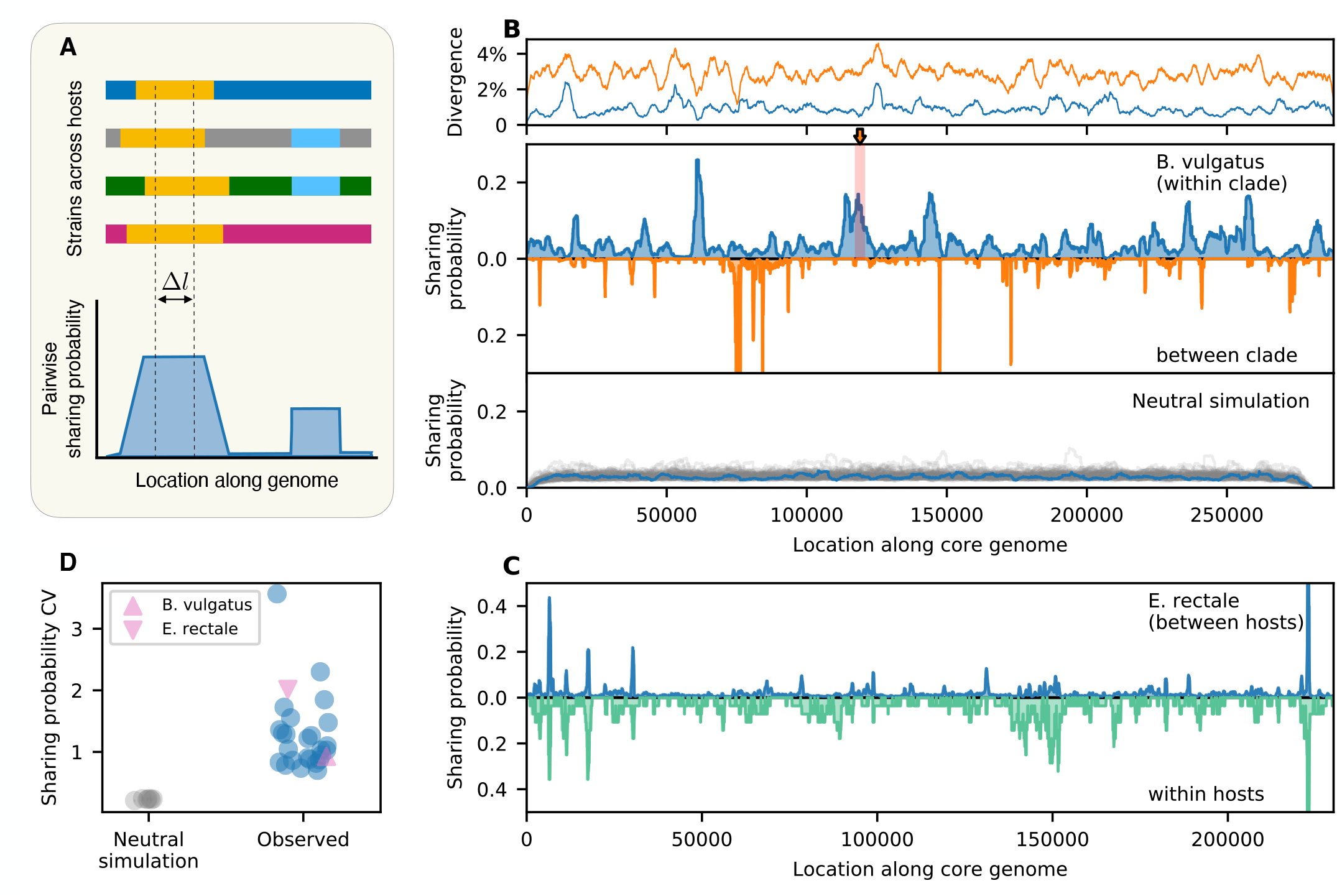
Global distribution of shared DNA segments reveals selection on recent transfers. (A) Schematic of pairwise haplotype sharing metric: for each position in the core ge<u>no</u>me, we compute the fraction of strain pairs from different hosts that have identical genotypes across a window of Δℓ ≫ 1/*d* synonymous sites (S1 Text 5.2). (B) Observed sharing landscape for *B. vulgatus* (middle panel); separate comparisons are performed for strains from the same clade (blue, Δℓ ≈ 1500 synonymous sites ≈ 10kb) or different clades (orange, Δℓ ≈ 220 synonymous sites ≈ 1.5kb). The top panel shows the average synonymous divergence computed in sliding windows of size Δ*l* = 3000. These landscapes reveal regions of elevated sharing across hosts (e.g. shaded region) that cannot be explained by local reductions in diversity. Red shaded region indicates the within-host recombination event in Fig. 5B-C. The bottom panel shows analogous sharing landscapes from neutral simulations (S1 Text 5.3), which display more even rates of sharing across the genome. Grey lines denote 100 simulation runs with the same parameters, while the blue line highlights one typical run. (C) Sharing landscape for *E. rectale*, computed for pairs of strains in different hosts (top) and co-colonizing strains from the same host (bottom). (D) Heterogeneous sharing landscapes across 27 species. Blue points show the coefficient of variation of the sharing probability across the genome for all species with sufficient between-host comparisons. *B. vulgatus* (within clade) and *E. rectale* are highlighted as pink triangles. Grey points show analogous values derived from neutral simulations across a range of parameter values (S1 Text 5.3); each point denotes the mean of 100 simulation runs, while lines show the standard deviation.

An example of this behavior is shown for *B. vulgatus* in Fig. 6B. A typical site in the *B. vulgatus* genome has a 3% chance of being shared in a segment longer than ∼10kb. However, we observed that many local regions were shared much more frequently than the genome-wide average, despite having comparable levels of genetic diversity (Figs. 6B & S24). Some of these peaks are driven by the expansion of a single dominant haplotype, while others correspond to multiple distinct haplotypes that are shared by different sets of strains (Figs. S25-S27). Similar “sharing hotspots” can be observed in other prevalent gut species as well (Fig. 6C,D).

This high degree of heterogeneity is inconsistent with simple neutral models of bacterial evolution. Simulations show that neutral models generate significantly tighter correlations between the average and maximum levels of sharing across the genome (*P* < 10^−8^; Student’s t-test; Figs. 6B,D & S28). We also asked whether this heterogeneity could be explained by varying recombination rates along the genome [51, 73, 74].

However, our simulations showed that the sharing hot spots in Fig. 6 are qualitatively distinct from traditional recombination hot spots. Local increases in the recombination rate actually decreased the probability of sharing longer segments (Fig. S29), since recombination tends to produce larger numbers of haplotypes with different combinations of mutations. Consistent with this finding, we observe few systematic correlations between the haplotype sharing landscapes in Fig. 6 and the recombination hot spots inferred from Fig. 3 (Fig. S30).

These analyses suggest that the heterogeneous sharing probabilities in Fig. 6B are likely driven by positive selection on fragments that are spreading through the population via recombination. Consistent with this hypothesis, we found that the regions with the highest levels of sharing are statistically enriched for certain functional genes (e.g. glycosyltransferases) that have previously been shown to be under selection in the gut [71] (S1 Text 5.5). We also found that the sharing landscape qualitatively differs for fragments that are shared within vs between the major *B. vulgatus* clades (Fig. 6B). This provides further evidence that the selection pressures are specific to the identities of the donated and recipient DNA sequences.

Finally, we asked how these global selection pressures were related to the within-host sweeps we detected in Fig. 5. For example, we found that the within-host sweep event in Fig. 5C occurred within one of the most prominent sharing hotspots in *B. vulgatus* (Fig. 6B), which is peaked around three RND efflux pump genes (Fig. S25). This suggests that both events were likely driven by a common set of selection pressures. However, this parallelism did not arise through selection of the same DNA sequences: while the sweeping haplotype in Fig. 5C was also present in a few other hosts in our panel, we found that several other distinct haplotypes contributed to the global sharing hotspot at this location (Fig. S26). This suggests that natural selection has promoted the transfer of multiple genetic variants at these loci – similar to a soft selective sweep [75]. Even larger differences were observed within the *E. rectale* populations in Fig. 5E. In this case, while we observed some overlap in the sharing hotspots within vs between hosts, we also identified several new hotspots that were only present among co-colonizing strains (Fig. 6C, Fig. S31). These significant differences in the locations of the within-host sharing events (*P* < 0.001, permutation test; S1 Text 5.4) provide further evidence that they were likely driven by selection on recent transfers within their hosts. More broadly, these results show that within-host sweeps are not always local versions of ongoing global sweeps, but may reflect distinct and repeatable selection pressures that are specific to the within-host environment (e.g. competition- vs colonization-related traits [76]). Understanding the tradeoffs that give rise to these different selection pressures is an interesting topic for future work.

## Discussion

Recombination is a ubiquitous force in bacterial evolution, but dynamics of this process are still poorly understood in many natural microbial populations. Here we sought to quantify these dynamics by leveraging the broad range of timescales inherent in the human gut microbiome ecosystem. By analyzing recent recombination events within a panel of 29 gut commensals, we were able to identify general trends across diverse bacterial species that inhabit a common host-associated environment.

The overall rates of recombination we observed across hosts are comparable to other bacterial species [25, 33, 47], and are consistent with the strong decay in linkage disequilibrium observed in global samples of gut bacteria [27, 77]. Across species, we found that recombination is responsible for introducing >10x as much variation as mutation (*T*_mrca_/*T*_mosaic_ ≳ 10; Fig. S16), which implies that the genomes of typical circulating strains are almost completely overwritten by recombination. These values are broadly consistent with previous observations in bacterial pathogens, though their different sampling strategies can make it difficult to perform detailed numerical comparisons (Fig. S32; S1 Text 3.10.1). The observation of such high rates of genetic exchange in commensal gut bacteria poses challenges for efforts to identify signals of parallel evolution in strains sampled from different hosts [71], or signals of codiversification across host populations [78, 79], since they imply that individual variants can frequently decouple from the genome-wide phylogeny. In this case, more elaborate methods like the haplotype sharing metric in Fig. 6 could be useful for resolving common selection pressures across hosts.

Although the long-term recombination rates in Fig. 3 represent an average over multiple host colonization cycles, it is useful to consider their implications when extrapolated down to the scale of a single host community. If we assume the recombination events in Fig. 3 accumulate largely neutrally (or via neutral hitchhiking [80]), then the rates implied by these data suggest that every site in the genome will be involved in more than a thousand recombination events within a single day (S1 Text 3.8). These ballpark estimates suggest that there will be numerous opportunities for adaptive mutations to spread between co-colonizing strains within a host (e.g. during a fecal microbiota transplant), even if the donor or recipient strain is present at a low frequency (e.g. ∼0.1%). However, since each recombination event originates in a single cell, it can still take tens of thousands of generations (∼5-50 years) before a typical ancestral lineage will be involved in a single *de novo* recombination event. The large gap between these timescales can help explain why recombination can be an important driver of adaptation in the gut (Fig. 5) [27], while also preserving the largely clonal structure observed in individual host populations [27, 45, 69, 81]. We emphasize that these extrapolations should be treated with a degree of caution, since they assume that most of the recombination events in Fig. 3 are effectively neutral. If the vast majority of these events were locally adaptive, then the true rate of recombination could be smaller than the apparent rates in Fig. 3 (S1 Text 3.8).

In addition to the overall rates, the enhanced resolution of our approach also provided new insights into the dynamics of recombination within the gut microbiota. Extending previous findings in other bacterial species [28, 60, 82–84] (see [16] for a review), we observed widespread strain-level variation in recombination rates within many commensal gut species — at least some of which could be attributed to existing population structure (e.g. “sub-species” [49] or “ecotypes” [85]). In these handful of examples, the comparative nature of our dataset helps illuminate the potential causes of this genetic isolation. By comparing the rates and lengths of successful transfers in species with different levels of genetic diversity, we obtained new evidence that the barriers to recombination are likely driven by negative selection on the recombined fragments (e.g. due to genetic incompatibilities), or related scenarios like incompatible restriction-modification systems [64–67], rather than passive mechanisms like ecological isolation or homology-dependent recombination rates Fig. 4. Our results suggest that understanding the causes and extent of these incompatibilities will be important for predicting the genetic cohesion and structure of bacterial species.

While our underlying approach relied on the presence of closely related strains to resolve individual recombination events, the widespread occurrence of these partially recombined genomes is still an interesting evolutionary puzzle. We previously showed [27] that the ecological structure of the human gut microbiome allows us to rule out common sampling biases (e.g. microepidemics or clonal blooms) that have been conjectured to play a role in other microbial species [21, 36, 46]. We also observed considerable variation across different commensal gut bacteria, with more than a quarter of the species in our panel containing just a handful of closely related strains from unrelated hosts. How could the same sample of hosts generate such a broad range of closely related strains in different species? The simplest neutral models predict a characteristic relationship between the mosaic timescale (*T*_mosaic_/*T*_mrca_) and the fraction of partially recombined genome pairs in the sample (Fig. S33B)[42]. However, we found that the observed fractions are often much higher than this baseline expectation, and show little correlation with the estimated recombination rates (Fig. S33A). This suggests that new evolutionary models will be necessary to understand this puzzling feature of many natural bacterial populations.

Our results suggest that at least some of the long-term recombination dynamics across hosts arise from within-host sweeps of transferred fragments in hosts with multiple co-colonizing strains. This could provide a potential mechanism for the strain-level variation in recombination rates we observed in many species, since both the colonization structure and propensity for sweeps can vary dramatically in different hosts [27, 69, 81, 86]. It remains unclear whether non-sweeping transfers could also play an important role in generating the long-term rates of recombination across hosts. Our results highlight the challenges involved in detecting these events, since we found that even unrelated strains can frequently share long stretches of DNA that are likely spreading through the global population via natural selection. These scenarios could potentially be distinguished with denser longitudinal sampling or larger samples of clonal isolates (e.g. using single-cell techniques [87]), which would allow us to distinguish between pre-existing and *in situ* transfers [68].

While our present data do not provide direct information about the underlying mechanisms of horizontal DNA exchange in these species, our findings impose some interesting constraints on the potential mechanisms that might be involved. Many of the species in our panel (e.g. *Bacteroides*) are not known to be naturally competent [88], but still have long-term recombination rates that are as high as other species that are (e.g. *Streptococcus pneumoniae* [47, 89]). Many gut commensals are known to engage in conjugative transfer, both *in vitro* and *in vivo* [90]. However, the time required for bacterial conjugation carries a substantial opportunity cost in the high growth regimes of the large intestine, and would need to be ameliorated by a corresponding fitness benefit or residence in a privileged spatial location [91]. Moreover, we observe little correlation between the overall rates of recombination in different species and their frequency of apparent multi-colonization (Fig. S34). This suggests that these and other mechanisms that require physical proximity between strains are not the major driver of the long-term recombination rates we observed across hosts. It is possible that other species (e.g. phage or another commensal bacterium in the larger gut community) could serve as intermediate vectors for horizontal transfer between strains that are physically segregated in different hosts. Such inter-species transfer events have recently been observed within individual gut microbiomes [3, 11, 14]. It remains to be seen whether the rates of this process are sufficient to generate the long-term recombination rates we observe within species.

An important limitation of our metagenomic approach is that it is primarily restricted to recombination events within the core genome. While this provides important information about the long-term rates of recombination within gut commensal species, it is possible that much of this core-genome hybridization could be driven by positive selection on linked accessory genes (e.g. antibiotic resistance genes). Future applications of our methods on growing collections of clonal isolates [92] could shed light on these functional targets of horizontal transfer [93], and thereby provide a fuller picture of the landscape of bacterial recombination within the gut microbiota.

## Acknowledgments

We thank S. Maslov and G. Birzu for useful discussions, and D. Wong, S. Walton, and J. Ferrare for comments and feedback on the manuscript. This work was supported in part by a Stanford Bio-X Bowes Fellowship (to Z.L.), the Alfred P. Sloan Foundation grant FG-2021-15708, NIH NIGMS Grant No. R35GM146949, and a Terman Fellowship from Stanford University. B.H.G. is a Chan Zuckerberg Biohub – San Francisco Investigator.

## Data and Code Availability

The raw sequencing reads for the metagenomic samples used in this study were downloaded from public repositories listed in Refs. [94–97]. All necessary metadata, as well as the source code for the sequencing pipeline, downstream analyses, and figure generation are available at GitHub (https://github.com/zhiru-liu/microbiome_evolution).

---

**Fig S1.**
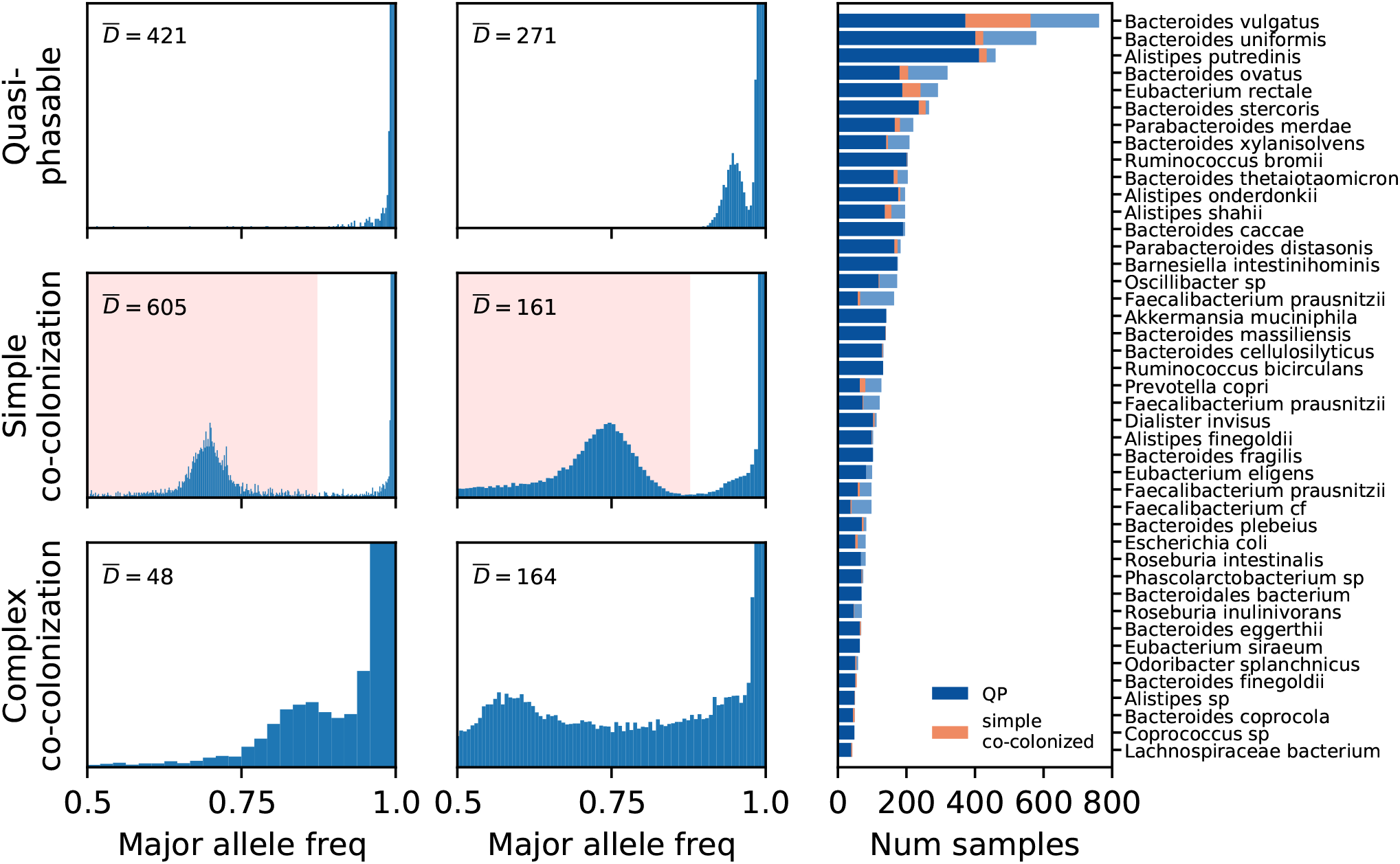
Inferring host colonization structure from the within-host site frequency spectrum. Left: Example site frequency spectra of 6 *B. vulgauts* samples from 6 unrelated hosts, showing only synonymous sites in the core genome. Quasi-phaseable (QP) samples (top) are dominated by a single strain of a given species [27]. Simple co-colonized samples (middle) are colonized by two major strains at intermediate frequencies. In this case, genetic differences between the two major strains can be reliably inferred (sites highlighted in pink regions; S1 Text 4.1). More complicated examples that do not fall into these two categories (bottom) were discarded from further analysis. In each of the panels, the vertical axis is scaled by an arbitrary constant and truncated to emphasize the peak at intermediate frequencies; the median read depth (*D*) is also listed for reference. Right: The distribution of QP samples and simple co-colonized samples among the 43 species studied. Species are sorted by the total number of samples.

**Fig S2.**
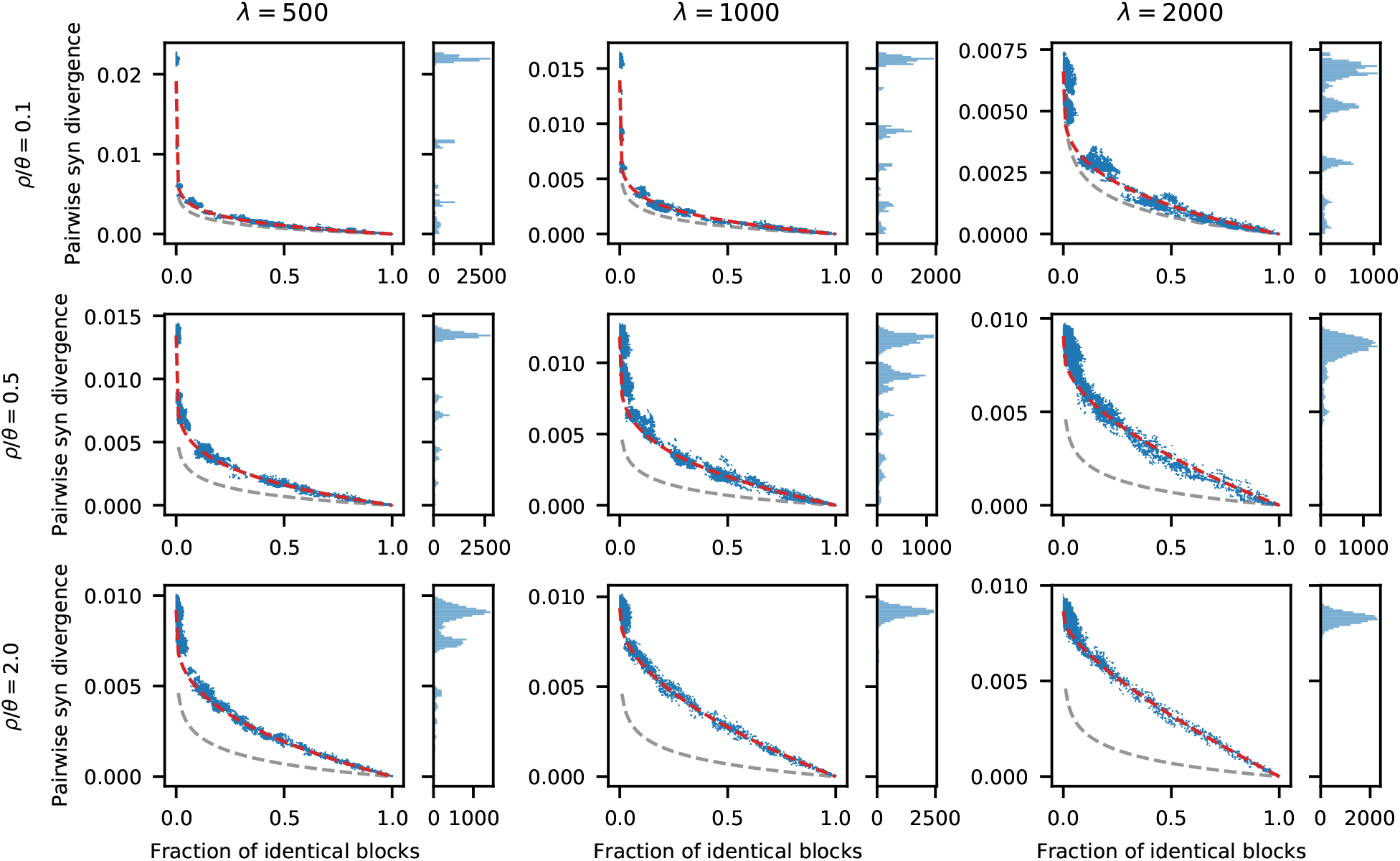
Joint distributions of pairwise diversity statistics in a simple neutral model. Analogous versions of Fig. 1C, E computed for simulated data from FastSimBac (S1 Text 5.3). Three sets of recombination lengths (*λ*) and scaled recombination rates (*ρ*) are shown. The partial recombination model (red lines) provides a good fit to the joint distribution of pairwise statistics in all parameter combinations, while larger values of *λρ*/*θ* (i.e. stronger recombination) leads to larger deviation from the random expectation (grey lines).

**Fig S3.**
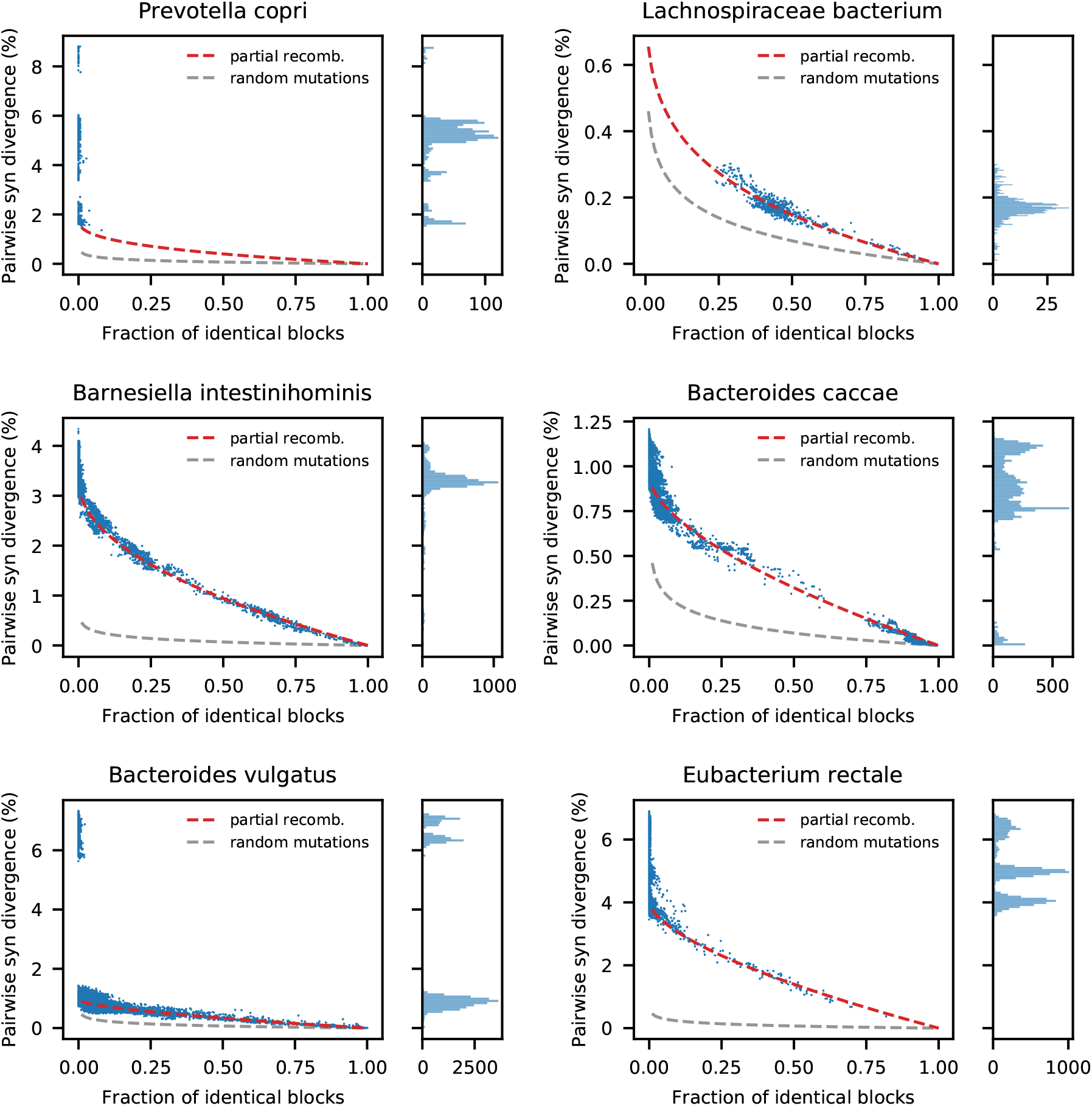
Joint distributions of pairwise diversity statistics for six example species. Analogous versions of Fig. 1B for six additional example species, which were chosen to illustrate a range of characteristic behaviors. Top panels show example species that have different distributions of partially recombined genomes: *Prevotella copri* lacks closely related pairs almost completely, while all pairs in *Lachnospiraceae bacterium* have identical blocks covering >25% of the genome. In this case, the joint distribution reveals distinct behaviors that are difficult to infer based on the genome-wide divergence distribution alone. Middle panels show example species with typical numbers of closely related pairs and minimal population structure: *Barnesiella intestinihominis* has a tightly clustered distribution, akin to *Alistipes putredinis* in Fig. 1B, while *Bacteroides caccae* possesses uneven distribution with a significant fraction of extremely close pairs. Bottom panels show example species with different degrees of population structure: *Bacterides vulgatus* features two major clades with 7% synonymous divergence, while *Eubacterium rectale* shows a less clearly separable population structure.

**Fig S4.**
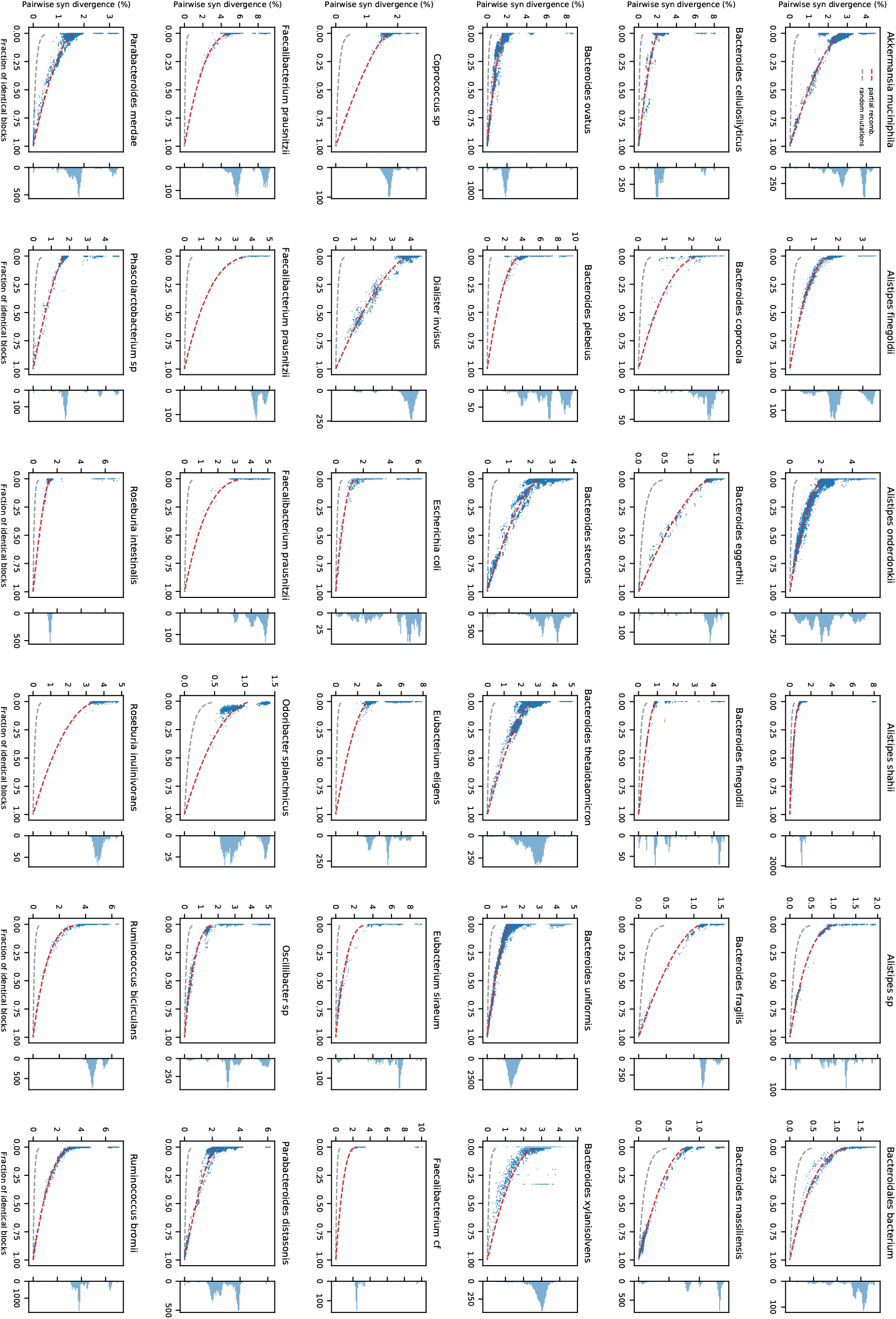
Joint distributions of pairwise diversity statistics for all remaining species. Analogous versions of Figs. 1C & S3 for all species not previously shown in these figures.

**Fig S5.**
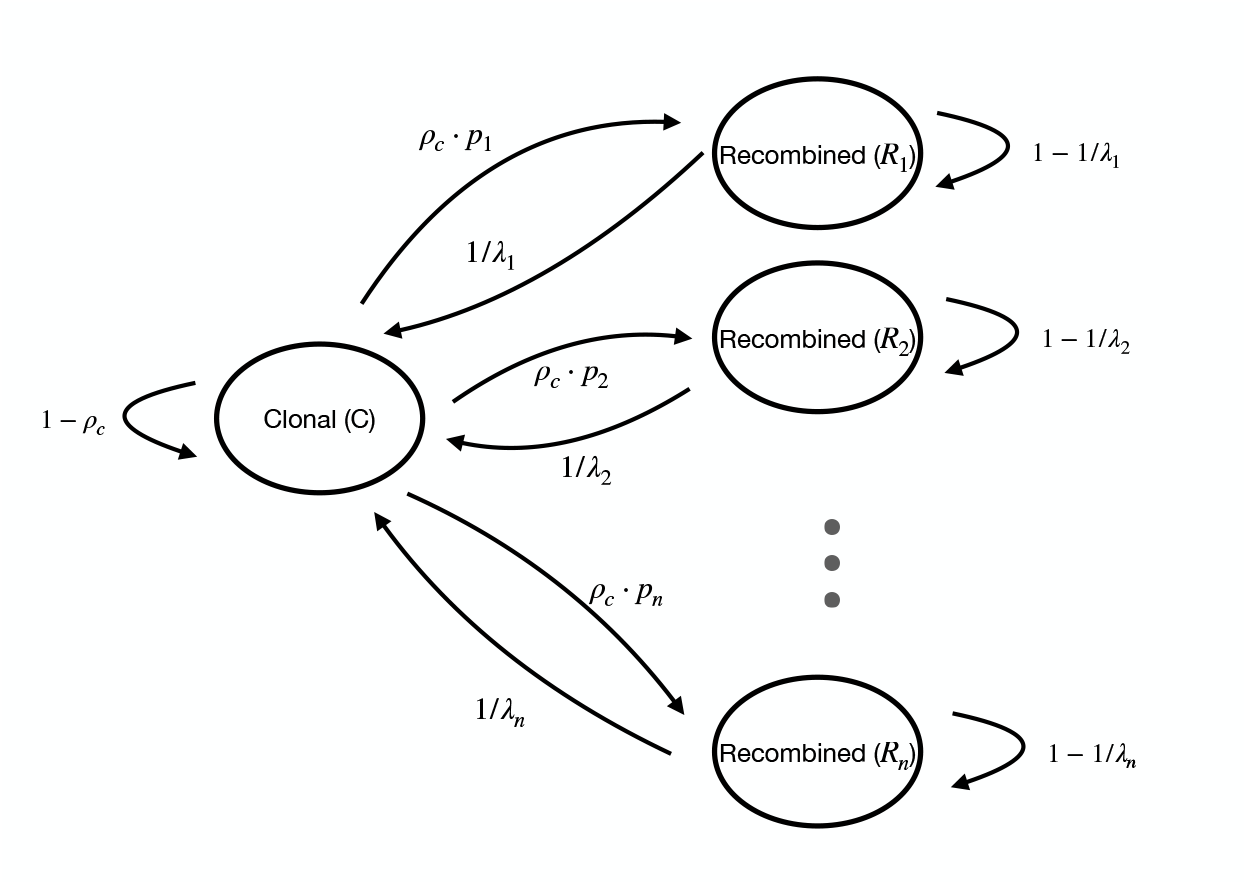
Schematic of the hidden states and transition rates in the close-pair HMM. The allowed transitions are sparse, in that each recombined state is only connected to the clonal state. This simplified structure directly follows from the assumption that recombination events rarely overlap for a sufficiently close pair.

**Fig S6.**
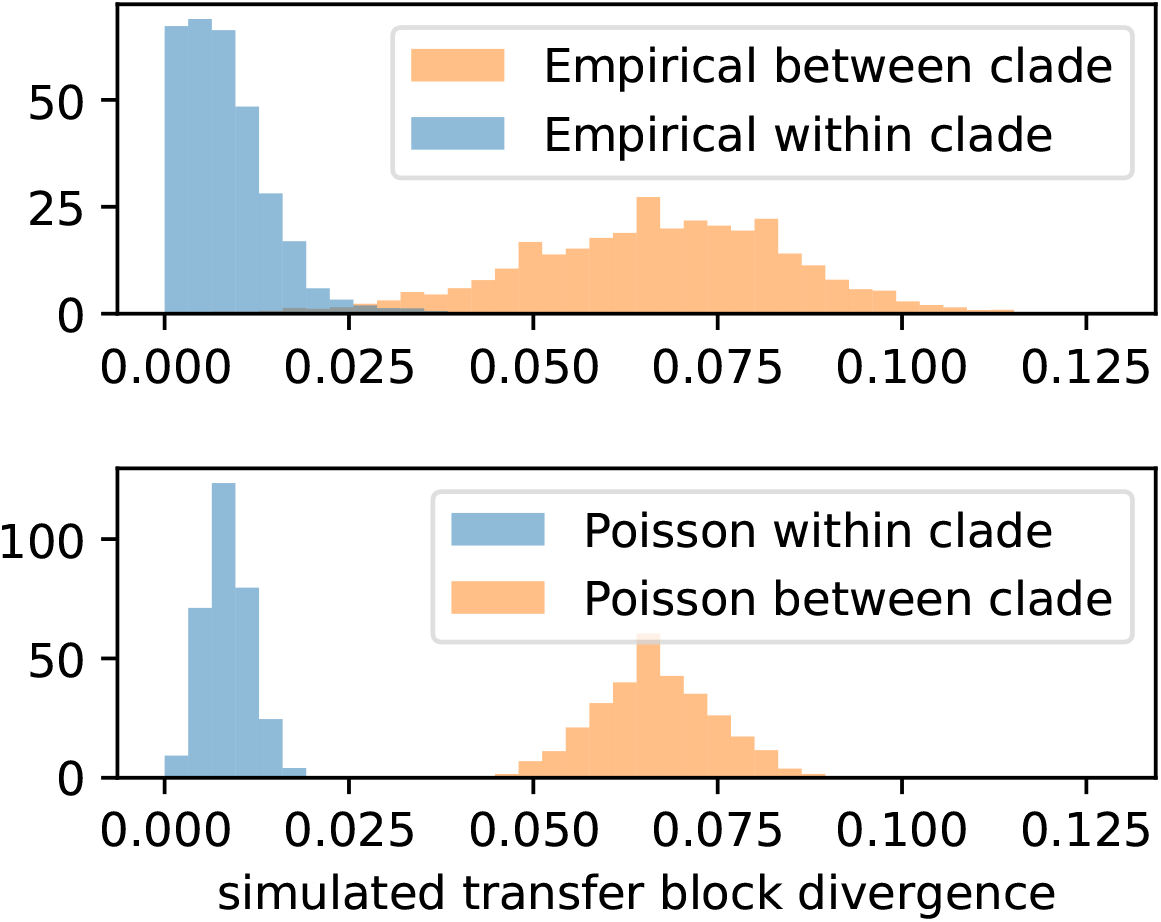
Empirical distributions of local divergence are broader than matching Poisson distributions. The top panel shows the empirical distribution of local pairwise divergence in *B. vulgatus*, computed by sampling blocks of 1000 synonymous sites as described in S1 Text 3.1. The bottom panel shows two corresponding Poisson distributions whose means match the empirical distributions above. The Poisson distribution does not capture the broad variation observed in the empirical distribution.

**Fig S7.**
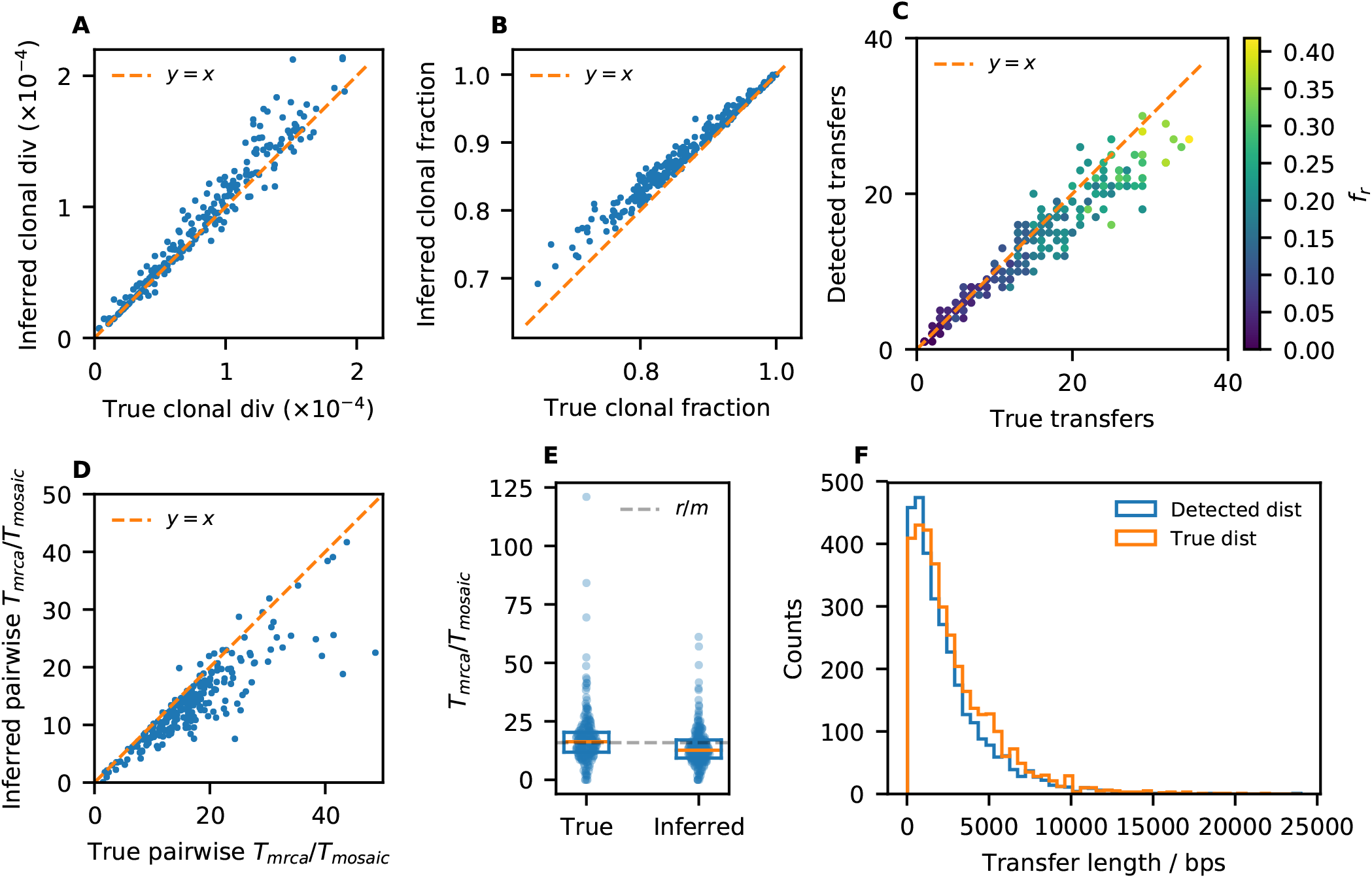
Validation of Close-Pair HMM (CP-HMM) algorithm using simulated data. A total of 256 pairs of close strains were simulated across a range of divergence times, using parameters matching *Bacteroides vulgatus* (S1 Text 3.2). (A) Correlation between the inferred clonal divergence and the true clonal divergence, 2*μT*. (B) Correlation between the inferred clonal fraction and the true clonal fraction. (C) Correlation between the number of detected transfers and the true number of transfers. Colors show the approximate fraction of recombined regions, defined as the true total length of recombination events divided by the genome length (overlapped regions will be counted more than once). (D) Correlation between the inferred *T*_mrca_/*T*_mosaic_ and the true *T*_mrca_/*T*_mosaic_ for each simulated pair. Panels (A-D) all show good agreement except for the most diverged pairs, which potentially reflect the influence of overlapping transfer events. (E) Distribution of true and inferred *T*_mrca_/*T*_mosaic_ for all simulated pairs. Box and orange lines indicate the interquartile range and median, respectively. Grey dashed line indicates the ground truth *r*/*m* value calculated from Eq. (S14) using the simulated parameters. These results show the pairwise estimates of *r*/*m* (or *T*_mrca_/*T*_mosaic_) can span a wide range due to Poisson sampling of mutations and recombination events, even in simulations with a single *r*/*m* value. (F) Distribution of detected transfer lengths vs the ground truth, showing good agreement. Detected transfers are expected to be slightly shorter than the ground truth, because the region between the last SNV and the end of a transfer (or between the start and the first SNV) is impossible for the HMM to detect.

**Fig S8.**
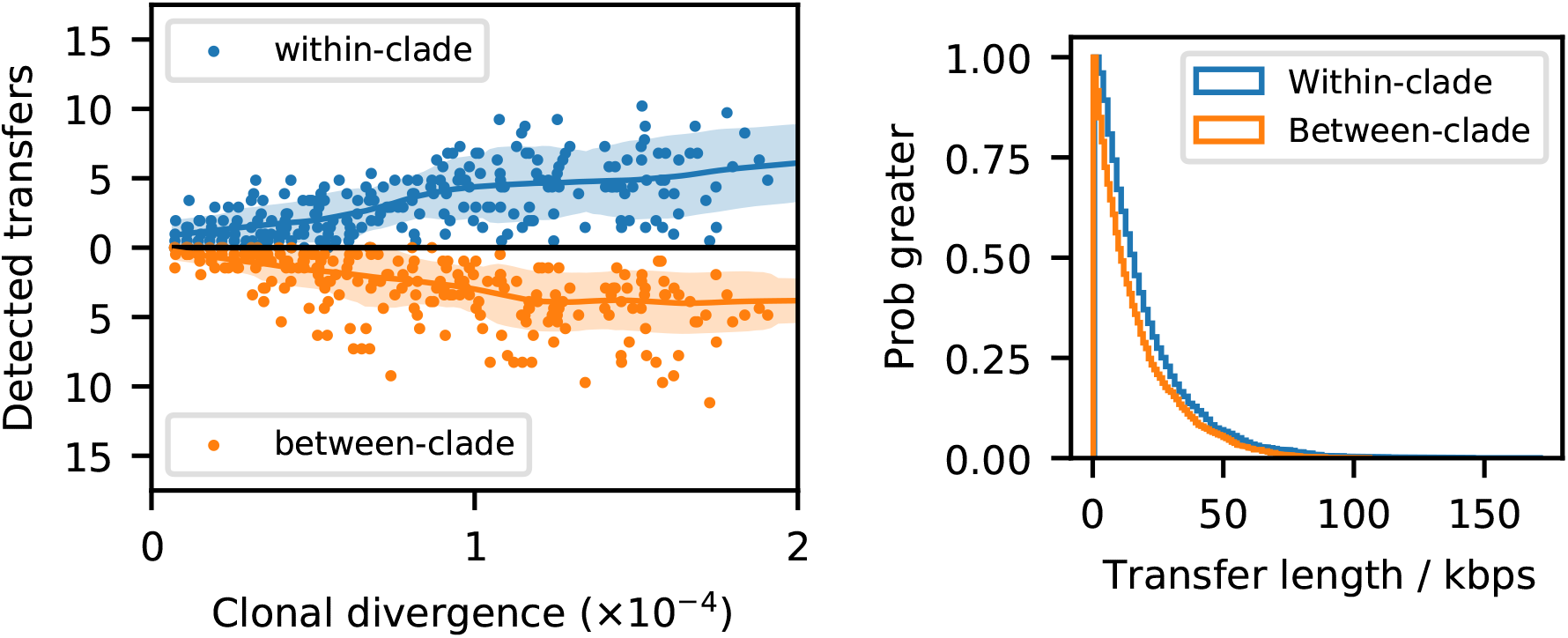
Comparing recombination dynamics in *B. vulgatus* using simulated data. Analogous versions of Fig. 2C-D created for the simulated data in Fig. S7 (S1 Text 3.2). In the simulation, between-clade and within-clade transfers are set to have the same rate and transfer length distribution. The simulated data therefore allow us to test whether the patterns in Fig. 2 were caused by effects such as a detection bias toward toward high divergence transfers, or the size difference between the two *B. vulgatus* clades. The number of detected transfers in the simulated data are comparable between the two transfer types, in contrast to the 5-fold difference in Fig. 2C; similarly, there is only a small (< 30%) bias in the length of the between-clade transfers, in contrast to the 7-fold reduction observed in data. The distinct patterns observed in *B. vulgatus* therefore cannot be explained by biases due to divergence or sampling of the two clades. These simulated data also show that using a single recombination rate does not generate large numbers of transfers at short divergence times, in sharp contrast with the observed data in Fig. 2C.

**Fig S9.**
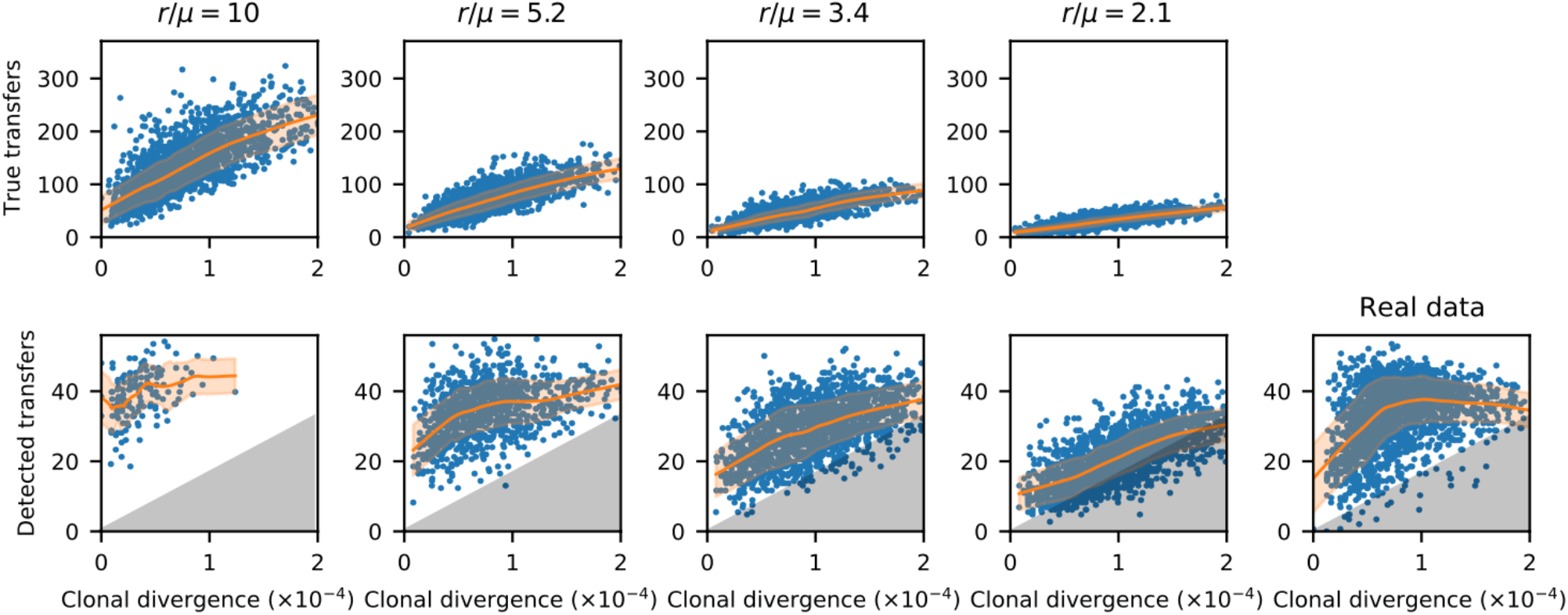
CP-HMM results for simulations approximating *A. putredinis.* We performed simulations using three different recombination rates while matching the total number of samples, the genome length and the mean recombination length of *A. putredinis* (S1 Text 3.5). The bottom panels show analogous versions of Fig. 3C computed from these simulated datasets; the original data from Fig. 3C are reproduced on the right for comparison. The same region of low recombination rate are shaded to highlight the difference between *r*/*μ* = 5.2 and real data. The top panels show the true number of transfers versus the true clonal divergence in the simulations (trend lines and confidence intervals are computed in the same manner as Fig. 3C). None of the simulated datasets reproduces the broad variation observed in *A. putredinis*, which contains pairs with both large numbers of transfers at short times and small numbers of transfers at long times. This suggests that a single recombination rate cannot explain the observed data.

**Fig S10.**
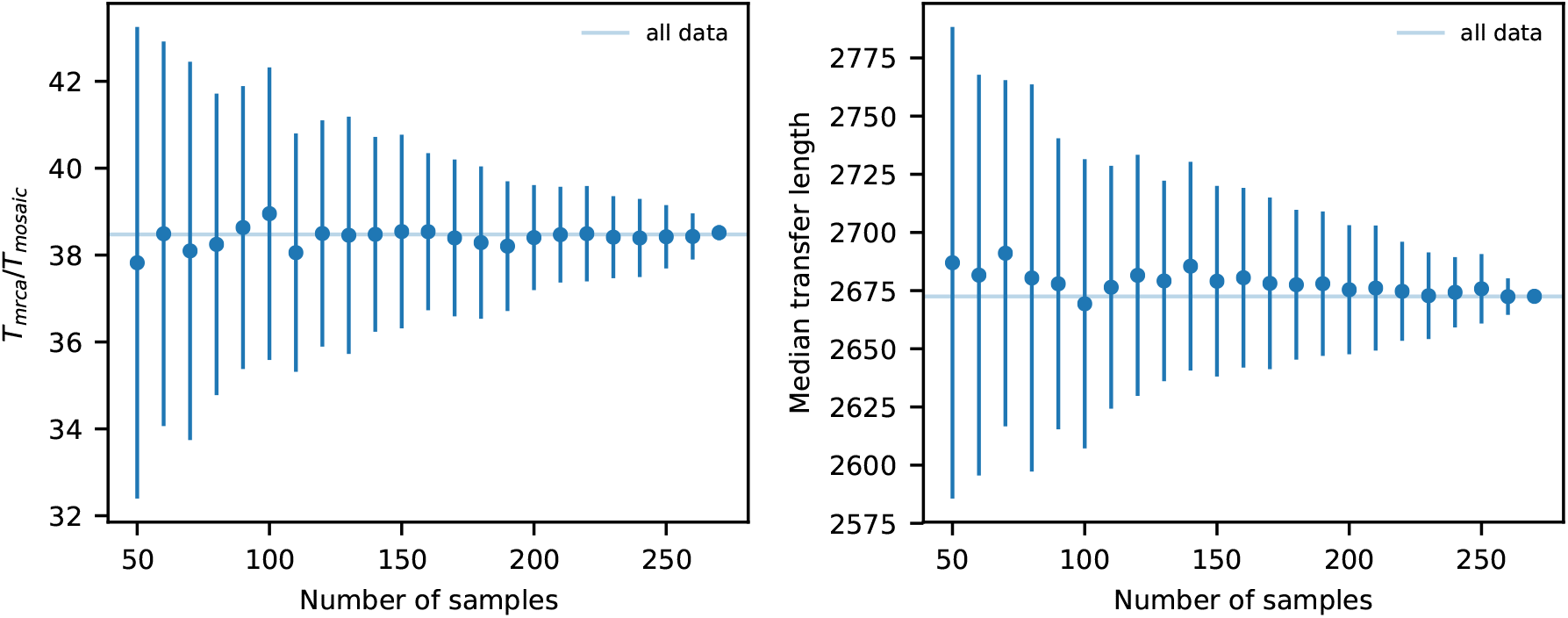
Dependence of CP-HMM results on the sample size. A series of downsampling experiments were performed using the *Alistipes putredinis* dataset, and CP-HMM was applied to each of the downsampled dataset. Left and right panels show the inferred *T*_mrca_/*T*_mosaic_ ratios (left) and the median inferred transfer lengths (right). Bars denote one standard deviation calculated from 100 replicates. The parameters inferred from the downsampled datasets remain very close to the values obtained from the full dataset.

**Fig S11.**
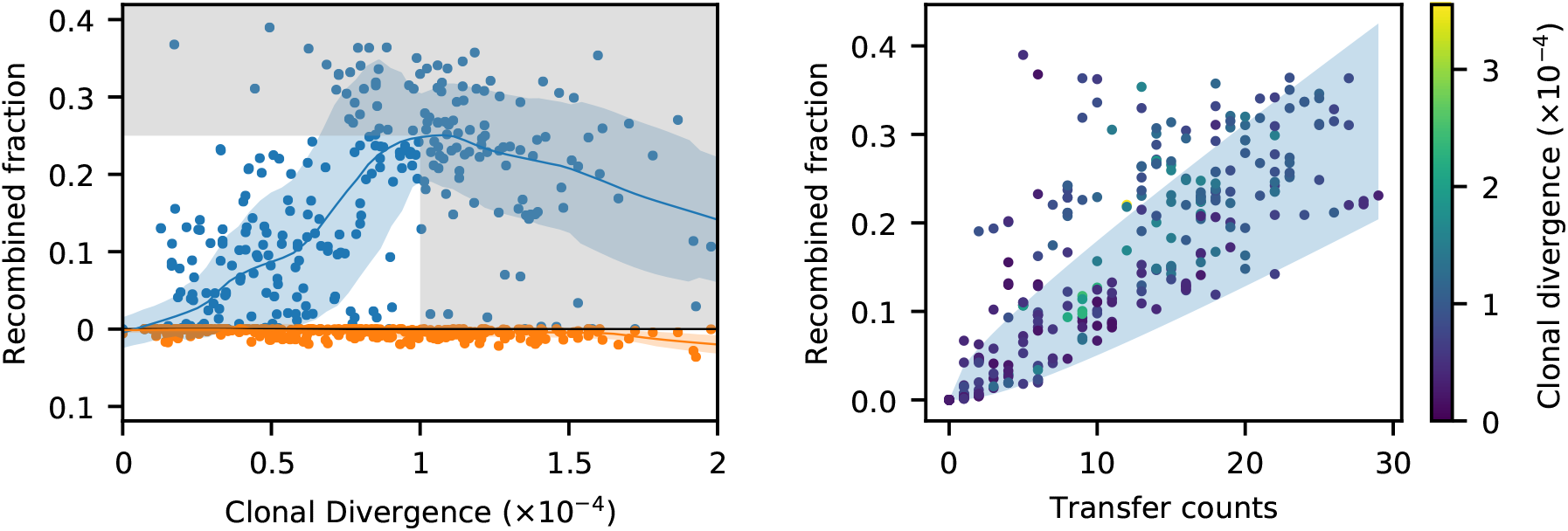
Total recombined fraction vs divergence in *B. vulgatus*. Left: analogous version of Fig. 2C showing the total fraction of the core genome composed of recombined regions as inferred by CP-HMM. In contrast to the number of transfers, this metric remains consistent when neighboring transfers overlap, providing an alternative check on the general patterns we observed in Fig. 2C. Points show the CP-HMM results for all pairs of strains with at least >50% identical blocks. Grey regions denote the points that were excluded by our subsequent filtering steps (S1 Text 3.3). Comparing these results with Fig. 2C shows that many important trends (e.g. the non-linear accumulation of transfers and the reduced recombination between clades) are also observed in this alternative metric. Right: Scatter plot showing the correlation between the number of detected transfers and the total recombined fraction. Each point is colored by the inferred clonal divergence for that pair. Blue shaded regions denote 95% confidence intervals for a null model where the length of each transfer is drawn from an exponential distribution with mean measured in Fig. 2D. This comparison reveals that in some pairs, a small number of transfers account for an unusually large fraction of recombined genome. Examples of these outliers are shown in Fig. S12.

**Fig S12.**
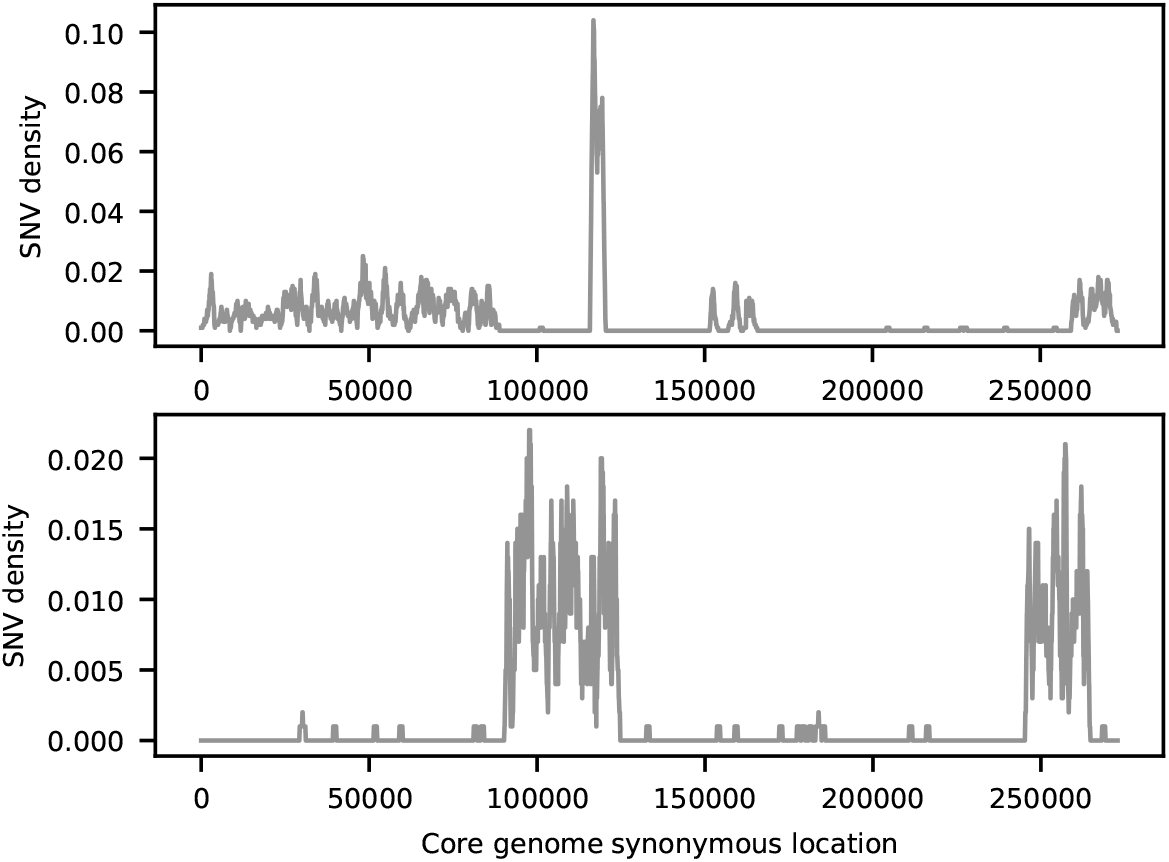
Examples of extremely long recombination events. Top and bottom panels show the synonymous divergence profiles of two pairs of closely related *B. vulgatus* strains with abnormally long recombination events. Top panel shows an example of a recombined region covering nearly half of the genome, while leaving extended regions unmodified. This pattern is unlikely to be generated by the merging of multiple independent transfers: for a typical recombination length of ≈2,900 synonymous sites, ≈40 such transfers must occur exclusively within a particular half of the genome (*P* < 10^−12^). Bottom panel shows an example of two isolated recombination events covering more than ≳50,000 synonymous sites (equivalent to ∼ 350kb in total genome length). Some of these long events appear to have multiple peaks in their associated SNV profiles, suggesting that they might be composed of several smaller (correlated) transfers. We also found similar long recombination events between pairs of cultured isolates (Fig. S13).

**Fig S13.**
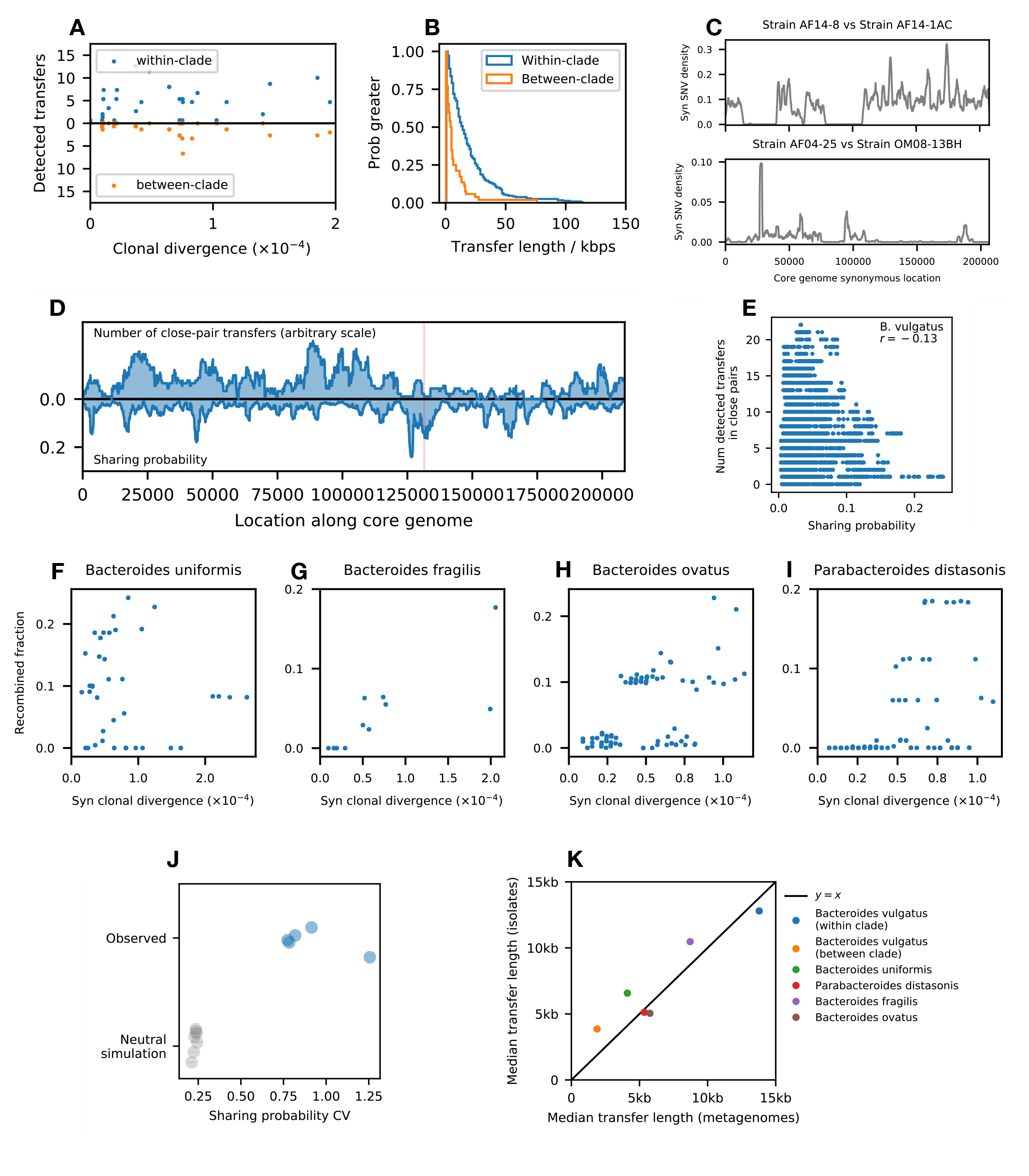
CP-HMM analysis of isolate genomes from commensal gut species. CP-HMM was applied to a total of 314 previously sequenced isolate genomes from 5 species of commensal gut bacteria (S1 Text 3.10). (A-B) Analogous versions of Fig. 2C-D for *Bacteroides vulgatus* isolates. The reduction in the rate and length of between-clade transfers relative to within-clade is quantitatively reproduced in the isolate data. (C) Examples of long putative transfer events in *B. vulgatus*. Both panels show the density of 4D SNVs between a pair of strains along the core genome. All four strains were originally sampled in Ref. 98. Top: Two strains sampled from the same host share two anomalously long segments, while retaining high divergence in the rest of the genome. Bottom: Two closely related strains sampled from unrelated hosts exhibit recombined regions that are concentrated in only half of the genome. This pattern is very similar to Fig. S12, suggesting a very large recombination event in the time since they last shared a common ancestor. (D-E) Analogous version of Fig. S30A-B for *B. vulgatus* isolates. The location of the within-host sweep event in Fig. 5C, showing an elevated sharing fraction; note that the location along core genome is different because a different reference genome was used for the isolate analysis. (F-I) Analogous versions of Fig. S15 obtained from isolate data. The apparent rates of accumulation of recombination events are consistent with our previous results using metagenomic data. (J) Analogous version of Fig. 6D. The CV of the sharing probability among isolate genomes is again much higher than neutral simulations. (K) Scatter plot of median transfer length inferred using metagenomic data vs isolates, showing good agreement.

**Fig S14.**
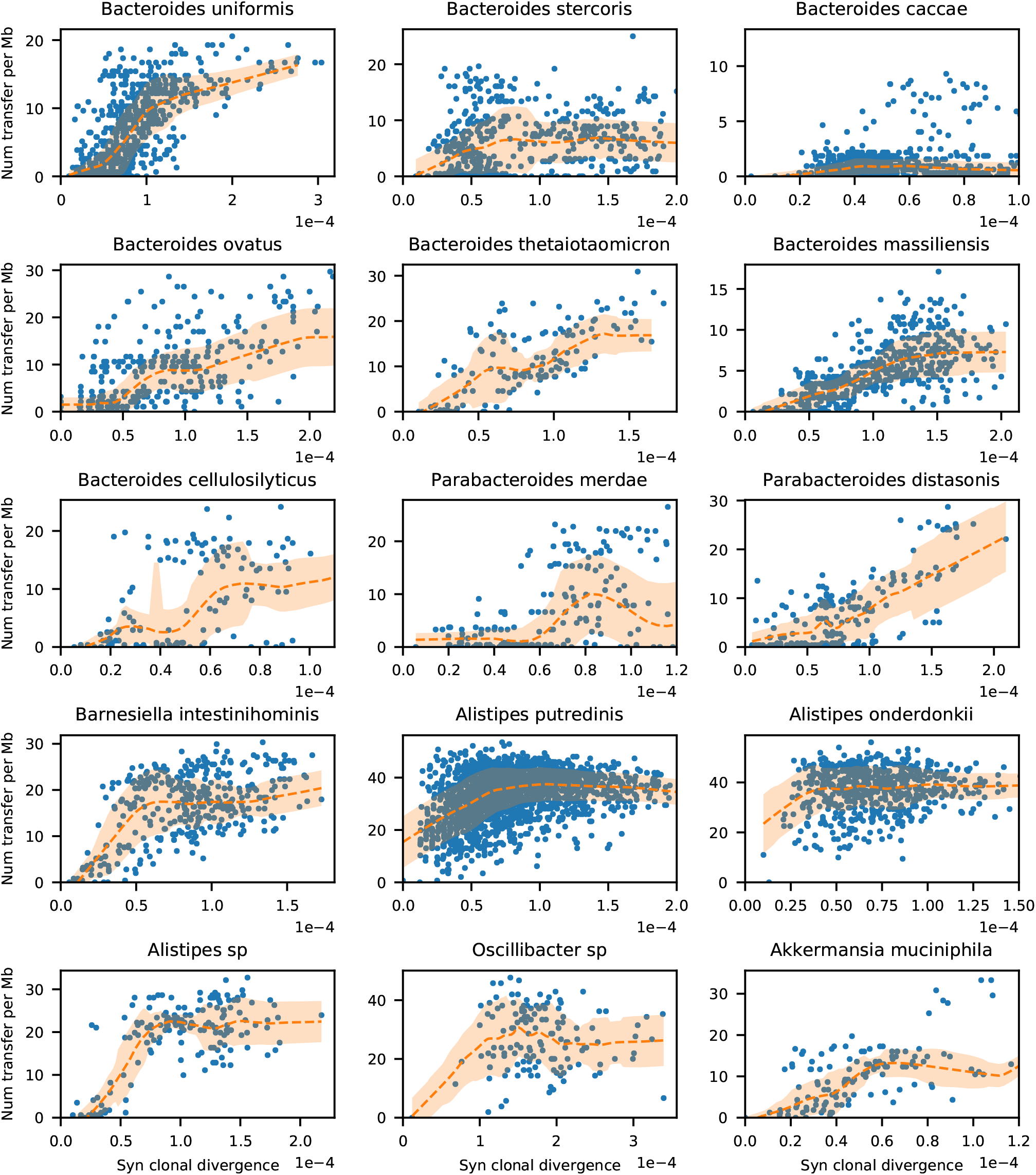
Number of transfers vs divergence for other prevalent gut species. Analogous versions of Fig. 3A-C for all species with > 100 closely related pairs (i.e. inferred clonal fraction > 75%). *B. vulgatus* is plotted separately in Fig. 2.

**Fig S15.**
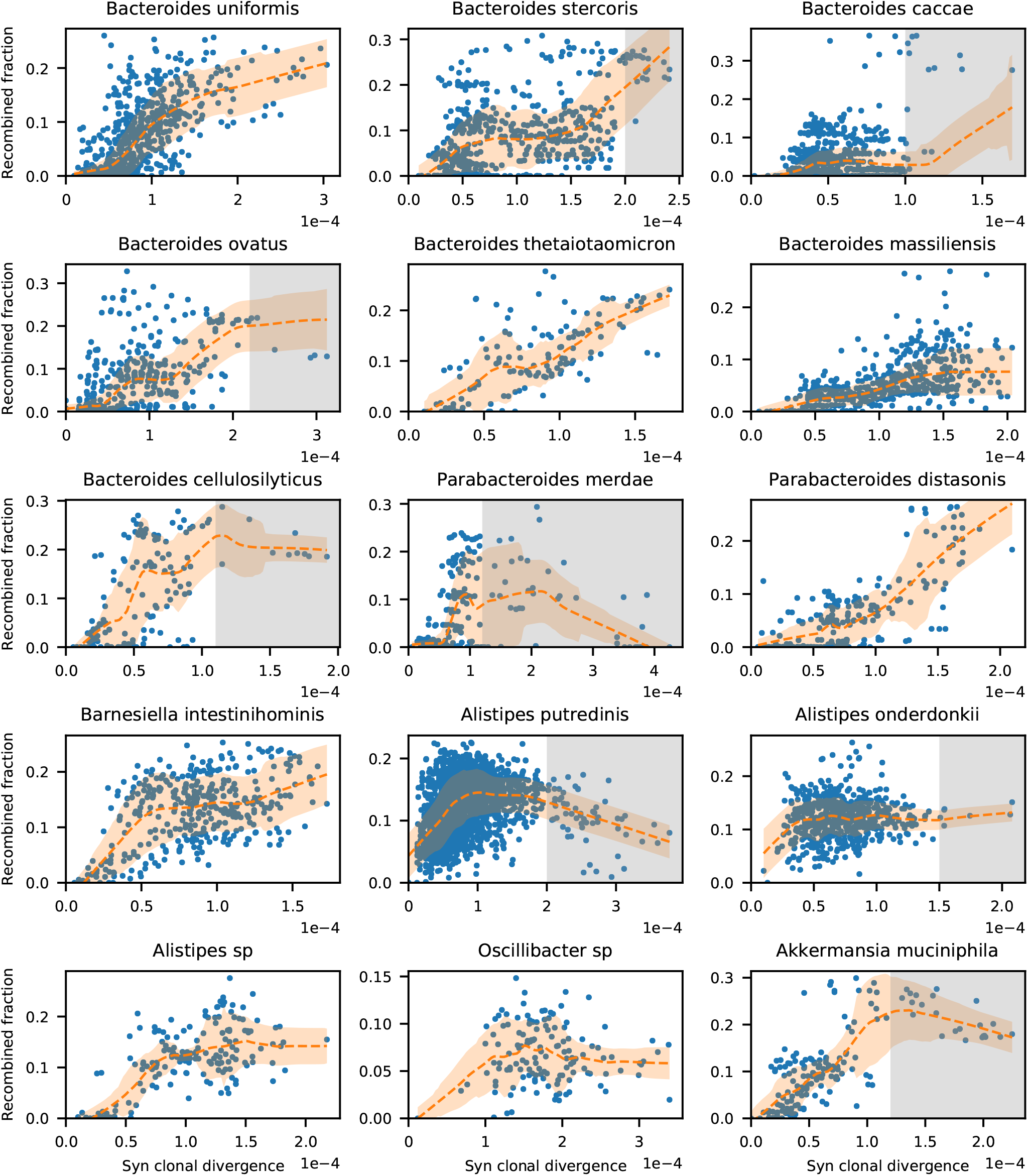
Total recombined fraction vs divergence for other prevalent gut species. Analogous version of Fig. S14 showing the total length of all recombined regions (similar to the *B. vulgatus* example in Fig. S11). Comparisons with the total number of counts in Fig. S14 show that the overall trends revealed by these two metrics are similar for most species, including key observations such as the high degree of variation in *A. putredinis*. Interestingly, *B. caccae* has a larger spread in this metric than in the number of transfers (Figs 3 & S14), which is consistent with the its broad distribution of transfer lengths (Fig. 3E).

**Fig S16.**
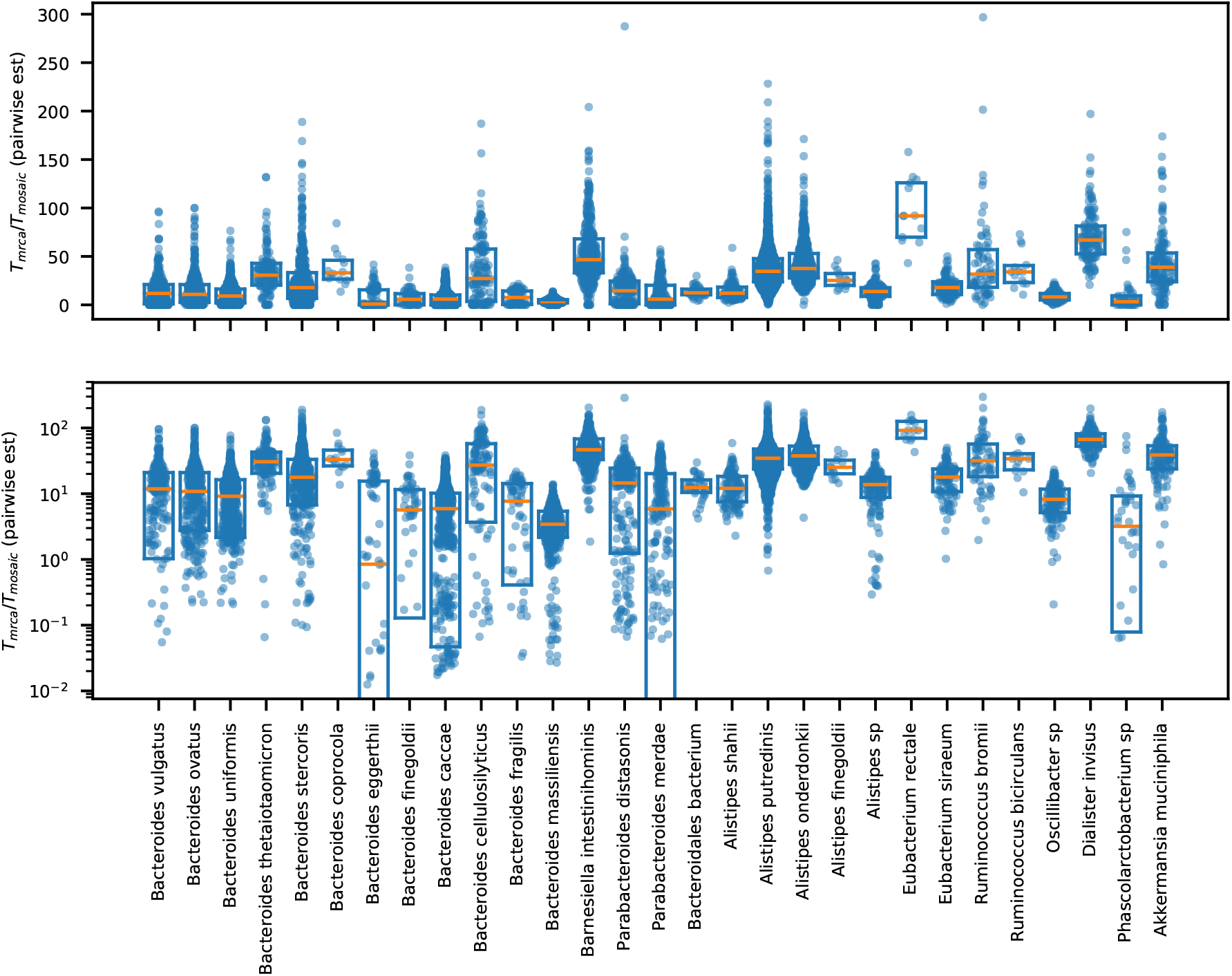
Distribution of. *T*_mrca_/*T*_mosaic_ **estimates from different pairs of strains.** Top: Symbols show the estimated values of *T*_mrca_/*T*_mosaic_ for all close pairs using the expression in Eq. (S16); box plots indicate the median and interquartile range. This metric is roughly equivalent to the “*r*/*m*” metric in the context of simple neutral models [25, 39]. Bottom: same data on a log scale.

**Fig S17.**
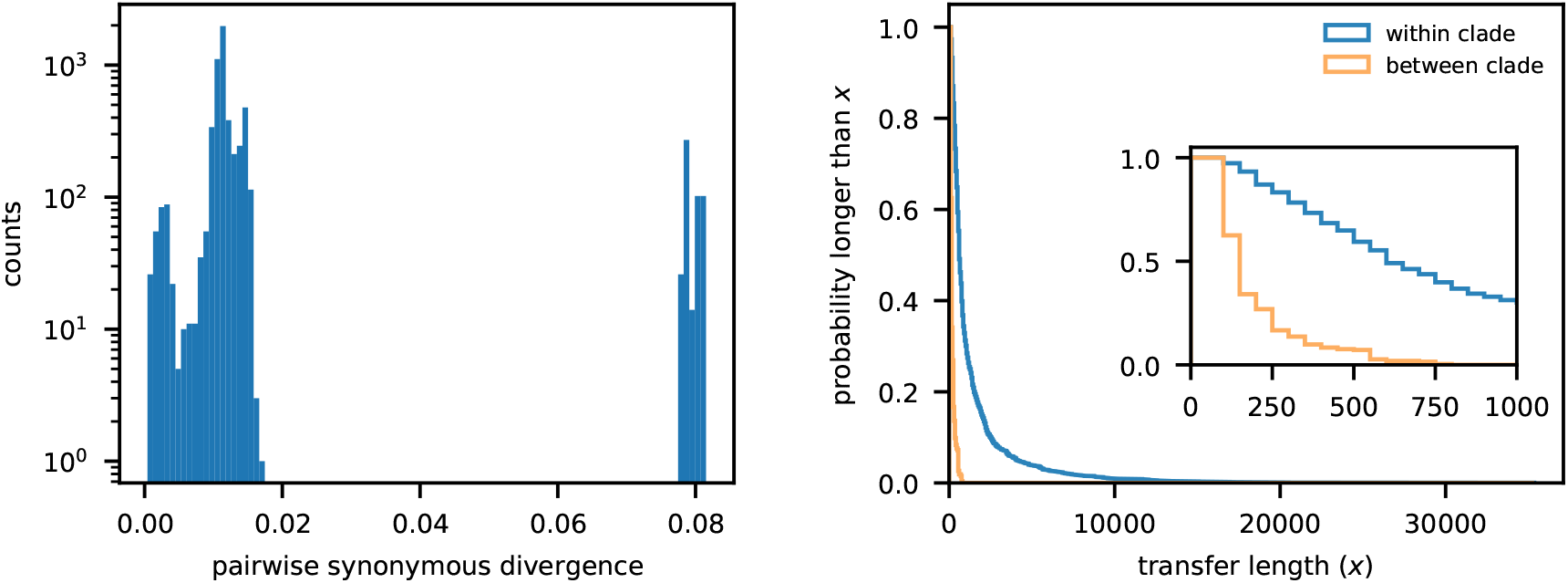
Within- and between-clade transfers in *A. shahii*. Left: Pairwise divergence distribution of *A. shahii*. Strains separate clearly into two major clades, similar to *B. vulgatus* (Fig. S3). Right: Analogous version of Fig. 2D constructed for *A. shahii*. As in Fig. 2D, within- vs between-clade transfers are inferred based on divergence of the transferred region. Inset shows a zoomed-in version to highlight the between-clade distribution. This shows that the reduction in transfer length for between-clade transfers is not specific to *B. vulgatus*.

**Fig S18.**
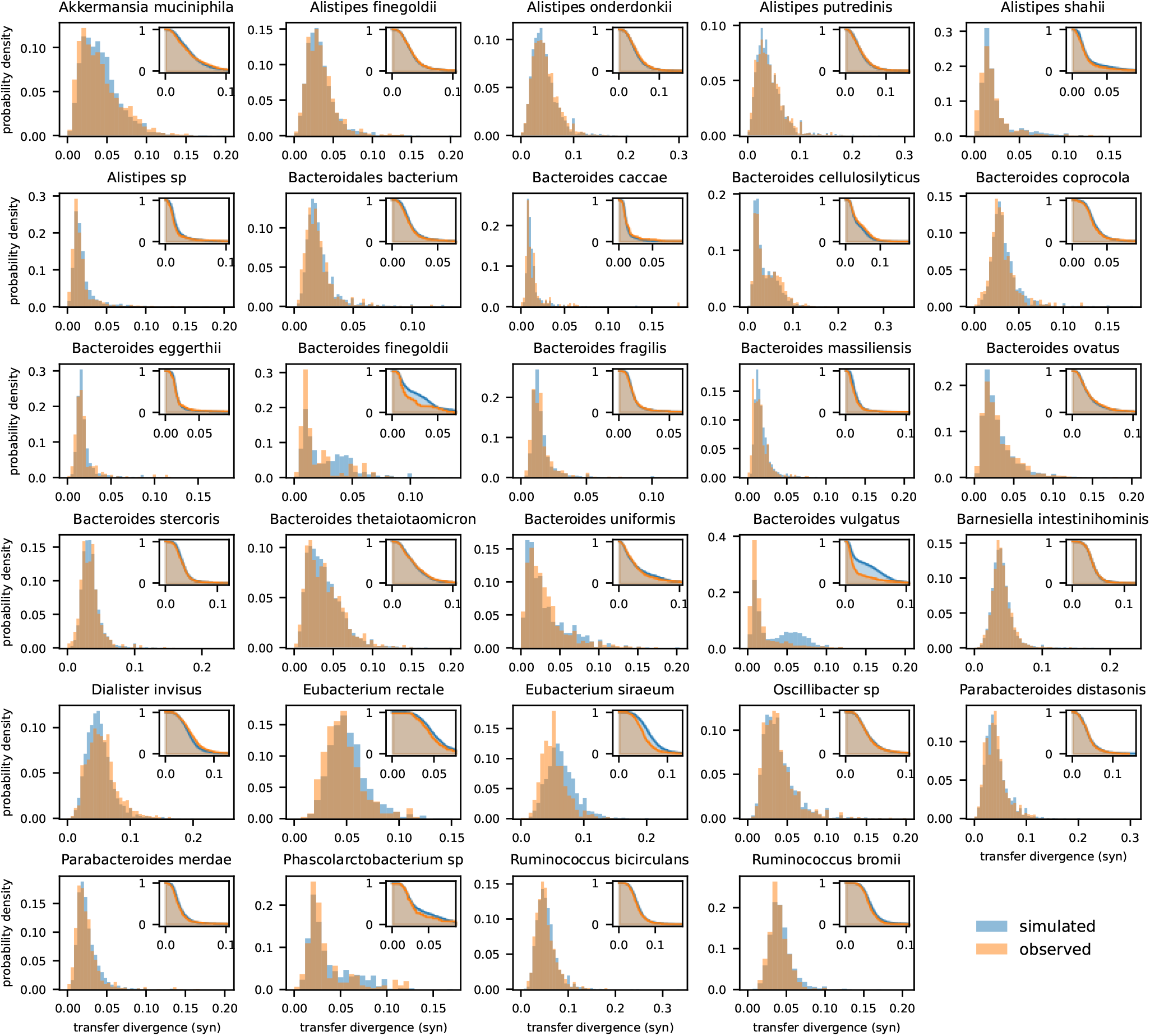
Distribution of divergence of detected transfers for each species. Orange histograms show the synonymous divergence of all detected transfers for all species shown in Fig. 3. For comparison, the blue distributions show the expectation for a null model in which we simulated transfers in the same set of close pairs by randomly sampling donated sequences from the larger set of genomes in our cohort. To preserve potential variation along the genome, the locations and lengths of the simulated transfers were chosen to match the actual detected events. For most species, the empirical distribution of divergence closely follows this null model, consistent with the picture that the accumulated transfers were obtained from random strains in the population. However, certain species (e.g. *B. vulgatus*, *B. finegoldii*) that have apparent population structure, as indicated by a second peak in the simulated histogram, show a significant decrease in transfers donated from the other clade. Importantly, this decrease of transfer efficiency is not seen for other closely related species with a similar range of transfer divergences (e.g. *B. thetaiotaomicron*).

**Fig S19.**
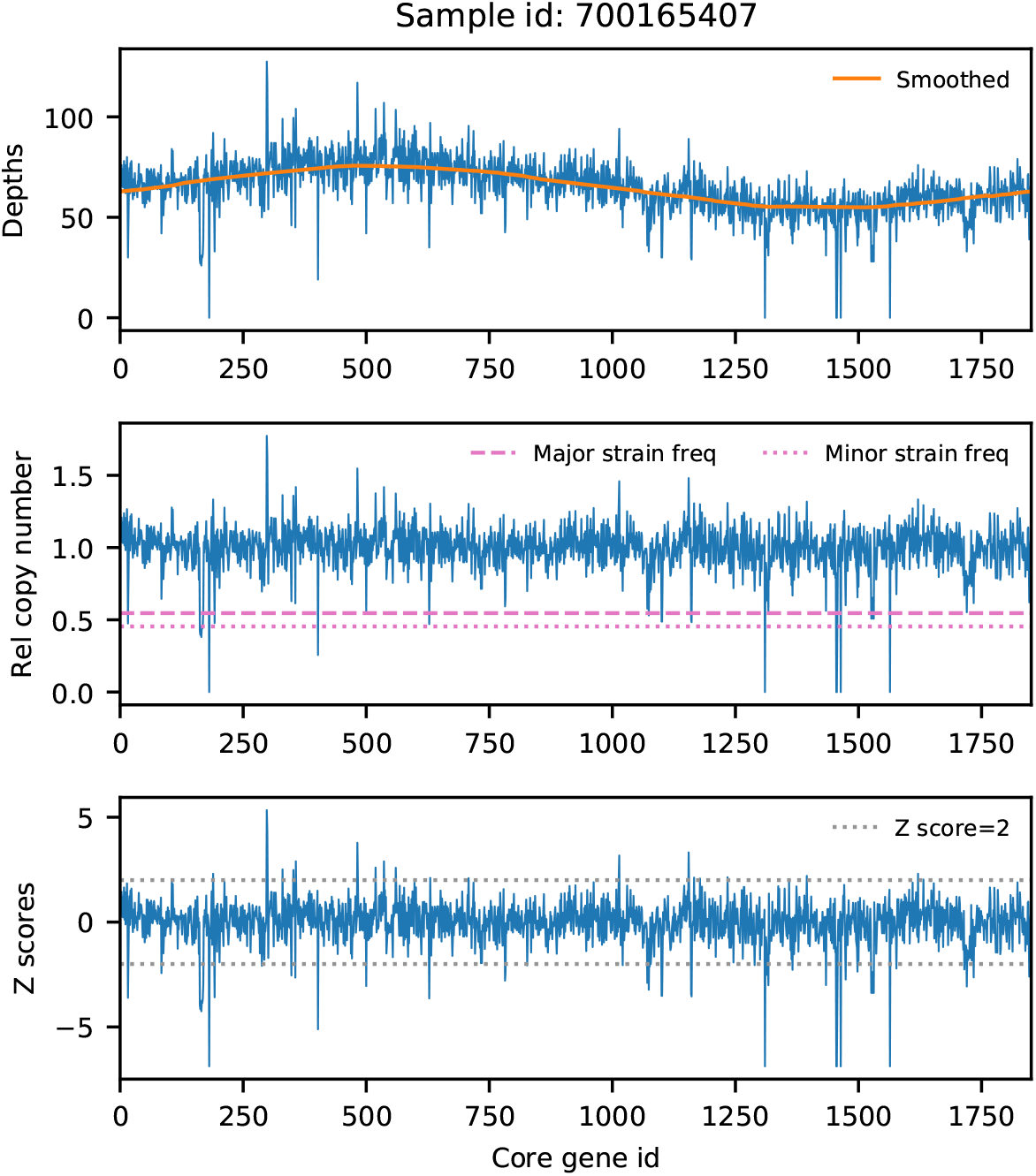
Filtering within-host gene deletion events using read depths. An example of our filtering algorithm in S1 Text 4.1 applied to a single sample that is co-colonized by two *B. vulgatus* strains. Top: Median read depth for all core genes, sorted according to their location on the reference genome. Orange line shows a moving average of 500 genes. Mid: Relative copy number of all core genes, computed as the median read depth of each gene divided by the moving average in the top panel. Dashed and dotted lines represent the inferred major and minor strain frequencies, respectively. If a gene is only present in the major or minor strain, then its relative copy number will be approximately equal to that strain’s frequency. If a gene is missing from both strains, the relative copy number will be zero. Bottom: Z-score of all genes computed using the relative copy number in the middle panel. Genes with |*z*| > 2 are flagged as potential gene deletion events and are filtered from all downstream analyses.

**Fig S20.**
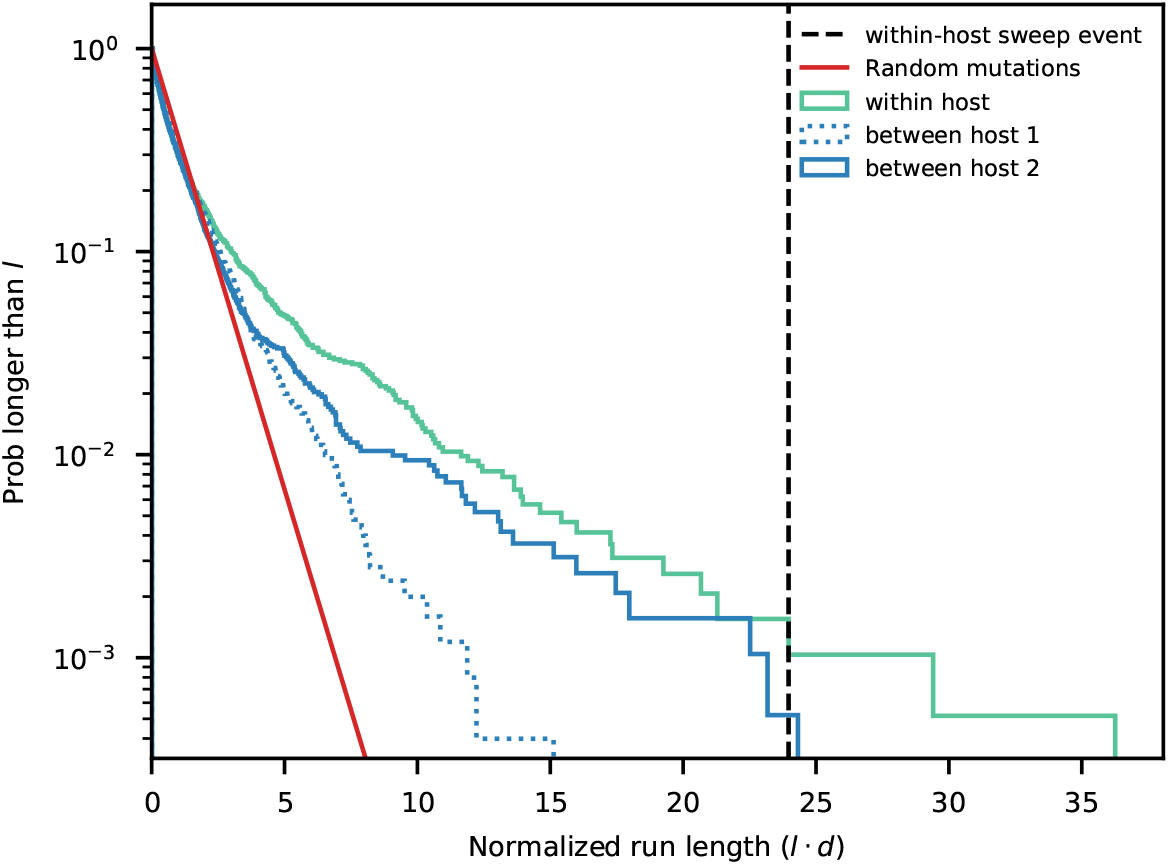
Distribution of shared fragment lengths for the genomes in. Fig. 5B. All homozygous runs of zero SNVs are identified for the pairs shown in Fig. 5B, and the cumulative distribution of run lengths is shown here. The run length is normalized by the pairwise divergence *d*. Only the second time point of the within-host example is plotted. For comparison, the black dashed line indicates the length of the within-host sweep event in Fig. 5C, while the red line shows the expected distribution if the SNVs were randomly distributed along the genome (S1 Text 4.2). These data show that random pairs of strains can sometimes share fragments that are even longer than the within-host sweep event (e.g. between-host pair 2).

**Fig S21.**
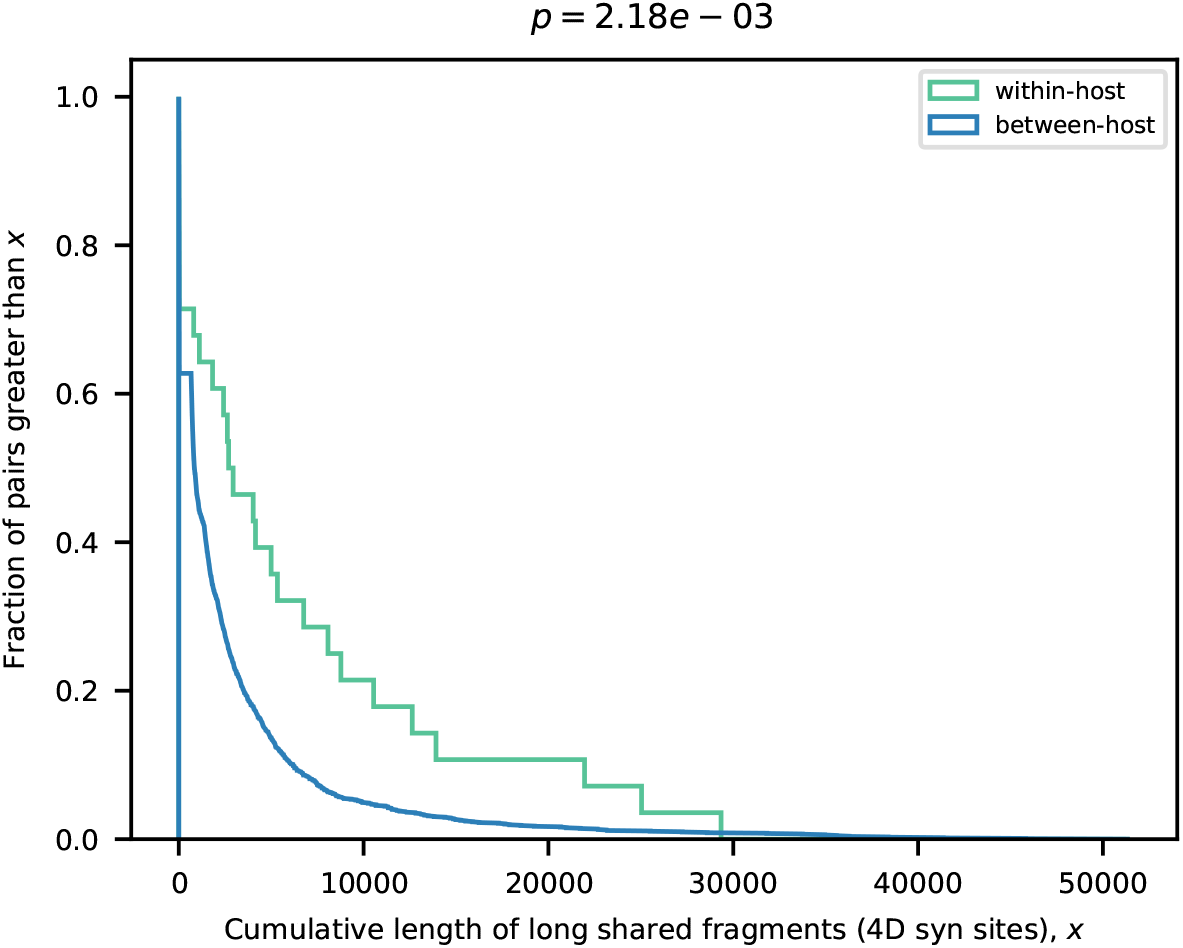
Distribution of the total length of long homozygous runs for *Eubacterium rectale*. Analogous version of Fig. 5D using an alternative test statistic, based on the total length of runs longer than 600 synonymous sites (equivalent to ℓ^∗^ · *d* ≈ 30). Similar to the “max run” statistic in Fig. 5D, this new metric also demonstrates that *E. rectale* is statistically enriched for within-host sharing.

**Fig S22.**
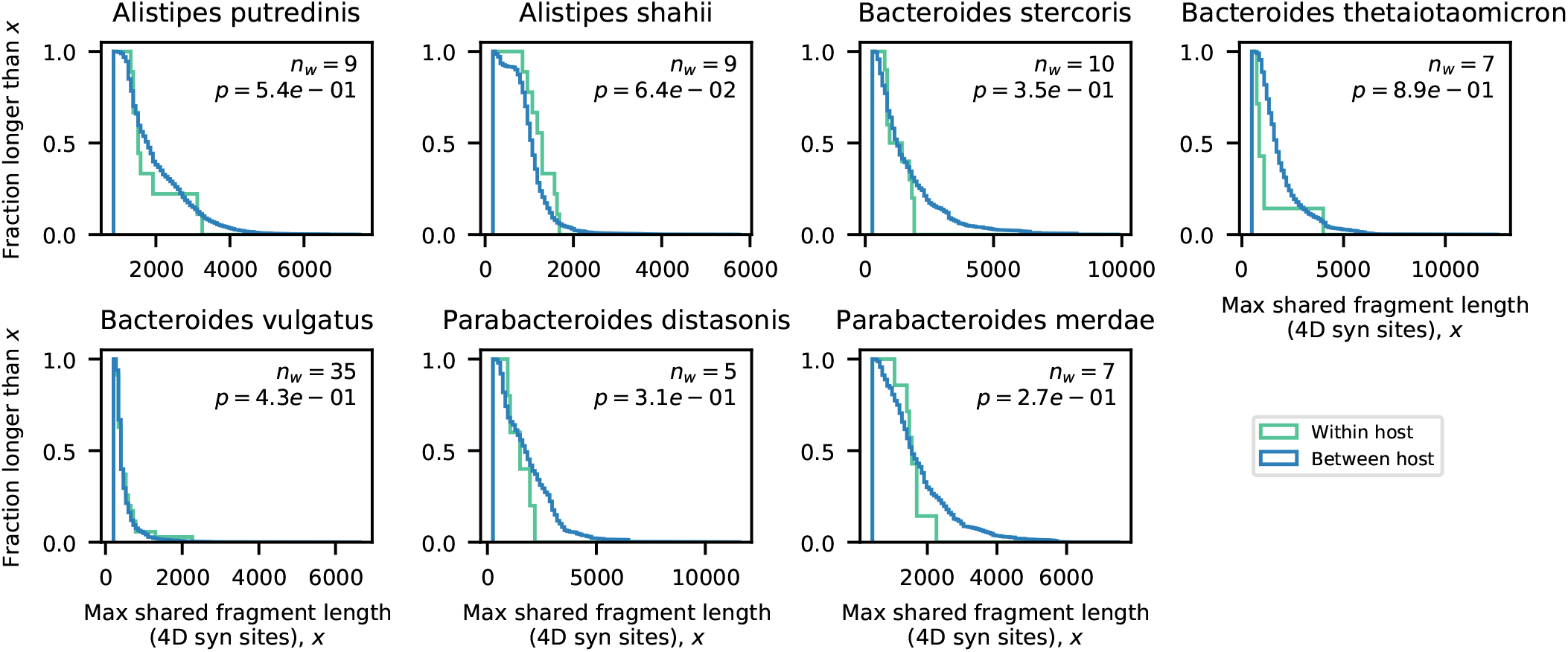
Distribution of the longest homozygous runs within hosts. Analogous versions of Fig. 5D for the remaining species that had at least 5 hosts that passed our filtering criteria (S1 Text 4.1). None of the species here, including highly recombinogenic ones such as *A. putredinis*, are statistically enriched for within-host sharing.

**Fig S23.**
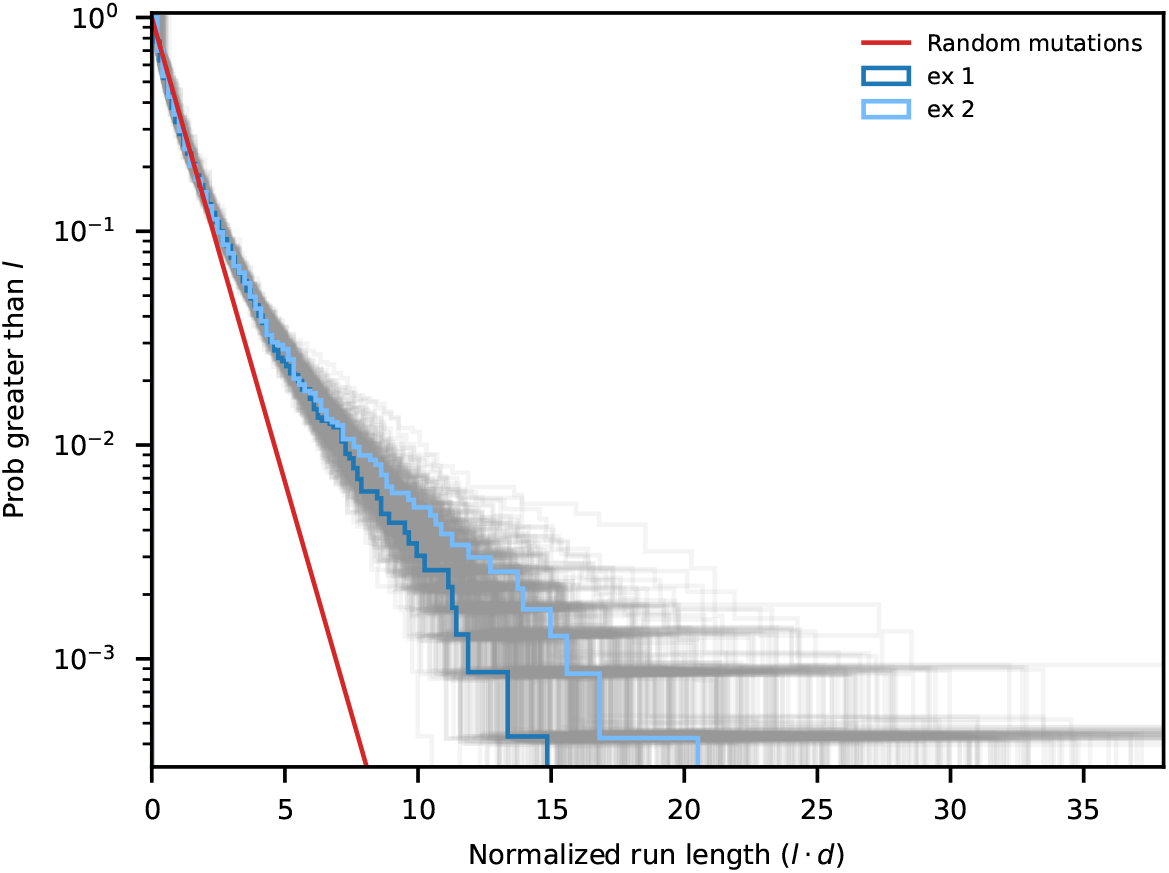
Distribution of shared fragment lengths in neutral simulations. Analogous version of Fig. S20 constructed from 500 simulated genomes from FastSimBac. Simulation parameters are identical to the lower right panel of Fig. S2. Only pairs with < 10% of identical blocks are shown here. The distribution of two random pairs are highlighted in blue as examples, and all others are plotted in grey. As expected from our approximate formula for the probability of observing long sharing fragments (Eq. S28), it is possible to find pairs in this neutral simulation that share a fragment as long as the within-host sweep event in Fig. S20.

**Fig S24.**
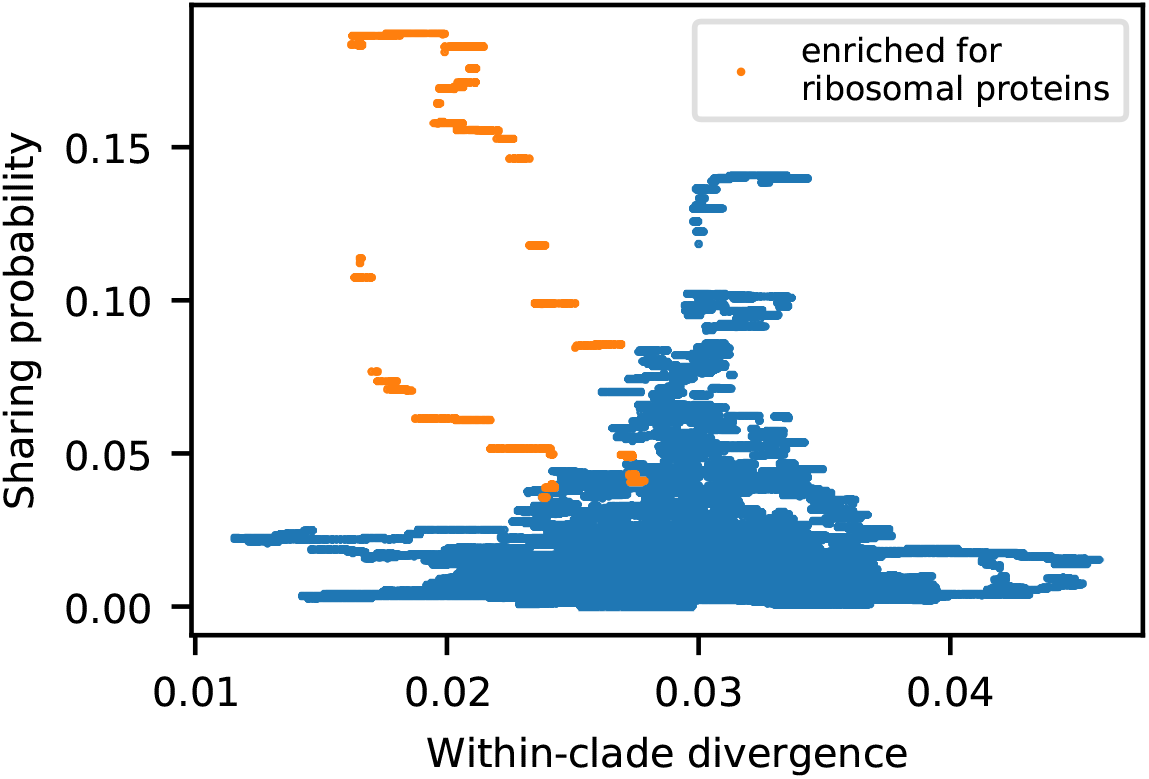
Local variation in the sharing of long fragments does not correlate with local divergence. Joint distribution the local sharing probability and divergence from Fig. 6C. These two variables are largely uncorrelated except for the region highlighted in orange, which corresponds to the major sharing peak around position 60,000. Interestingly, this region has the highest concentration of ribosomal proteins along the genome, which could potentially be the driver of this sharing hotspot.

**Fig S25.**
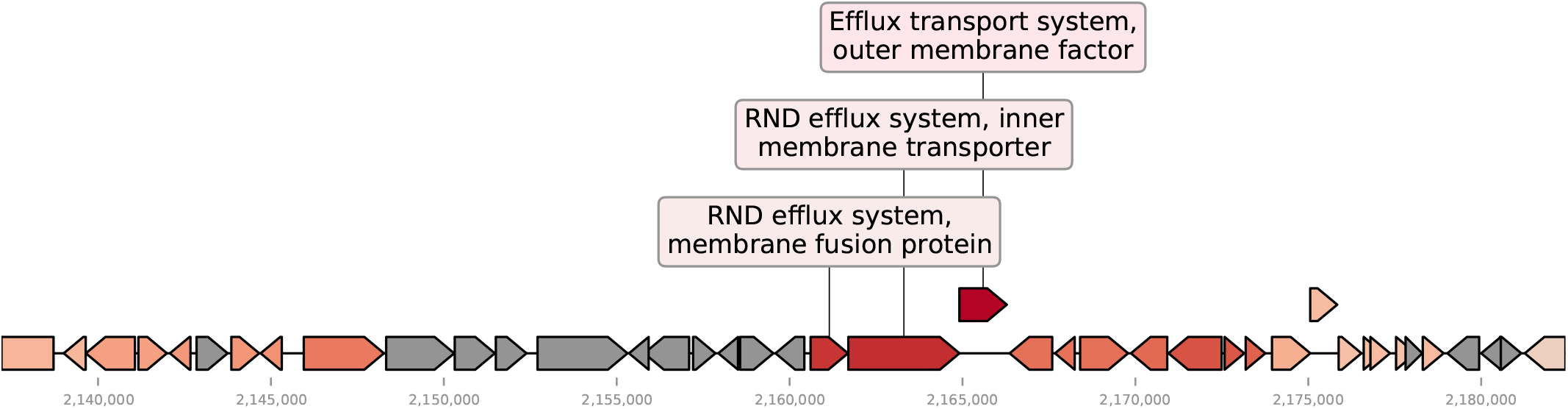
Global sharing probability of genes involved in the within-host sweep event in. Fig. 5C. Shown here is a portion of the *B. vulgatus* reference genome, roughly corresponding to the genes in the putative within-host sweep event in Fig. 5C (Table S5). Three genes encoding a RND efflux system are annotated with text boxes. Core genes are colored according to their global sharing probabilities computed in Fig. 6B, with warmer color representing higher frequency. Non-core genes are shown in grey. This detailed view reveals that the RND efflux system has the highest sharing probability in this region. This suggests that selection for different genetic variants of this efflux pump could be driving both the within-host sweep event and the global sharing hotspot.

**Fig S26.**
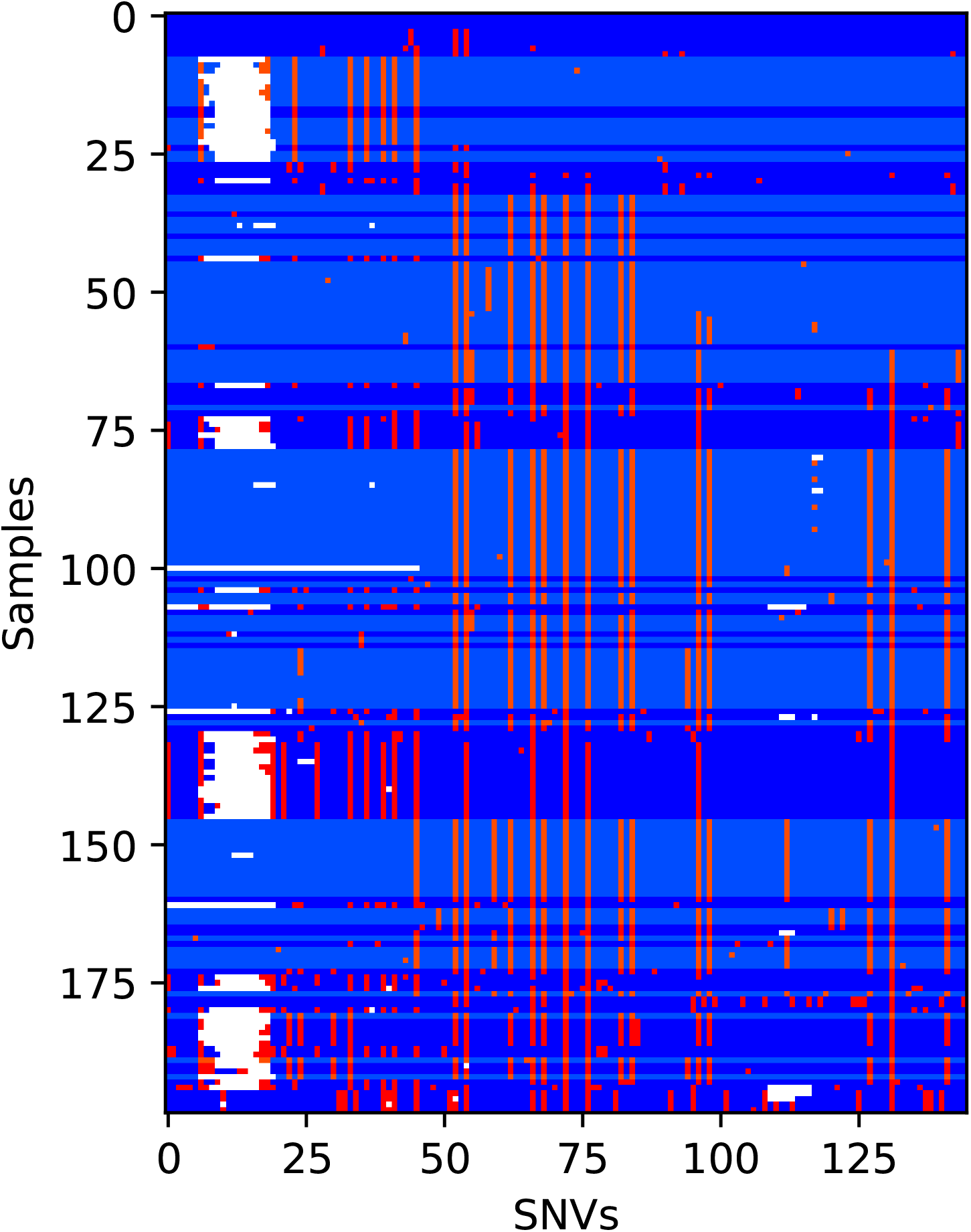
Multiple haplotypes contribute to a global sharing hotspot. Visualization of the observed haplotyes within the global sharing hotspot in Fig. S25. Columns represent all variable sites in three genes encoding an efflux pump in Fig. S25. Rows represent the “quasi-phased” strains sampled from unrelated hosts that share the same major clade as the within-host strains in Fig. 5C. Sites that match the within-host alleles are shown in blue, while those with different alleles are shown in red; white tiles denote missing data. Strain rows are sorted according to their sequence divergence relative to the within-host sequence in Fig. 5C. Strains rows are highlighted in lighter colors if they are in the pairs that contribute to the global sharing probability at these genes (Fig. 6). These data show that while a few strains share the same haplotype as the within-host strains (completely blue rows on the top end), the majority of strains contributing to the sharing hotspot do not share the same sequence. In addition, the highlighted regions show that multiple distinct haplotypes (light colored blocks with the same red locations) contribute to this sharing hotspot. This suggests that the sharing hotspots may be driven by soft selective sweeps of transferred genome segments.

**Fig S27.**
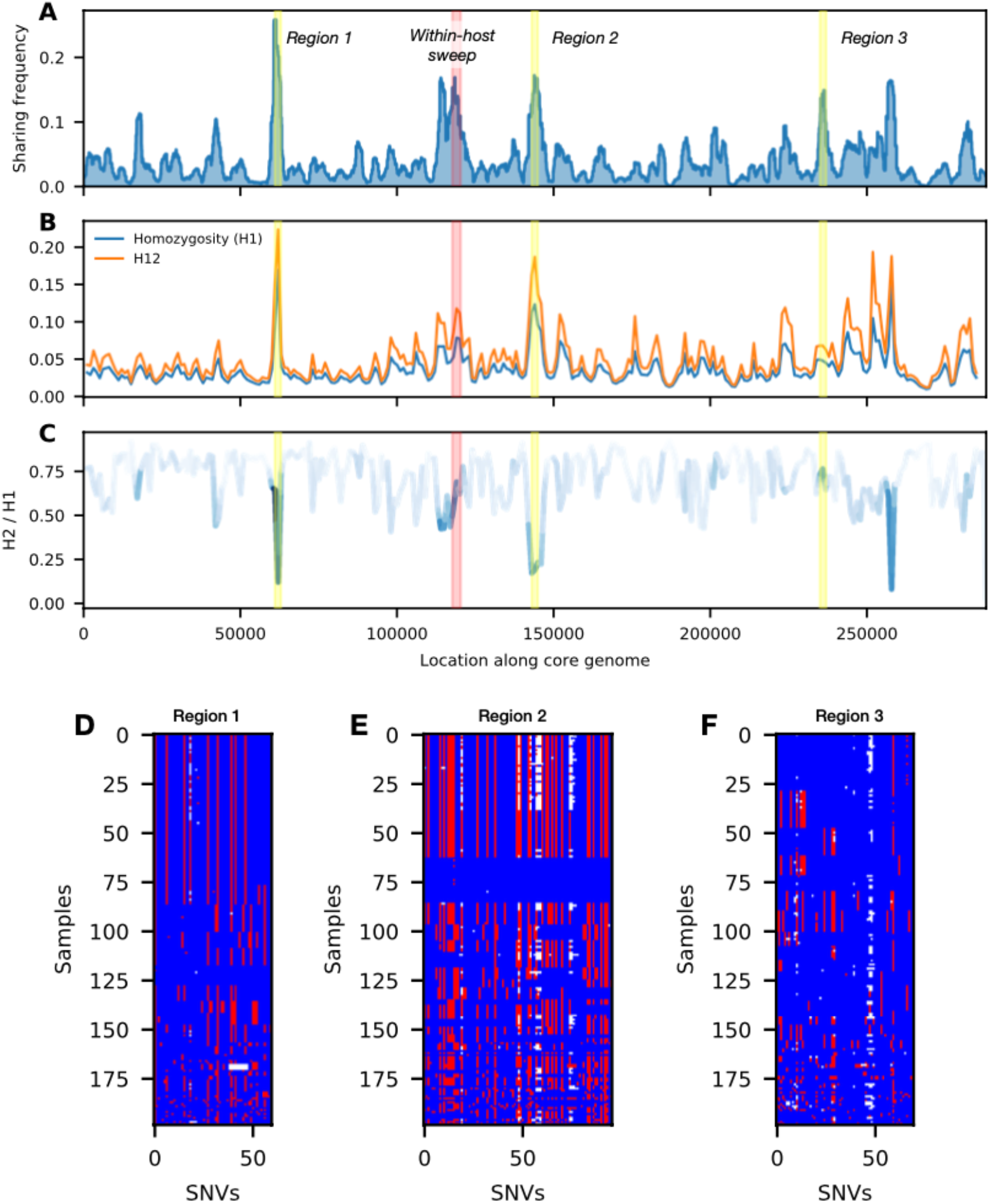
Comparison between sharing probability and haplotype homozygosity. The within-clade sharing landscape of *Bacteroides vulgatus* is compared with various selection tests based on haplotype homozygosity. (A): Same data as Fig. 6B. In addition to the within-host sweep event (Fig. 5C, three more 2kb regions corresponding to sharing hotspots are highlighted in yellow. (B) Haplotype homozygosity scan along the core genome. H1 is the conventional haplotype homozygosity, while H12 is the analogous statistic with the largest two haplotypes combined [99]; elevated regions of H1 or H12 indicate candidate regions undergoing selective sweeps, with H12 having better power for detecting soft sweeps. Scans were performed in 2kb windows, and haplotypes were defined by clustering samples with <2 SNV differences in the window. (C) Ratio between H2 (the haplotype homozygosity after excluding the largest haplotype [99]) and H1. This ratio quantifies the hardness of a sweep, with lower values indicating the presence of one dominant haplotype. (D-F) Visualization of haplotypes in the highlighted regions detected by sharing probability, analogous to Fig. S26. Regions 1 and 2 are examples of hard sweeps dominated by one haplotype; in these cases, the sharing probability metric gives comparable values to H1 and H12. Region 3 is a candidate of a very soft sweep, which is hard to detect for both H1 and H12. These results suggest that the sharing probability metric is capable of recovering signals of existing selection tests based on haplotype homozygosity, and has enhanced power for detecting soft sweeps.

**Fig S28.**
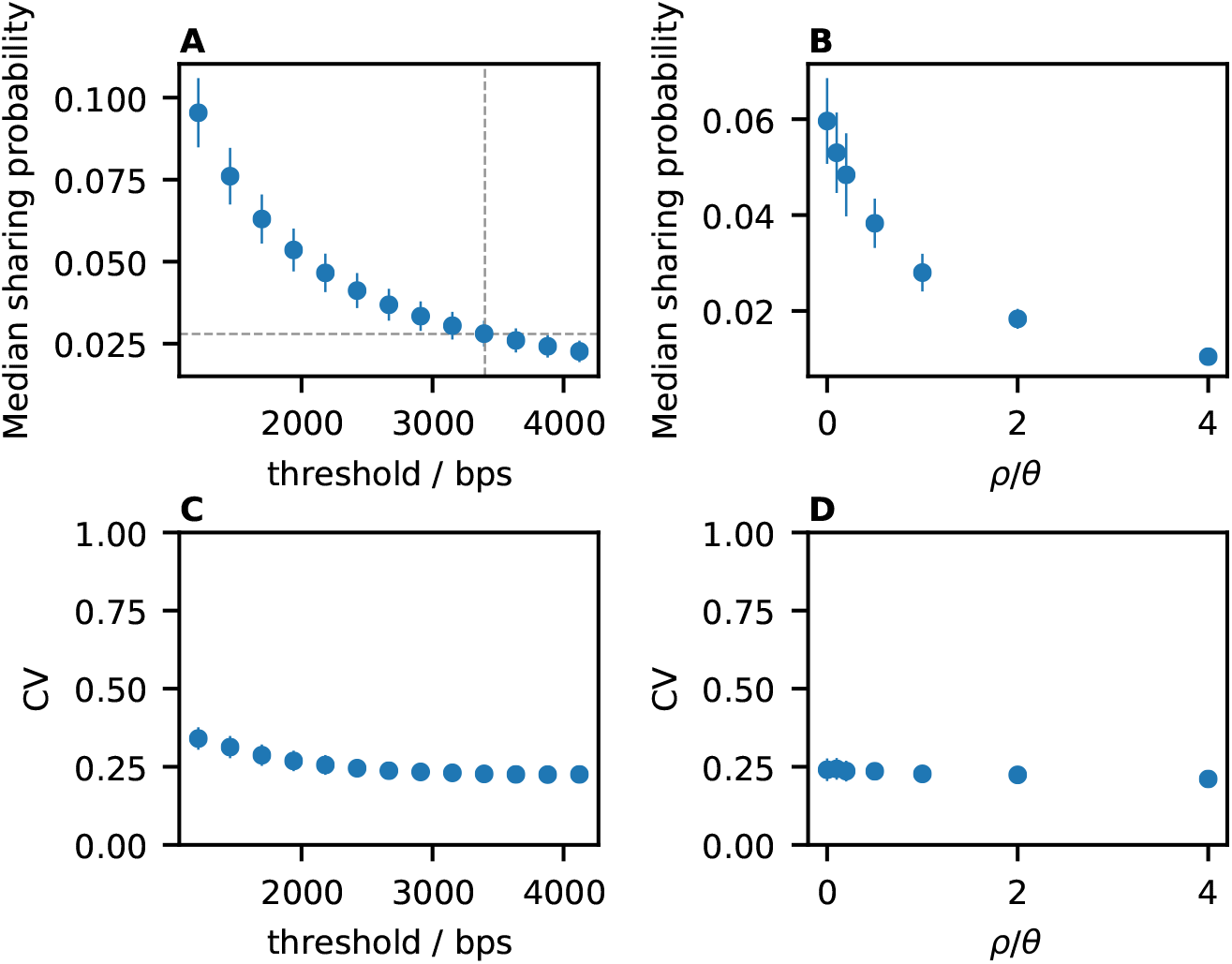
Sharing landscape statistics in neutral simulations. Data shown here are generated using FastSimBac (S1 Text 5.3). (A) Median sharing probability across all genome positions for a range of threshold lengths (*ρ*/*θ* = 1, same data as Fig. 6B). Crosshair shows the threshold length that matches the median within-clade sharing probability of *B. vulgatus*. (B) Median sharing probability for a range of recombination rates. (C,D) Coefficient of variation (CV) of the sharing landscape for each of the datapoints in panels A and B, respectively. In all panels, error bars represent one standard deviation among 100 replicates. These results show that neutral models predict small fluctuations in the sharing probability across the genome, regardless of the threshold length or *ρ*/*θ*.

**Fig S29.**
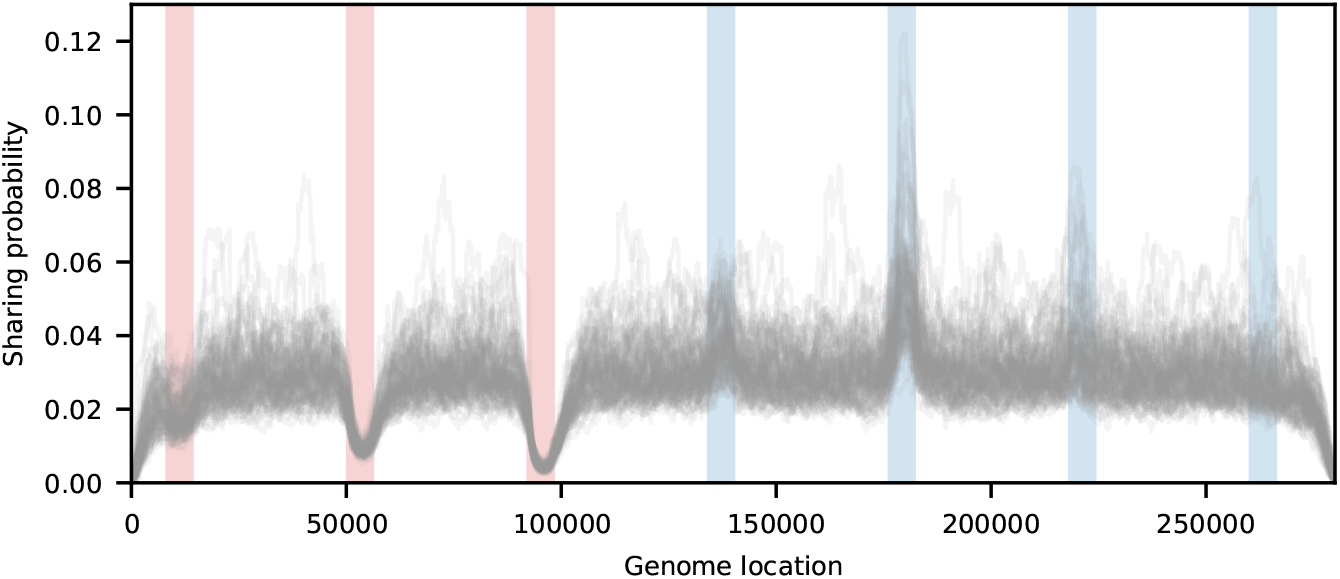
Effects of recombination rate variation on the sharing of long segments. FastSimBac simulations of a neutral population with various recombination hotspots/coldspots along the genome, highlighted in red/blue. Data shown here are 100 simulations using the same base parameters as Fig. 6B, with local modifications of the recombination rate in the highlighted regions; from left to right, the recombination rates relative the default value are {2, 5, 10, 0.5, 0.1, 0.01, 0}. These simulations show that the sharing probability decreases dramatically in recombination hotspots across all replicates. Conversely, lowering the recombination rate increases the sharing probability slightly, but not enough to reproduce sharing hotspots observed in real data. Curiously, this enhancement of sharing due to reduced recombination is non-monotonic.

**Fig S30.**
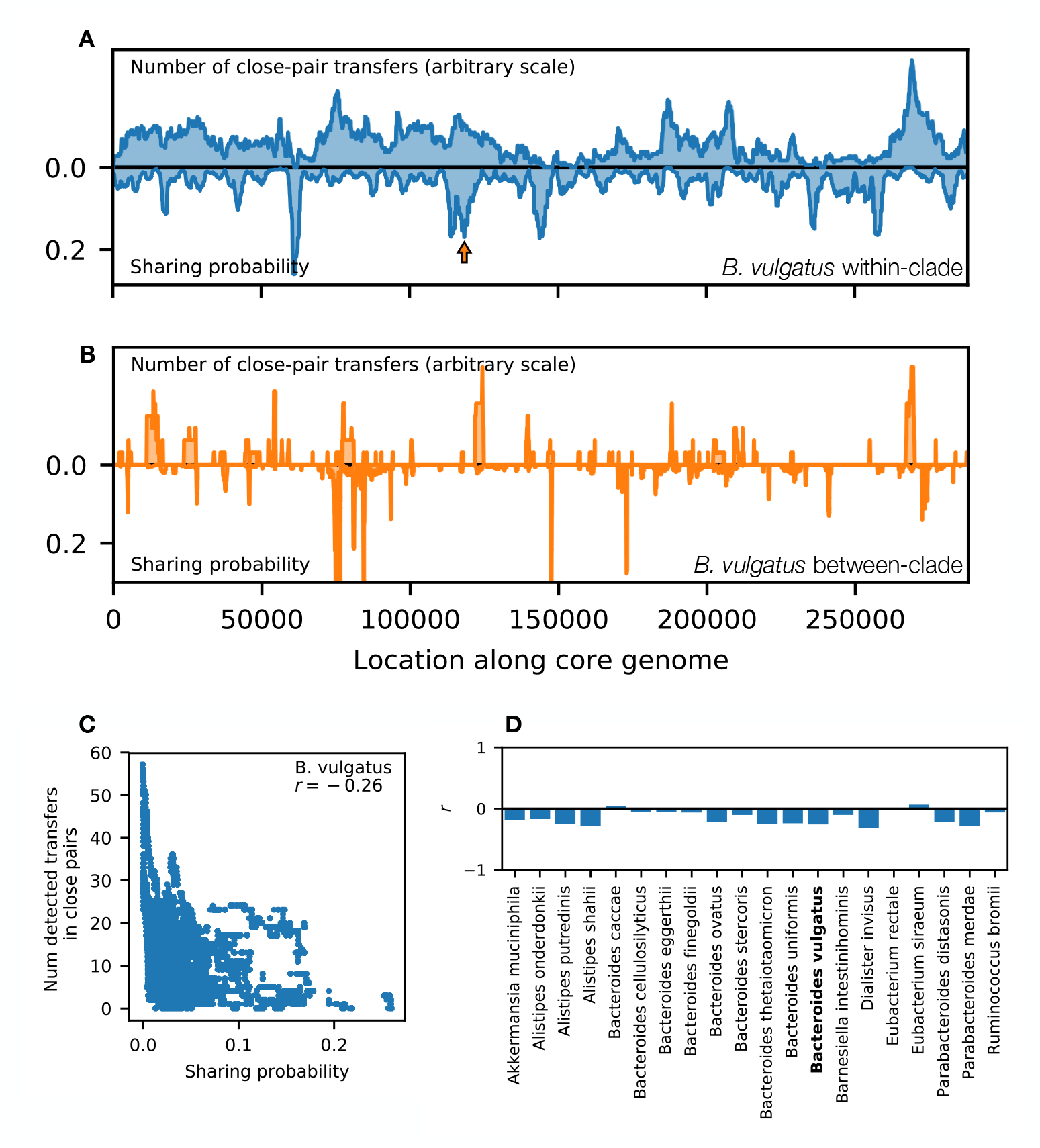
Comparing the landscapes of detected transfers and recent haplotype sharing. (A) Example landscapes computed for strains within the major clade of *Bacteroides vulgatus*. Top: the number of detected transfers between closely related pairs at each location. A deduplication step was performed to avoid overcounting transfer events (S1 Text 3.9). Bottom: same data as Fig. 6B (within-clade). Only four-fold degenerate (4D) sites along the core genome are shown. Visually, there is only limited correlation between the two landscapes. Although the highest peaks in either landscape corresponds to the lowest trough in the other, we also observe regions with high values of both haplotype sharing and detected transfers (e.g. the sweep region in Fig. 5, marked by orange arrow). (B) Analogous version of panel A for recombination between the major clades of *B. vulgatus*. (C) Scatter plot showing weak correlation between the values of two landscapes in panel A at each site along the core genome. Inset shows the Pearson correlation coefficient, *r*. (D) Correlation coefficient of landscape values for all species with sufficient data, analogous to panel B. The limited correlation in *B. vulgatus* is observed across all species, suggesting that the two metrics might reflect different aspects of the recombination dynamics.

**Fig S31.**
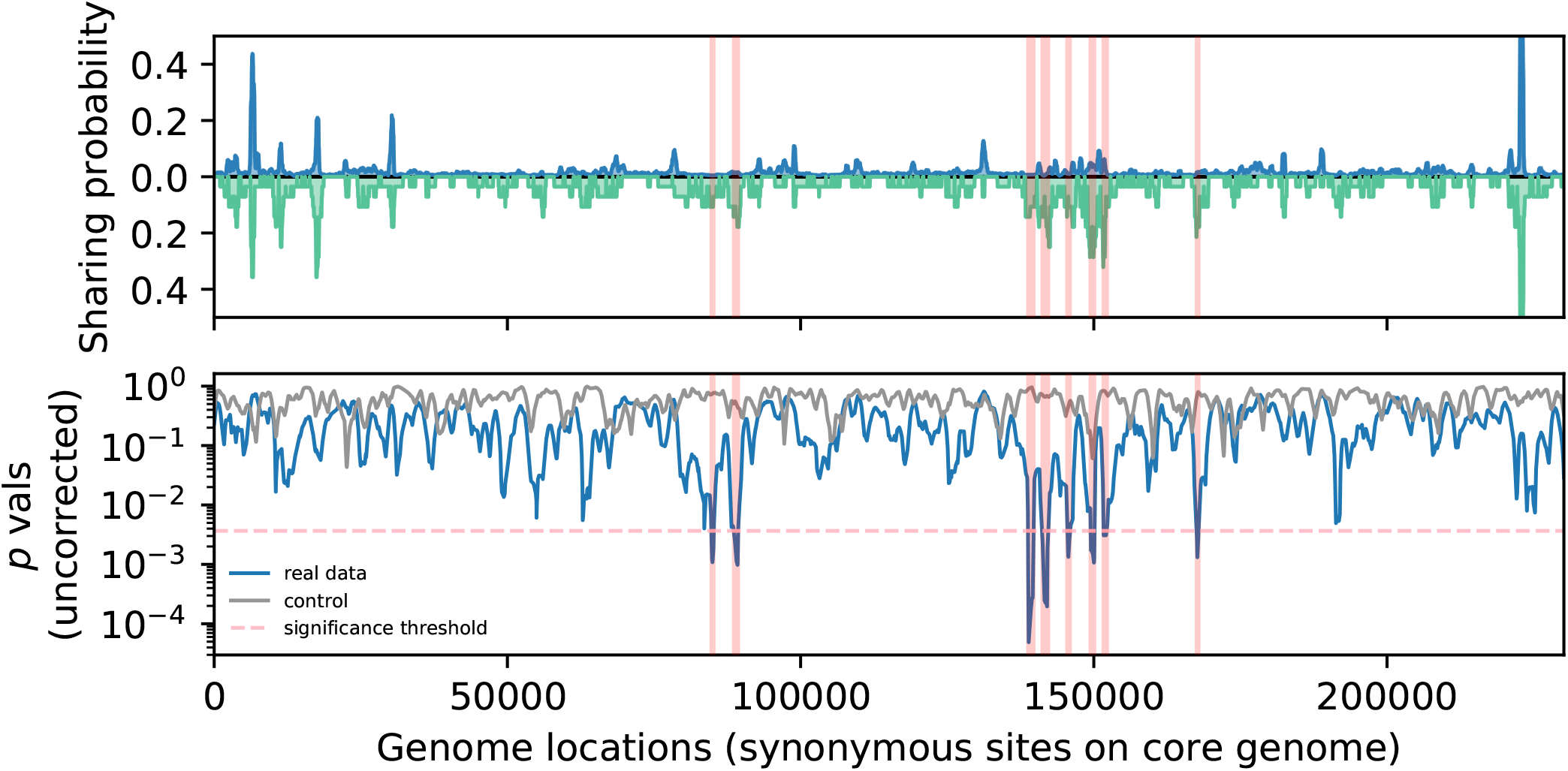
Within-host enrichment in the sharing landscape of *Eubacterium rectale*. Top panel: Analogous version of Fig. 6C, that highlights the regions that were identified to be enriched in within-host sharing (S1 Text 5.4). Bottom panel: First level *P* values from double-bootstrap permutation test (S1 Text 5.4), averaged in sliding windows of 1000 sites. Grey curve shows the result of a negative control, obtained by applying the test to an independent permutation. Pink dashed line represents the genome-wide significance level (*P* < 0.05) for first level *P* values. Regions with statistically significant differences in within-host sharing are highlighted in both panels.

**Fig S32.**
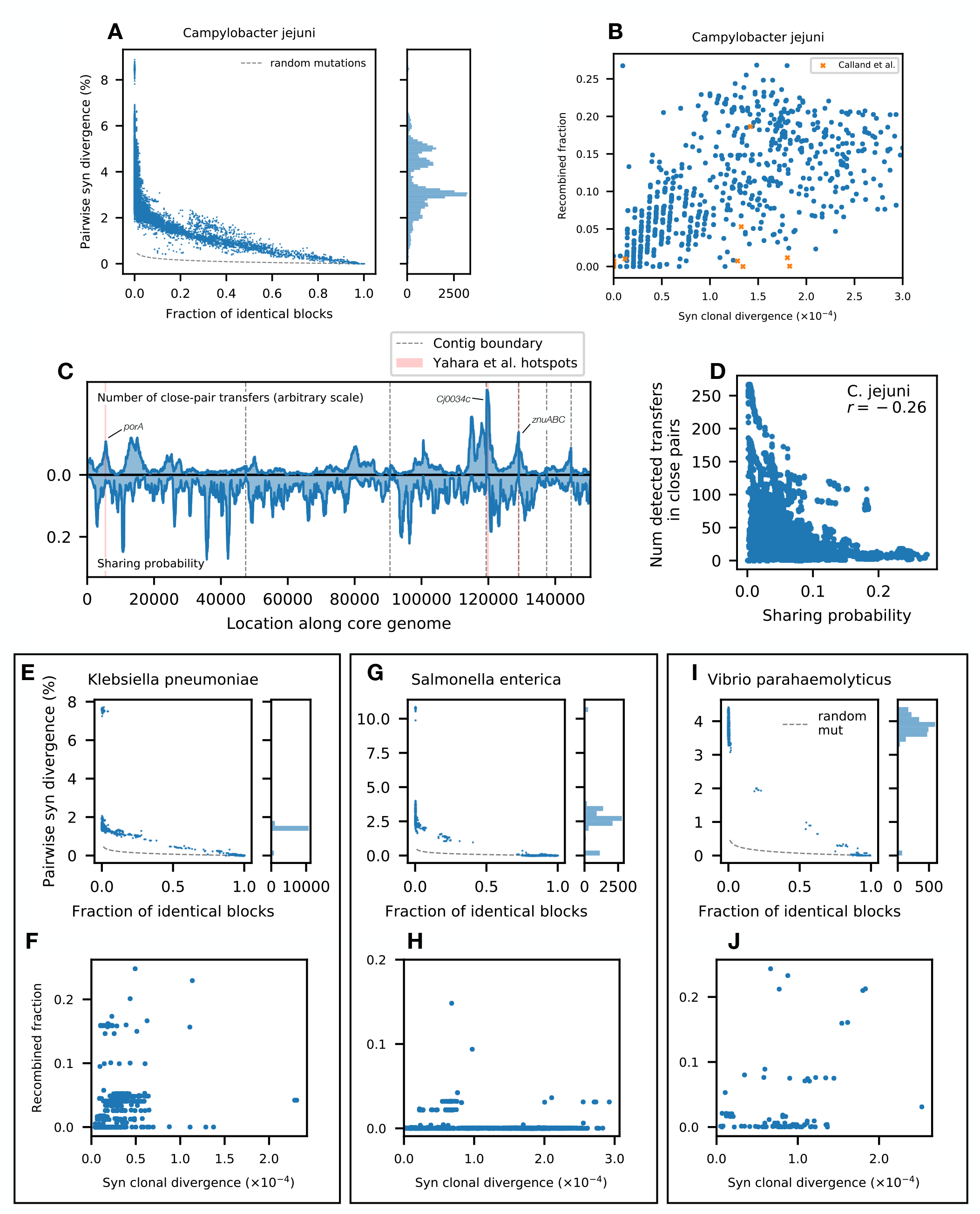
CP-HMM analysis of isolate genomes from bacterial pathogens. CP-HMM was applied to a total of 685 previously sequenced isolate genomes from 4 commonly studied bacterial pathogens (S1 Text 3.10). (A) Analogous version of Fig. 1C for *Campylobacter jejuni*, showing a large number of closely related pairs. (B) Analogous version of Fig. 3A-C. Orange crosses show the analogous data points reported in Ref. 48, where recombined regions were inferred for a smaller sample of 12 strains using Gubbins [29]. These data demonstrate that the apparent recombination rates reported in Ref. 48 are consistent with the results inferred using the CP-HMM algorithm. (C-D) Analogous versions of Fig. S30A-B. The peaks in the landscape of detected transfers reproduce the “recombination hot regions” previously reported in Ref. 73. However, the sharing landscape (C, bottom) reveals a different set of locations that are enriched for recent haplotype sharing. This suggests that these two landscapes reflect different aspects of the interplay between recombination and natural selection. (E-J) Analogous versions of A-B for three other pathogen species. It is worth noting that *Salmonella enterica* contains both a large number of pairs with minimal recombination and many pairs with significant recombination. This is consistent with previous studies showing that r/m in *S. enterica* can differ by 10-fold between sublineages [61].

**Fig S33.**
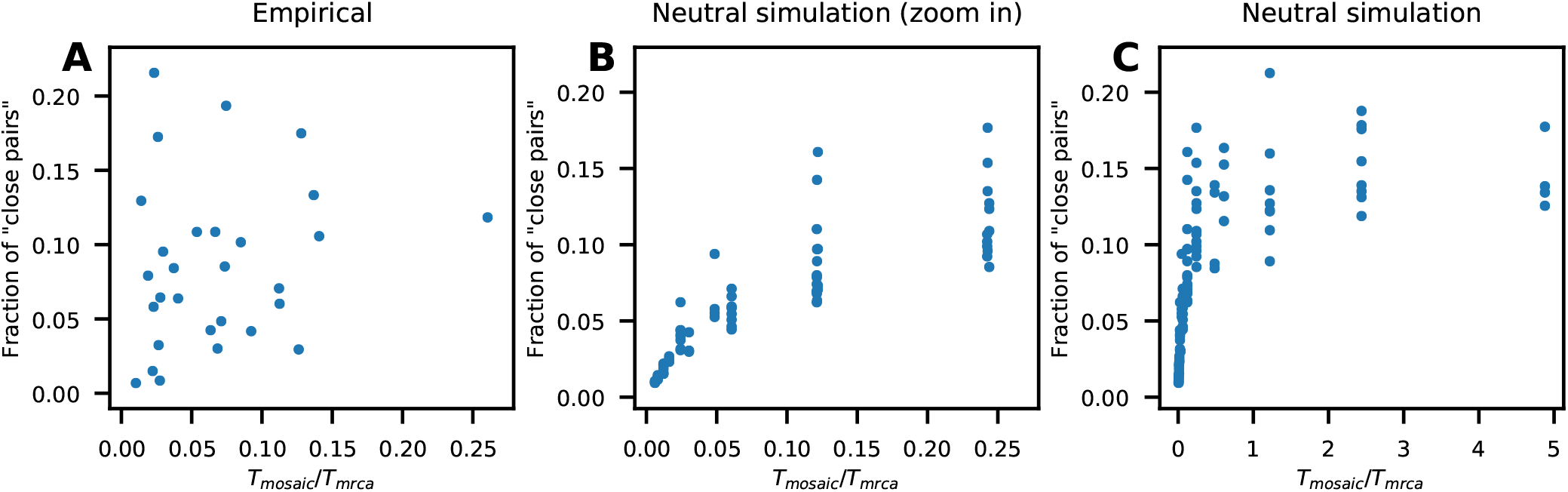
Correlation between *T*_mosaic_/*T*_mrca_ and the number of closely related pairs in different species. (A) Horizontal axis shows the average *T*_mosaic_/*T*_mrca_ estimates for each species in Fig. S16, while the vertical axis shows the fraction of genome pairs that have > 20% identical blocks (S1 Text 2). (B) Correlation between the true value of *T*_mosaic_/*T*_mrca_ and the number of closely related pairs in neutral simulations. We simulated 100 populations using FastSimBac (S1 Text 5.3) with *ρ*/*θ* = {0.1, 0.5, 1, 1.5, 2}, *θ* ≈ 0.008, *λ* = {500, 1000, 2000, 5000, 10000}, sample size 200, covering a wide range of *T*_mosaic_/*T*_mrca_ values. (C) Analogous version of panel B showing a wider range of *T*_mosaic_/*T*_mrca_ values. Unlike the neutral simulations, the observed data show little correlation between the fraction of close pairs and *T*_mosaic_/*T*_mrca_. Moreover, some species can have high fractions of close pairs at very low *T*_mosaic_/*T*_mrca_ values, in sharp contrast with the predictions of the neutral simulations.

**Fig S34.**
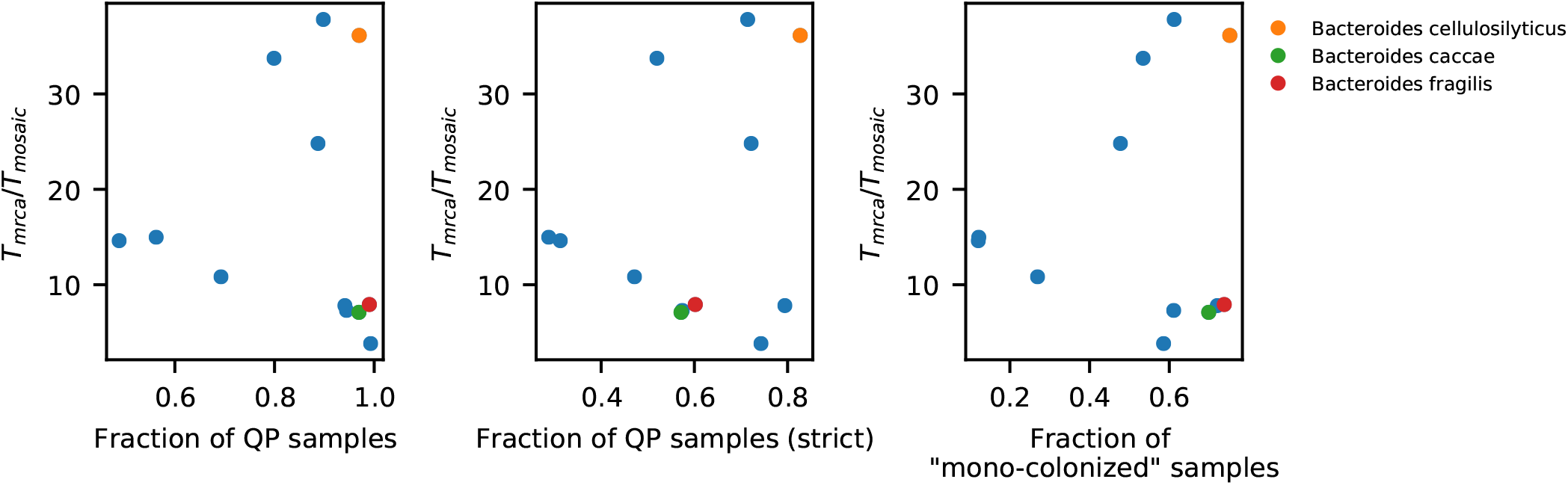
Colonization strategies are not correlated with recombination rates inferred from close pairs. Data shown here are from 12 species in the *Bacteroides* genus, with each point representing a species. In all three panels, the vertical axis shows the estimated *T*_mrca_/*T*_mosaic_ (Fig. S16). Left panel: Horizontal axis shows the fraction of “quasi-phaseable” (QP) samples for that species, which serves a proxy for the degree of single- vs multi-colonization (Fig. S1). For example, *B. fragilis*, a species known to have mostly clonal within-host colonization structure [71], has a QP fraction of > 99%. Middle panel: horizontal axis shows the fraction of QP samples under a stricter definition of quasi-phaseability – instead of defining alleles with frequencies between 0.2 and 0.8 as polymorphic [27], this stricter version expands the frequency range to between 0.05 and 0.95. Right panel: horizontal axis shows an even stricter definition of single-colonization – any site with more alternative alleles that what is expected under a Poisson error model will be considered a polymorphic site, and a sample is “single-colonized” if the polymorphism rate is lower than 0.1%. Interestingly, all three panels show little correlation between the apparent degree of multi-colonization and inferred recombination rates. In particular, species that exhibit the lowest levels of multi-colonization can have drastically different rates of recombination (e.g. *B. cellulosilyticus* vs *B. caccae*). This lack of correlation suggests that physical proximity may not be the main driver of the variation in the inferred recombination rates between species.

**Table S1. Metadata of metagenomic samples used in this study.** We analyzed a collection of 932 samples from 693 individuals, collated in a previous study [27]. This included samples from 250 individuals from the Human Microbiome Project (HMP) [94, 95], 250 individuals from Ref. 97, 185 individuals from Ref. 96, and 8 individuals from Ref. 100. Listed are the subject identifiers, sample identifiers, run accessions, country of the study, continent of the study, visit number, and study.

**Table S2. Number of close pairs across species.** This table contains statistics of closely related strains across 43 species in our cohort. For each species, we computed the fraction of identical genome blocks for all pairs of genomes from unique hosts, and recorded the number of pairs with >20%, >50%, >80% identical blocks. This table also contains the number of genomes in each species (“num_qp_samples”). Some species (e.g. *Prevotella copri*, *Roseburia inulinivorans*) have substantially fewer closely related pairs than others with comparable number of genomes.

**Table S3. Detected transfers in the closely related pairs of 29 species.** This table contains all the locations and divergences of recombination transfers shown in Figs 2 & 3. Listed are the species names, sample identifiers for each pair of strains, if the transfer is between-clade (‘Y’, ‘N’, ‘NA’), if the transfer is included in Fig. 3 (‘TRUE’, ‘FALSE’), divergences (all sites or synonymous sites only), locations of transferred regions, and if the transfer is a potential duplicate of other detected transfers (‘TRUE’, ‘FALSE’) (see S1 Text 3.9).

**Table S4. Species with high-quality dual-colonized samples.** Listed are species with >5 high-quality dual-colonized samples that passed the filters described in S1 Text 4.1.

**Table S5. Annotations for genes in the within-host sweep example of *Bacteroides vulgatus*.** Listed here are genes involved in the within-host sweep example in Fig. 5 that have within-host SNVs at the first time point. Gene annotations are downloaded from PATRIC [101].

**Table S6. Clonal divergence thresholds** *d*^∗^ **and clonal fraction thresholds** *f* ^∗^. Clonal fraction thresholds *d*^∗^ and clonal fraction thresholds *f* ^∗^ for selecting close pairs in certain species (S1 Text 3.3).

**Table S7. Metadata of isolate genomes used in S1.3.10.** Listed are the species names, species types (commensal or pathogen), genome accessions, and other information compiled in Ref. 92.

## S1 Text: Supplemental Methods

### 1 Metagenomic pipeline

We utilized a collection of metagenomic data that was collated in a previous study [27]. This collection consists of 932 fecal samples from 693 subjects from North America, Europe, and China, some of whom were sequenced at 2-3 timepoints roughly 6 months apart. We analyzed these data using the same reference-based pipeline described in Ref. 27. In short, we used the MIDAS software package [102] to align the raw sequencing reads from each sample to a large collection of reference genomes representing different bacterial species. The distribution of coverage across each genome was used to estimate the relative abundances of each species in each sample, as well as the presence and absence of individual genes. Following Ref. 27, we defined the core genome of each species to be the subset of protein coding genes that were present in >90% of the samples in which the species could be reliably detected. We chose to focus on the core genome to limit the impact of plasmids and other mobile genetic elements, which can be horizontally transmitted at much higher rates than typical chromosomal DNA. By restricting our attention to core genes, we aimed to infer the baseline rates of homologous recombination across the entire genome, which are critical for understanding the genetic structure of bacterial populations [16].

We used MIDAS’s snps module to identify single nucleotide variants (SNVs) in each species based on the raw read pileups in each sample. These initial SNVs were subsequently filtered based on their absolute and relative coverage, as well as their location along the genome, using the same procedures and parameters described in Ref. 27; to reduce the effects of reference bias, no additional filtering is done using the allele counts as this stage (see below). The end result of this pipeline is a list of coverage values (*D*_*i*_) and alternate (non-reference) allele frequencies (*f*_*i*_) at each genomic position in the reference genomes that are detected in a given sample. Filtered sites and other missing data are assigned a coverage value of *D* = 0. These lightly processed SNV frequencies served as the basis for all of our downstream analysis.

The frequencies of the SNVs within each sample provide information about the lineage structure of the within-host population (Fig. S1). Ref. 27 showed that the lineage structure in many samples is sufficiently simple that the genotype of the dominant lineage can be inferred with a high degree of confidence. We used this “quasi-phasing” approach to infer the genotypes of 5416 strains from 43 different species, using the same procedures and parameters described in Ref. 27. These quasi-phased genotypes served as the basis for all of our subsequent between-host comparisons. Importantly, our downstream analyses only consider sites with high quality genotype calls, regardless of their polarization. Treating the remaining sites as missing data (rather than reference alleles) ensures that our between-host comparisons do not sensitively depend on the choice of the reference genome used for read mapping.

In S1 Text 4, we describe the additional methods we developed for analyzing non-quasi-phaseable samples to quantify recent recombination events within hosts.

### 2 Identifying partially recombined genomes from pairwise diversity statistics

To test whether the local recombination model in Fig. 1A could explain the broad range of diversity in many gut species, we used a lightweight approach similar to Refs. [19, 28], which compares the joint distributions of two diversity statistics that can be directly estimated from pairwise comparisons between quasi-phaseable samples.

For each species, we first filtered the data to retain at most one quasi-phased strain per household. We then considered all pairwise comparisons between the strains in these unrelated hosts. For each pair of strains, we identified the subset of four-fold degenerate (4D) synonymous sites in the core genome that were covered in both samples. We used this subset of sites to compute the average divergence across the genome (*d*) and the fraction of multi-site “blocks” that contained zero SNVs (*f*). Based on the divergence scales in our data, we chose a block size of *l* = 1000 4D sites, so that a typical gut bacterial genome contains 200-400 such blocks.

We compared these data to a null model in which SNVs accumulated independently and uniformly along the genome. This would be the case, e.g., in an asexual model with a uniform mutation rate along the genome. In this case, the independent accumulation of SNVs leads to a simple relationship between the average divergence (*d*) and expected fraction of identical blocks (*f*). The probability that a block of length *l* contains no SNVs is given by

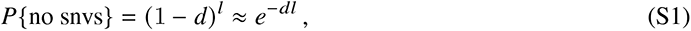

where we have used the fact that *d* ≪ 1 and *l* ≫ 1. Since an individual genome usually contains many blocks, we can equate *P*{no snps} with the expected fraction of identical blocks *f*, yielding

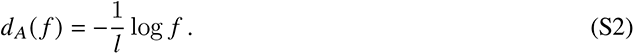

This expression was used to plot the dashed lines in Fig. 1B, S3, S2.

Deviations from this null expectation can naturally arise in simple models of bacterial recombination (Fig. 1A). For example, in the simplest model where we assume that each block is replaced by a recombination event at a constant rate *R* per generation, the probability that a block has not been modified by recombination is given by

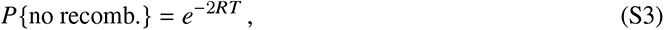

where *T* is the coalescence time of the given pair of strains. Similarly, the probability that a block has not been modified by mutation is given by

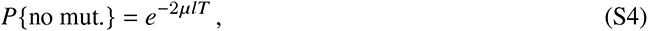

where *μ* is the mutation rate per site per generation. Then, the expected fraction of identical blocks is given by *f* = *P*{no recomb.}· *P*{no mut.}, while the fraction of recombined blocks is given by *f*_*r*_ = 1 − *P*{no recomb.}. <u>If</u> we additionally assume that the recombination events import segments of the same characteristic divergence, *d* (corresponding to the average pairwise divergence within the species), then a typical transfer event will introduce many more SNVs than mutation in the clonal regions. This suggests that we can approximate the genome-wide divergence *d* by the recombined regions alone:

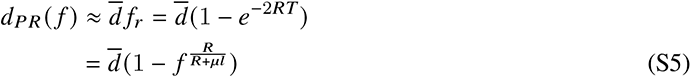

This provides a simple a<u>lt</u>ernative model connecting the fraction of clonal blocks with the total genome-wide divergence. In practice, *d* can be measured directly from the average pairwise diversity of the species, while the compound parameter 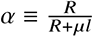 can be fit using the estimator

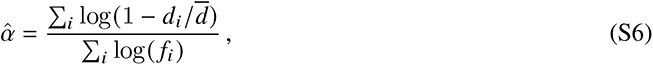

where *d*_*i*_ and *f*_*i*_ are the observed values for each pair that has nonzero *f*.

To examine the usefulness of these expressions, we simulated well-mixed, neutral populations with mutation and localized recombination (5.3), and calculated the same pairwise statistics across a range of parameter values (Fig. S2). We found that for small recombination rates or short transferred fragments, the joint distribution of *d* and *f* is well-described by Eq. (S2), while increasing either parameter will lead to larger deviations from this null model of random mutations. In simulations with higher recombination rates, the <u>m</u>ajority of pairs have a small fraction of identical blocks and cluster around the average pairwise divergence *d* (Fig. S2). We observed similar patterns in most of the gut bacterial species in our cohort, consistent with the picture of “quasi-sexual” species in Fig. 1. For these parameters, we found that the recombination model in Eq. (S5) accurately captures the results of the simulations, with *α* reflecting the relative strength of recombination vs mutation. These results suggests that our theoretical expressions Eqs. (S2) and (S5) can be used in combination to determine whether the pairwise diversity of a population is better described by an asexual model or a partial recombination model.

Consistent with our simulations, we found that the smallest divergence values in many species were well-predicted by their corresponding fraction of identical blocks. To quantify how well this additional variable explained the variation in pairwise divergence within species, we fit the joint distribution of pairwise statistics with a modified version of Eq. (S5),

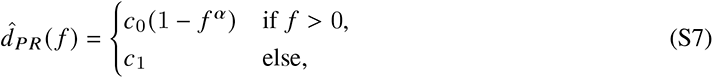

where *c*_1_ is the average divergence across all pairs with *f* = 0, and *c*_0_ is the average divergence across all *within-clade* pairs with *f* = 0. The reason we treated the *f* = 0 case separately is that some species exhibit significant variation in pairwise divergence at *f* = 0 due to population structure, which is not included in the simple model above. After fitting the partial recombination model in Eq. (S7), we estimated the goodness of fit (*R*^2^) using the standard formula,

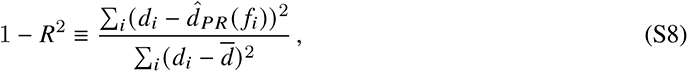

where *d*_*i*_ and *f*_*i*_ are the observed values for each pair, and *d* is the average divergence across all pairs.

We found that for some species (e.g. *Eubacterium rectale*), the large range of variation among close pairs is well explained by partial recombination model (Fig. S3), but *R*^2^ is sometimes overwhelmed by the large number of points at high divergence values (whose variation arises from population structure rather than the amount of clonal inheritance). To better capture the power of this variable in explaining the *range* of divergences, we also computed a weighted goodness-of-fit statistic, *R*_*Y*_^2^, that gives each range of *y* values an equal weight. More specifically, we divide the entire *y* range into 500 bins (*B*_*i*_), count the number of points in each *y* bin (*n*_*b*_), and use the 1/*n*_*b*_ as the weight of each data point:

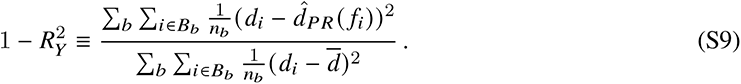

We found that this modified metric successfully captures the explanatory power of *f* for species like *E. rectale*. For most species, however, *R*^2^ is approximately the same as *R*^2^.

Finally, the pairwise divergence distribution revealed that some species (e.g. *Bacteroides vulgatus*) had a strong population structure of two or more clearly separable clades (Fig. S3). For these species, we used the pairwise divergence matrix to cluster the strains into two major clades using the linkage and fcluster functions in the SciPy hierarchical clustering library [103]. We then computed analogous versions of *R*^2^ and *R*^2^ that were restricted to strains from the largest clade (Fig. 1). Again, we found that explicitly removing the population structure in this way was essential for revealing the explanatory power of *f* in cases where our naive metrics would be overwhelmed by the large divergences that have accumulated between the major clades.

### 3 Identifying individual recombination events from pairs of closely related strains

To compare recombination dynamics across different species of gut bacteria, we developed a hidden Markov model method (CP-HMM) to automatically identify recombination events from the spatial divergence profiles of closely related strains (Fig. 2B). This HMM approach is conceptually similar to the one employed in Ref. 28, but with important technical modifications that we describe in more detail below. These pairwise HMMs can be viewed as a lightweight version of existing phylogenetic approaches like ClonalFrameML [31] or Gubbins [29]. Since the genealogies of closely related pairs are particularly simple (Fig. 1B), this pairwise approach can implicitly capture various forms of selection, non-equilibrium demography, and other deviations from the simple neutral models assumed in previous work, even when there is insufficient data for a complete phylogenetic reconstruction. This additional flexibility will be important for our analysis of the gut bacterial species below.

#### 3.1 Description of the CP-HMM method

CP-HMM is designed to model the spatial divergence profiles observed between pairs of closely related strains. As in Fig. 1, we focus on the subset of 4D synonymous sites in the core genome that are covered in both samples. To minimize the effects of correlated mutations (e.g. due to mapping artifacts or multi-nucleotide mutations [104]), we first coarse-grain the genome into blocks of *b* = 10 4D sites and assign a binary label to each block (0 =“no SNVs in block”, 1=“one or more SNVs in block”). Since all species in our study have a typical heterozygosity less than 10%, we expect that normal regions of the genome will usually have fewer than one SNV per block, and thus will be minimally impacted by this coarse-graining scheme.

We assume that these coarse-grained divergence profiles can be described by a hidden Markov model with two classes of hidden states: a clonal state *C* and a finite number of recombined states {*R*_*i*_}, each of which is associated with a corresponding pair of parameters, (*λ*_*i*_, *θ*_*i*_), representing the average length and average divergence of the recombined fragment. The transitions between the hidden states are governed by the sparse transition matrix:

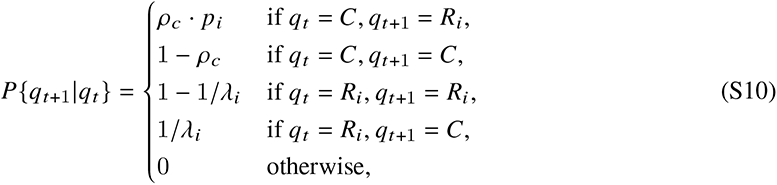

where *ρ*_*c*_ is the overall probability of transitioning to a recombined segment from a clonal region, and *p*_*i*_is the probability that the recombined segment will be of type *R*_*i*_ (Fig. S5). In cases where the reference genome of the species contains multiple contigs, we assume that the hidden states in each contig are obtained from independent draws from the same transition matrix in Eq. (S10). Finally, given a sequence of hidden states {*q*_*t*_ }, the corresponding emission probabilities are

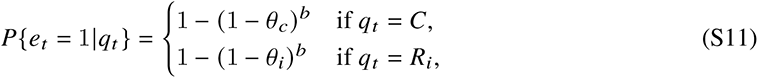

where *θ*_*c*_ can be interpreted as the average divergence within the clonal region.

These assumptions define a hidden Markov model for the coarse-grained divergence bins as a function of the underlying parameters *ρ*_*c*_, *θ*_*c*_, and {*p*_*i*_, *λ*_*i*_, *θ*_*i*_}. In practice, we found that the exact numerical values of these parameters did not affect our main findings as long as they had the correct order of magnitude. Since different pairs of strains have diverged from each other for different amounts of time, the *ρ*_*c*_ and *θ*_*c*_parameters will generally vary widely across strains, and must be inferred for each pair using the iterative approach described below. For simplicity, we assume that the remaining parameters, {*p*_*i*_, *λ*_*i*_, *θ*_*i*_}, are shared by all strains from the same species. We fixed these parameters using the following considerations.

We assumed that the average lengths were similar for all recombined states (*λ*_*i*_ ≈ *λ*), with a few exceptions described below. Since this length scale is initially unknown, we developed an iterative approach to infer the relevant value of *λ* for each species in a self-consistent manner. Starting from an initial guess of *λ* ≈ 1000, we ran the CP-HMM algorithm described below to identify an initial collection of recombination events from the data. We then updated *λ* based on the average length of the detected transfers in the first iteration. We repeated this process several times for each species until the *λ* estimates converged to a steady value. In practice, we found that these estimates converged relatively rapidly (< 10% change after two iterations).

The remaining parameters, {*p*_*i*_, *θ*_*i*_}, can be viewed as a discrete approximation to the underlying probabilities of importing fragments with different levels of divergence. Given our large sample sizes, we assumed that this distribution could be approximated by the *empirical distribution* of local divergence values observed in random pairs of strains from our larger sample. Specifically, we divided the core genome into blocks of Δℓ = 1000 4D sites, and calculated the frequency of local divergence values (in 40 equally spaced bins) across all pairs of strains in our sample. We note that these empirical distributions of local divergence are much broader than a Poisson distribution with the same mean (e.g. Fig. S6), which would be the null expectation assuming a constant local coalescence time. This discrepancy illustrates the complex genealogical history along a bacterial genome, as well as the necessity of using an empirically-derived estimate of *p* (*θ*). For species with a strong population structure, such as *Bacteroides vulgatus*, *p* (*θ*) naturally separates into distinct distributions when conditioned on sampling pairs from the same clade or different clades (Fig. S6). In these cases, we utilized this clean separation to further divide the set of recombined states into two types, representing within-clade and between-clade recombination events, whose corresponding values of (*p*_*i*_, *θ*_*i*_) are given by one of the two empirical histograms. We also allowed the length scales *λ*_*i*_ to vary for within-clade vs between-clade events, and used the same iterative approach described above to estimate these parameters for each clade.

With the above information, the hidden Markov model is fully specified. For each pair of strains, CP-HMM infers a sequence of hidden states {*q̂*_*t*_ } and model parameters (*ρ̂*_*c*_, *θ̂*_*c*_) using standard HMM algorithms [105]. In principle, the *ρ*_*c*_ and *θ*_*c*_ parameters are related to the per-base pair recombination and mutation rates for the pair of strains, respectively. However, for reasons described below, we did not work with these global parameters directly, but instead sought to identify individual recombination events and clonal divergence values directly from the sequence of {*q̂*_*t*_, *q̂*_*t*_ } values.

For example, we identified individual recombination events from this sequence by computing all continuous stretches of recombined states. To eliminate potential mapping artifacts, we excluded all events containing ≤ 50 synonymous sites, which are much shorter than the typical recombined segments (Fig. 3). Such events could originate from misaligned indels that generate enough SNVs that would cause them to be mis-identified as short recombined segments. For each of the recombination events that passed this filter, we estimated the local synonymous divergence of the recombined fragment based on the total number of SNVs that it contains (Figs 4 & S18).

Similarly, we defined the “clonal fraction” of the genome to be the total fraction of sites with *q̂*_*t*_ = *C*. Since undetected recombination events can massively inflate the number of SNVs in the clonal regions, we estimated the clonal divergence using an approach that filters out these regions with anomalously high local SNV density. Specifically, we coarse-grained the clonal regions into blocks of 1000 synonymous sites, and computed the average clonal divergence using only those blocks with ≤ 2 SNVs. We found that this procedure greatly improved the accuracy of estimated clonal divergence in our validation analysis below.

##### 3.1.1 Relation to previous work

Our CP-HMM model is most similar to the HMM used in the pairwise analysis in Ref. 28, but with a few important distinctions. First, our model utilizes a range of recombined states as opposed to a single one. A negative binomial distribution is used in Ref. 28 to account for the fact that recombined regions consist of mosaics of Poisson distributions with different rates. However, for species like *B. vulgatus*, a negative binomial distribution cannot capture the underlying bimodal distribution caused by population structure. We sought to overcome this problem by explicitly introducing a large number of recombined states, each of which corresponds to a different (Poisson) divergence rate. We took advantage of our large data set to fit these rates empirically from the non-closely related pairs as described above. This non-parametric approach provides a more flexible route for inferring the underlying divergence distribution of the transferred segments (Fig. S18), and can be naturally generalized to account for clades and other forms of population structure (Figs 2 & S17). Second, instead of coarse-graining the alignment into large (a few kb) blocks, our model can achieve nearly site-level classification. This level of resolution is important for estimating not only transfer lengths, but also clonal divergence times. In fact, there will be an artificial bias had we used much larger blocks. Consider a long transfer spanning multiple blocks, where blocks on the two ends are only partially covered by the transfer. The divergence of the two end blocks will typically be lower than the mean divergence in the transfer. These end blocks will have some probability of being classified as clonal sequence, which will drive up the estimated clonal divergence. Since every transfer is susceptible to this bias, the clonal divergence estimation can be artificially correlated with the number of detected transfers. The relationship between the number of transfers and the clonal divergence – a central piece of our analysis – can therefore appear to be artificially linear, masking the high degree of variation that we actually observed. This subtle mode of error necessitates our more fine-grained approach.

#### 3.2 Validation on simulated data

To validate our algorithm’s performance, we designed a simple computational model to simulate mutations and recombination transfers accumulated between a pair of closely related strains. Similar to [39], our model simulates the spatial distribution of SNVs along the genome. To better match the format of the real data, we simulate only the synonymous sites on the core genome. Let *L* be the number of sites and *T*_*div*_ be the divergence time between the pair. As a simple model of recombination, we assumed a constant mutation rate *μ* and recombination rate *r* at each site along the genome, so that the total number of mutations and transfers will follow Poisson distributions with means 2*μT*_*div*_ *L* and 2*rT*_*div*_ *L*, respectively.

After drawing the total number of mutations and transfers to introduce, we first introduced the mutations randomly along the genome, and then introduced the transfers. Since real SNV densities can vary along the genome due to local variation of mutation rates, selective pressure and recombination history, preserving this variation is crucial to evaluating the performance of the HMM. To account for such patterns, we sampled the sequence of the recombinated fragment from one of the real genomes in our cohort. More specifically, we randomly chose one strain at the beginning of the simulation to serve as the fixed focal strain. Next, for each recombination event, a random other strain is chosen as the donor, a random site is chosen as the starting point, and a transfer length is drawn from an Exponential distribution with mean *λ*. The corresponding sequence of the donor strain will then overwrite the focal genome, introducing additional SNVs alongside the mutation events above. For simplicity, we do not model a circular genome, so a transfer stretching beyond the alignment’s end will be truncated. Transfers and mutations are allowed to overlap.

Using the genomes of *B. vulgatus*, we first generated 256 simulated pairs in total for a range of 16 different values of *T*_*div*_. To closely mimic the patterns of real data, we matched the recombination length and overall recombination rate to the inferred values from the within-clade pairs of *B. vulgatus* (which corresponds to *λ* = 2600 and *r*/*μ* = 0.65). We applied our CP-HMM method to this simulated data and recorded the statistics of detected transfers and clonal divergences. In Fig. S7 we compared our algorithm’s results with ground truth. Panel (A-C) plot the CP-HMM results for the inferred clonal divergence, the number of detected transfers, and the inferred clonal fraction, all of which appear to be in excellent agreement with the ground truth. We also pooled the lengths of all detected transfers and verified that the distribution follows the input exponential distribution almost exactly (Fig. S7D).

The slight under-detection of recombination events at high divergence times (i.e., in more recombined genomes) can arise from two sources: (i) extremely closely related transfers that have few mutations and (ii) merging of overlapping transfers. Both sources will be become more significant for more diverged pairs as the number of sampled transfers increases. While our current divergence-based approach cannot detect the first class of events, it is possible that variation in the accessory genome could provide further insight into these close transfers; we leave such an analysis for future work. The second class of events can be alleviated by filtering highly diverged pairs. Any residual merged transfers will bias the distribution of transfer lengths. However, the close agreement in Fig. S7D suggests that the second issue is not significant over the range of divergence times we have considered. Based on these validation results, we decided to restrict our downstream analysis to pairs of strains in which the clonal fraction was ≳ 75% (see below). This restriction ensures that the CP-HMM algorithm should have similar detection efficiencies across species, even when the species have significantly different levels of average pairwise divergence.

By using the empirical distribution of local divergence as a prior, we have implicitly assumed that the donor individuals are uniformly sampled from the (sequenced) population. To evaluate whether our results were sensitive to this assumption, we artificially increased the prior probability for between-clade transfers in *Bacteroides vulgatus* by 5 fold and reran the CP-HMM algorithm. We found that the signal in the data was sufficiently strong to overwhelm this change in prior, yielding quantitatively similar results compared to what we observed before. In particular, the median transfer lengths differed by only ≈ 10%, and we still detected ∼ 5 times more within-clade transfers than between-clade transfers. Some within-clade transfers are no longer detected because of the lowered prior probability. Our method for estimating clonal divergence has taken this potential under-detection into account, and the resulting *T*_mrca_/*T*_mosaic_ estimates only decreased by ≈ 10%. This result indicates that CP-HMM’s performance does not strongly rely on the strict adherence to the uniform sampling assumption.

#### 3.3 Application to data from the gut microbiome

For each species, we applied our CP-HMM model to all pairs of strains where the fraction of identical blocks was > 50%. We then refined this pool to focus on a subset of pairs where the clonal fraction was sufficiently high. This second step accounts for the fact that the clonal fraction is not always equivalent to the fraction of identical blocks, because the clonal region still allows mutations to occur at rate *θ*_*c*_. To obtain the relevant subset of pairs, we used the scatterplots of clonal fraction vs clonal divergence in each species (Figs S11 & S15) to manually identify a critical divergence value *d*^∗^, above which the typical clonal fraction first dropped below *f* ^∗^ ≈ 75 − 85%. We then excluded all pairs of strains in which *d* or *f*_*c*_ exceeded these critical values.

The *d*^∗^ and *f* ^∗^ values for each species are illustrated in Figs S11 & S15 and are listed in Table S6. The inferred recombination events and clonal divergence values the remaining pairs were used to create the scatter plots in Figures 2, 3, and S14.

Due to the discreteness in the number of mutation and recombination events, these scatter plots often have many overlapping points. To better visualize the underyling trend in data, we computed trend lines for each species using LOWESS (locally weighted scatterplot smoothing), a common local regression technique [106]. We chose an implementation provided by the Python package statsmodels [107], using the tri-cube weight function and 1/3 as the fraction of the data used for each estimation.

To quantify the local spread around these trend lines, we developed a procedure for computing quantiles of weighted residuals. For any given x value, we found *k* nearest neighboring points and computed their weights given by the tri-cube weight function. After sorting the points by the their absolute residuals, we obtained the normalized cumulative weight as a function of the residual. Then, we inverted the function numerically to obtain the residual sizes corresponding to desired quantiles. In the scatterplots, we chose *k* to be 1/5 of the total number of points and plotted the residuals corresponding to 66% quantile.

##### 3.3.1 Evaluating the effects of different sample sizes

To evaluate how the sample size affects our CP-HMM results, we performed a series of downsampling experiments with the *A. putredinis* dataset (as low as 1/10 of the original size), and examined how the inferred transfer length and recombination strength (*T*_mrca_/*T*_mosaic_) varies with the number of metagenomic samples (Fig. S10). While the variance among bootstrap replicates increases slightly as the dataset size decreases, the inferred parameters remain very close to the values obtained for the full dataset.

However, while the raw size of the dataset seems to play a relatively minor role for this range of parameters, the uniformity of the sample (i.e. how well it represents the broader population of a given species) can be much more important. For example, skewed sampling can lead to technical issues: *Lachnospiraceae bacterium* has no fully recombined pairs (Fig. S3), so our CP-HMM algorithm cannot obtain a prior for the divergence of the transferred fragments, and has trouble identifying potential transfer events. Skewed sampling can also overrepresent a specific set of closely related strains, e.g. when sampling clinical outbreaks in bacterial pathogens [21, 36, 46] (see our analysis of *Salmonella enterica* in S1 Text 3.10). In these cases, the inferred recombination parameters can be biased by the recent evolutionary history of the clonal bloom, rather than the long-term dynamics of the broader species.

#### 3.4 Correspondence between the number of transfers and the fraction of recombined genome

The crucial assumption that individual transfers do not overlap breaks down for more diverged pairs. These overlapping transfers are difficult to resolve into individual events, causing the number of transfers to be less representative of the overall rate of recombination. However, since overlapping transfers contain many more SNVs than the clonal regions, the non-clonal regions can still be reliably inferred by our model. This suggests that the *total fraction* of recombined regions can be used as a complementary metric to test if our previously observed patterns are robust to the effect of overlapping transfers.

Figure S11 shows an example of this metric for *B. vulgatus*, which complements the number of transfers shown in Fig. 2. Similar to Fig. 2C, we observed an overall trend of higher recombined fraction at longer divergence times. However, we once again observed pairs with large amounts of recombination at low divergence, and small amounts of recombination at high divergence. In addition, we saw that the vast majority of recombined regions are due to within-clade transfers, reflecting both the higher rates and longer transfer lengths we previously observed. This metric therefore confirmed the robustness of results reported in the main text. Similar results are observed when we apply this recombined fraction metric to other species with sufficient number of close pairs (Fig. S15).

This alternative metric also allowed us to ask if recombined regions tend to consist of many transfers of typical lengths, or a smaller number of anomalously long transfers. To distinguish these scenarios, we considered a simple model where the lengths of each transfer were drawn from the same exponential distribution, with a mean length given by our previous analysis (Fig. 2D). We used this model to compute a 95% confidence interval for the fraction of recombined genome for different numbers of transferred fragments.

By comparing the observed data with this null distribution, we identified many pairs of *B. vulgatus* strains where a small number of transfers covered an anomalously large fraction of the genome (Fig. S11).Further examination of these outlier pairs revealed extremely long recombination events that are hard to explain with this simple exponential model (Fig. S12). Long recombination events with similar length scales have previously been observed for several other bacterial species [39, 108, 109]. Intriguingly, some of the longest recombination events in our cohort appear to have multiple peaks in their associated divergence profiles, suggesting that they might actually be composed of several smaller transferred fragments. However, these non-contiguous transfers would still have to be strongly correlated with each other to cluster within the same localized regions, suggesting that they are best regarded as a single compound recombination event. We also observed long homozygous regions in a pair of *B. vulgatus* isolate genomes sampled from the same person (S1 Text 3.10, Fig. S13). This pattern is most likely generated by a few very long within-host transfer events, rather than correlated transfers that have compounded over multiple hosts. Understanding the mechanisms behind these long transfers is an interesting avenue for future work.

#### 3.5 Comparing the recombination dynamics of *B. vulgatus* and *A. putredinis* to simulated data

##### B. vulgatus simulations

To evaluate whether the recombination dynamics in *B. vulgatus* (Fig. 2) can be explained by different clade sizes, counting noise, or other potential detection biases in the CP-HMM algorithm, we repeated our analysis using the simulated data from S1 Text 3.2 (Fig. S8). Although we observed some degree of variation in the number of transfers due to counting noise, the simulated relationship between the number of transfers and clonal divergence was mostly linear. In contrast, the observed *B. vulgatus* data show a marked excess of pairs that accumulated a large number of transfers at low clonal divergence. This shows that the heterogeneities observed in the data cannot be explained by counting noise alone. We also find that the simulated data show few differences between within- and between-clade transfers when their underlying recombination parameters are the same Fig. 2. This indicates that the dramatic difference we observed in the *B. vulgatus* data cannot be explained by differences in the relative clade sizes, or a detection bias for transfers with different levels of divergence.

##### A. putredinis simulations

To test how well a single underlying recombination rate can capture the broad variation in the realized transfer frequency in *A. putredinis* (Fig. 3C), we again leveraged the computational model we developed for validating our HMM. We set the genome length and recombination length to be identical to those of *A. putredinis* and simulated three recombination rates *r*/*μ* = 5.2, 3.4, 2.1, which roughly span the range of realized recombination frequency in data. Since we need to compare the size of variation at different divergence times with real data, it is crucial to control how the number of data points changes over divergence. For example, the apparently large variation at small clonal divergences in Fig. 3C might be caused by a higher density of points. To account for this effect, we fitted an underlying distribution of divergence times to the distribution of observed clonal divergences in *A. putredinis*. We assumed that the true divergence time for each pair was a hidden random variable, which could take one of 20 discrete values covering the whole range of observed divergences. This hidden variable parameterized a corresponding Poisson random variable for the number of clonal mutations. The probability distribution for the hidden variables can be inferred from the observed distribution of clonal mutations in data, using the standard expectation–maximization (EM) algorithm [105]. From this inferred distribution, we sampled the same total number of pairs as in *A. putredinis* and proceeded with the rest of the simulation steps.

The result of applying our CP-HMM algorithm to this set of simulations is shown in Fig. S9. We found that no single recombination rate can capture the full range of variation in *A. putredinis*. High recombination rates only reproduced the points in the upper-left corner (corresponding to pairs that accumulate a large number of transfers at short divergence times), while lacking the points in the lower-right corner. The low recombination rate simulation produces the opposite pattern. This indicates that a combination of low and high recombination rates are needed to explain the pattern observed in *A. putredinis*.

#### 3.6 Estimating the divergence distribution of donor and recipient DNA sequences

One of the most common explanations for the reduced gene flow between diverged clades (e.g. Fig. 2) is that the recombination rate is reduced for DNA segments with higher sequence divergence [17]. To test if this divergence dependent recombination rate plays a role for the gut bacteria we have analyzed here, we computed the distribution of donor-recipient divergence of all detected transfers within each of the species in Fig. 3. For comparison, we obtained a null expectation of this distribution by simulating transfers with our collection of observed genomes, similar to our approach in S1 Text 3.2. If the effect of sequence divergence was significant, we would expect to observe a depletion of transfers with higher divergence (or an excess of transfers with low divergence) when compared to this null distribution.

In practice, the distribution of donor-recipient divergence can be influenced by other factors, such as the uneven distribution of transfers along the genome (e.g. recombination hotspots) or potential biases in the transfer detection step. To account for these effects, we simulated transfers with the genomic locations as the observed transfers, and applied the CP-HMM algorithm to the simulated data. The detailed simulation algorithm is as follows:

1. For each pair of closely related strains with observed transfers, we randomly chose one of the two strains as the focal genome, and the other as the reference genome.
2. For each observed transfer between this pair, we sampled a genome segment with the same start and end positions from a random strain of the same species. We then overwrote the corresponding regions of the focal genome with sampled segments. If the donor segment was completely identical to the recipient sequence (e.g. from another closely related strain), we repeated the sampling step.
3. The SNV differences between the focal genome and the reference genome now contain a mixture of SNVs in the clonal regions and SNVs imported from simulated transfers. We then applied our CP-HMM algorithm to the simulated SNV profile to obtain a corresponding set of inferred transfers. The divergence of each of these inferred transfers were then added to the null distribution.
4. We repeated steps 1-3 for 5 repetitions per close pair to ensure that the null distribution was sufficiently sampled.
5. We then repeated steps 1-4 for all close pairs across all species in Fig. 3.

The comparisons between the observed distribution of donor-recipient divergences and the simulated null model are shown in Figs 4 & S18. We found that the observed distribution closely tracks the simulated distribution in the majority of species, with only a few notable exceptions discussed in the main text. These results indicate that the typical donor-recipient divergence in our panel is not sufficiently high to suppress the recombination rate on its own, and that other incompatibilities are likely needed to explain the observed genetic isolation between clades in *B. vulgatus* and *B. finegoldii*.

**_3.7_ Estimating the ratio between** *T*_mrca_ **and** *T*_mosaic_

One of the immediate consequences of being able to identify recombined regions is that we can now estimate the ratio *T*_mrca_/*T*_mosaic_ for every species with enough close pairs. Recall that *T*_mosaic_ is the time it takes for the genome of a strain to be completely overwritten by recombination events, while *T*_mrca_ is the average coalescence time between a pair of genomes. The ratio between these two timescales characterizes the degree of clonality or quasi-sexuality of a given bacterial population.

It is useful to connect this ratio to basic biological parameters in the context of a simple neutral model with a well-mixed population. Let *μ* and *r* be the per site per generation mutation and recombination rate, *l*_*r*_ the typical recombination length, *L* the genome length, and *d* is the typical pairwise divergence. Then, we can express *T*_mrca_ and *T*_mosaic_ in those parameters:

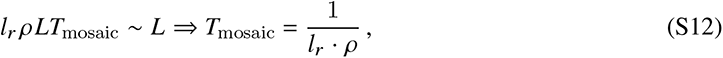

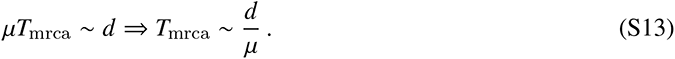

Therefore, in this simplest neutral model, we have

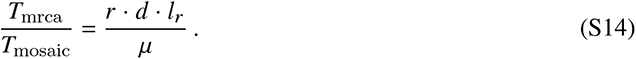

One can also interpret this quantity as a ratio between the number of mutations obtained through recombination in a given generation vs the number of mutations obtained through mutation (often referred to as “*r*/*m*” in previous work [25, 39]). This quantity differs from the bare ratio *r*/*μ* by the additional factor *d* · *l*_*r*_, which represents the typical number of mutations introduced per recombination event.

Empirically, the connection between *T*_mrca_/*T*_mosaic_ and underlying biological parameters is less straight- forward. Part of the reason arises from our finding that in the presence of population structure, *l*_*r*_, *r* and *d* are no longer a single set of numbers. Formally, we would need to sum over all these heterogeneous sources of recombination transfers to calculate *T*_mosaic_.

However, in practice, our understanding of the extent of this heterogeneity is limited. For species like *B. vulgatus*, we could infer two sets of these parameters to account for within- and between-clade transfer separately, as demonstrated in the main text. But for the majority of species with complex population structures, such as *A. putredinis*, it is much harder to even enumerate the types of transfers we need to consider separately. Potentially, there could be a continuum distribution of these recombination parameters, as argued in [28].

Nevertheless, the *T*_mrca_/*T*_mosaic_ ratio can still be studied empirically without explicit inference of the above biological parameters. The idea is that we can estimate the actual time it takes for a strain to be fully covered by recombination transfers, irrespective of the underlying heterogeneity. This time, *T*_mosaic_, will vary for different strains and could be highly random, but on average it should reflect the characteristic level of recombination of a given species.

Concretely, for each closely related pair, our CP-HMM algorithm can identify the total fraction of recombined genome, *f*_*r*_, as well as the clonal divergence, *d*_*c*_. If we assume that both *f*_*r*_ and *d*_*c*_ increase linearly in divergence time *T*, then

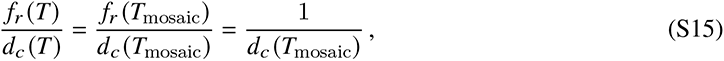

where we have used the fact that *f*_*r*_ (*T*_mosaic_) = 1. This implies that

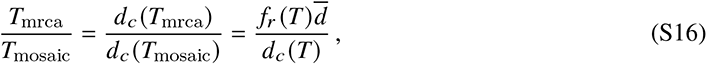

which allows us to estimate *T*_mrca_/*T*_mosaic_ for each pair of closely related strains.

In Fig. S16, we plot the distribution of these *T*_mrca_/*T*_mosaic_ estimates for all close pairs across a panel of different species. These estimates can also be viewed as a distribution of “*r*/*m*” values. We see that these estimates can vary significantly within certain species, potentially reflecting more than a single recombination rate or other deviations from the simple neutral model above. But the broad distribution within species does not overshadow the systematic differences between species. We can obtain an average *T*_mrca_/*T*_mosaic_ ratio for each species by calculating an average *d*_*c*_ (*T*_mosaic_):

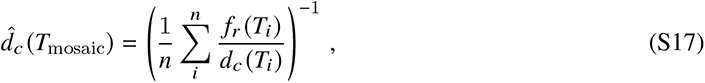

and substituting this value into Eq. (S16):

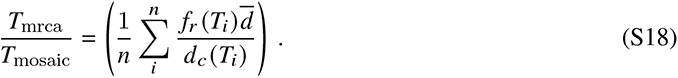

The resulting estimates for each species are shown in Fig. S16.

#### 3.8 Estimating the number of recombination events in a single gut microbiome

We can gain additional intuition for the recombination rate estimates in Fig. 3 by extrapolating them to the scale of an individual gut microbiome. For example, one important quantity is the total number of recombinants that are produced within a given species population each day. This quantity can be estimated from the product

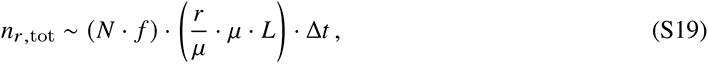

where *N* is the total number of bacterial cells in the gut microbiome, *f* is the relative abundance of the species, *μ* is the per site mutation rate, *r* is the corresponding recombination rate, *L* is the total length of the genome, and Δ*t* is the number of generations that take place per day. If we assume that successful recombination events accumulate largely neutrally (or via neutral hitchhiking), then the relevant *r*/*μ* values can be estimated from the apparent rates of accumulation in Fig. 3D; these range from 0.01−1, with most species clustering near *r*/*μ*∼0.1. Using estimates of the remaining parameters from the existing literature [*N*∼10^13^−10^14^ [110], *μ*∼10^−10^−10^−9^ [111], *L*∼10^6.5^ [112], Δ*t*∼1−10 [91, 113]], we expect that a moderately abundant species (*f* ∼3−30%) will produce anywhere from 10^6^ to 10^12^ recombinant offspring each day.

To convert this estimate to a per-site rate, we note that each site will have a probability ℓ_*r*_ /*L* of being covered by a given recombination event, where ℓ_*r*_ is the typical length of a transferred fragment. Using the typical lengths inferred from Fig. 3E (ℓ_*r*_ ∼10^3.5^ − 10^4.5^), we conclude that each site in the genome will be involved in 10^3^−10^10^ unique transfers each day.

To convert these population-level estimates to a per-lineage rate (e.g. to estimate the total number of recombination events that are expected within a single isolate genome), we need to drop the factor of *N f* in Eq. (S19). This implies that a typical genome will accumulate homologous recombination events at a total rate

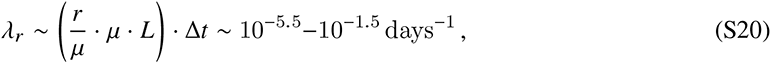

with most genomes clustered near the middle of this range (*λ*^−1^∼3000 days, or ∼ 10 years).

All of these extrapolations assume that recombination events accumulate largely neutrally (or via neutral hitchhiking) when averaged over multiple host colonization cycles. If this assumption is violated, than the apparent rates of accumulation in Fig. 3D will differ from the true value of *r*/*μ* by an additional factor of *γ* = *N p*_est_, which represents the average “establishment probability” of a newly acquired fragment. If the between-host patterns are driven by widespread positive selection (*N p*_est_ ≫ 1), then the true values of *r*/*μ* will be much smaller than the apparent values in Fig. 3D. Conversely, if most recombination events are negatively selected (*N p*_est_ ≫ 1), then the true values of *r*/*μ* will tend to be larger than the apparent values in Fig. 3D. Understanding the relative contributions of these selective forces is an interesting topic for future work.

#### 3.9 Deduplicating detected recombination events

One limitation of our pairwise detection method is that a single recombination event could be detected in multiple pairs. As a concrete example, consider three closely related strains, A, B and C, that have diverged very recently *T*_*ABC*_ ≪ *T*_mosaic_. Suppose A and B have diverged more recently from a common ancestor at *T*_*AB*_ < *T*_*ABC*_ and this common ancestor acquired a recombined fragment between *T*_*AB*_and *T*_*ABC*_. This recombination event would then show up *twice* when we compared the genomes of pairs AC or BC. On the other hand, if both A and B acquired different recombined fragments after they diverged, we would also see two events when we compared AC and BC.

For most of our pairwise analyses, we do not need to distinguish these two scenarios as long as we average across pairs. However, if we are interested in non-pairwise statistics, such as how frequently one region is transferred among many strains, we need to identify unique recombination events to minimize the impact of sampling bias. For example, if the common ancestor of A and B in the example above happened to have many more descendants in the sampled dataset, the same recombination event would be vastly over-represented if we naively retained all detected events.

In some cases, it is possible to infer the clonal relationships between strains using regions not affected by recombination and leverage the genealogical tree to identify unique events along the branches (e.g. Gubbins [29] or RecHMM [53]). Here, we adopt a simpler but approximate approach to deduplicate detected events using their genomic locations and sequence identity. The basic idea is that genomes that share the same transfer event by descent will show up in different pairs at approximately the same start and end locations with approximately the same sequence identity. This approach is conservative in that it will merge separate transfer events of the same region of the genome with the same donated haplotype. However, we assume that it is unlikely for two independent transfer events to have nearly identical start and end locations, and thus we expect these cases will have a small impact on the overall patterns of recombination that we detect.

More specifically, we first group together all transfers by their start and end locations, treating them as candidate duplicates if their start and end locations are in the same genome bin of 1000 sites. Next, we examine all the imported SNVs within the transfer region, and compute the fraction of SNVs that are identical between detected events. We consider two events to be duplicates if they have similar start and end locations (defined by the first step) and share > 98% of imported SNVs. This criterion is stricter than simply requiring a high sequence identity between the transferred fragments, since the sequences are always only a few percent diverged within a particular species.

We applied this deduplication procedure to all the detected events in Fig. 3 and found a total of 57,179 approximate unique events. We have included the uniqueness of each event in Table S3 and used only unique events in the analysis of transfer lengths (Fig. 2D and Fig. 3E). These unique events allowed us to study whether some genomic locations are biased toward more frequent recombination events (Fig. S30). We found that while recombination events can be detected throughout the genome, the landscape of detected transfers varies significantly along the genome, similar to the sharing landscape Fig. 6. Additional examination suggests that at least some of the peaks in the landscape coincide with putative recombination hot spots (Fig. S32C, [73, 74]). Interestingly however, when aggregated across the genome, we observed only weak correlations between the transfer landscape and the sharing landscapes across species (Fig. S30C). This lack of correlation suggests that these two landscapes can provide complementary information about the complex interplay between recombination and natural selection.

#### 3.10 Application to isolate genomes

We obtained alignments of high quality, isolate genomes of nine bacterial species from the Unified Human Gastrointestinal Genome (UHGG) collection [92] (Table S7), including five gut commensals in our metagenomic panel and four additional pathogens. We applied our CP-HMM model in the same fashion as our analysis of quasi-phased metagenomic genomes and repeated a number of analyses on the inferred transfer events analogous to Figures 1, 2, and 3.

Many of our results are reproduced using isolate genomes. We showed that the patterns of recombination barrier in B. vulgatus (Fig. 2) is quantitatively reproduced in the isolate analysis (Fig. S13AB) – among 40 close pairs, we detected 239 within-clade transfers and 52 between-clade transfers, consistent with the 5-fold reduction observed in the main text analysis. Other quantitative validations include the rate of accumulation of recombination events (Fig. S15 and Fig. S13F-I) and the median transfer lengths (Fig. S13K). We also reproduced qualitative results such as the significant variation in the sharing landscape (Fig. 6, Fig. S13J); and interesting examples of a putative within-host recombination event and a large recombination event (Fig. S13C), suggesting the generality of the events in Fig. 5 and Fig. S12.

##### 3.10.1 Comparisons with previous studies of bacterial pathogens

Our analysis of the pathogen species in Fig. S32 allows for a more direct comparison of our methods with the results of previous studies. For example, by analyzing the locations of the detected transfers in *Campylobacter jejuni* (Fig. S32BC), the most deeply sampled pathogen species in our dataset, we were able to reproduce the exact locations of recombination hot spots previously identified by the orderedPainting algorithm in Ref. 73. We also used the data from Ref. 48 to obtain a pairwise recombination fraction vs clonal divergence plot (analogous to Fig. S15), and found these data to be broadly consistent with our CP-HMM results (Fig. S32B). Comparing species-level summaries of recombination parameters like *r*/*m* is more challenging, however.

We compiled *r*/*m* estimates (or close equivalents) from two previous studies [25, 33] for the subset of pathogen species we analyzed in Fig. S32 (Table S1). These data show that the estimated values of *r*/*m* can differ by orders of magnitude in different studies — even the rank ordering of *r*/*m* for different species can vary widely. For example, Ref. 25 reported an *r*/*m* value for *Helicobacter pylori* that is >40 times larger than *Klebsiella pneumoniae*, but Ref. 33 estimated this value to be >20 times smaller than the *K. pneumoniae* rate. Similarly, our CP-HMM analysis suggests that the *r*/*m* value for *C. jejuni* is >30 times greater than that of *Salmonella enterica*, while Ref. 25 reported it to be ∼13 times smaller.

Some of these discrepancies are expected due to the heterogeneity in recombination rate within species, as demonstrated both by our current results as well as a large number of previous studies [28, 60, 61, 82, 83]).

**Table S1.** Comparison of inferred recombination rates for several pathogen species.

| Species name | $r/m$ [25] | $\rho = \phi_{pool} \bar{f}$ [33] | $T_{mrca}/T_{mosaic}$ |
| --- | --- | --- | --- |
| *Vibrio parahaemolyticus* | 39.8 | - | 18.3 |
| *Salmonella enterica* | 30.2 | - | 0.9 |
| *Helicobacter pylori* | 13.6 | 148 | - |
| *Campylobacter jejuni* | 2.2 | 200 | 39.5 |
| *Klebsiella pneumoniae* | 0.3 | 3248 | 23.1 |
| *Staphylococcus aureus* | 0.1 | 23.1 | - |

For example, in the case of *S. enterica*, our pairwise estimates reveal that some of the discrepancies might be caused by the fact that a small number of pairs with low clonal divergence acquired a large amount of transferred DNA, while a large number of pairs showed no evidence of recombination (Fig. S32G, H). This heterogeneity is consistent with previous findings that the *r*/*m* of *S. enterica* can differ by 10-fold between sub-lineages [61]. Summarizing such wide variation with a single value of *r*/*m* is inherently difficult, and will sensitively depend on the composition of the sample and the averaging scheme employed. In more sophisticated inference methods, this average scheme is not defined explicitly, but emerges through a complex process of likelihood optimization of an underlying parametric model. A potential benefit of semi-parametric approaches like CP-HMM (and related methods [42, 48]) is that they decouple the detection of individual recombination events from the averaging step at the end. By directly examining scatter plots like Fig. 2 and Fig. 3, it is possible to gain more insight into how different features of the data contribute to a given *r*/*m* estimate, and to more easily compare results across studies (e.g. Fig. S32B).

### 4 Identifying within-host recombination among co-colonizing strains

The within-host diversity of a given species can vary widely from host-to-host, and also within the same host over time [27]. In the previous two sections, we focused on the simplest subset of samples, where a single strain is present at high frequencies. The site frequency spectra (SFS) of these “quasi-phaseable” (QP) samples contain very few alleles at intermediate frequencies (top left of Fig. S1), so that the haplotypes of the dominant strains can be inferred with a high degree of confidence [27].

In this section, we revisit a subset of the remaining non-QP samples – which by definition contain multiple co-colonizing strains – to identify signatures of recent or ongoing recombination within hosts. Frequently, a non-QP sample features a single intermediate peak in the SFS, indicating the coexistence of two diverged strains. The basic signature of recombination between these strains will look like the inverse of the close-pair analysis above: rather than looking for regions of high divergence among closely related pairs, we search for regions of low divergence (i.e. identical DNA sequence) within a normally diverged background (*d*∼1%). These regions of shared DNA sequence will manifest as a local depletion of genetic diversity in the larger metagenomic sample (Fig. 5).

#### 4.1 Sample selection

In the idealized scenario of two coexisting clonal populations, all fixed differences between the two strains should be found at a certain intermediate frequency corresponding to the relative frequency of the two strains. Thus, in principle, we should be able to readily identify all fixed differences between two strains in these dual-colonized samples.

However, as shown on the left of Fig. S1, the observed SNVs usually form a broader distribution around the peak frequency. Sampling noise from finite sequencing coverage is likely a major contributor to this spread, but the large widths observed in some high coverage samples suggests additional processes might be involved. In either case, when the shoulder of the intermediate frequency peak merges with the main peak at 100% frequency, we can no longer reliably detect all fixed differences between the two co-colonizing strains. This could lead to spurious signals of recombination where none truly exist.

To avoid this difficulty, we restricted our attention to a particularly simple subset of dual-colonized samples which were identified by two criteria: (i) the SFS should have a single, pronounced peak at intermediate frequency, and (ii) the intermediate peak should be clearly separated from the main peak. We can then use this clear separation to reliably identify fixed differences (or the lack of fixed differences) between the pair of co-colonizing strains. To implement these criteria programmatically, we first smoothed the SFS with a Savitzky-Golay filter and detected peaks using the SciPy signal processing library [103]. We next quantified the quality of separation between peaks by computing the trough-to-peak ratio. Only samples with a ratio smaller than 0.2 were deemed to be simple enough for further analysis. The location of the trough was then used as a cutoff to classify sites as fixed differences between two strains (or not). Examples of simple and complex co-colonized samples are shown in Fig. S1.

We note, however, that this frequency-based classification will only for sites that are present in both of the co-colonizing strains. If a site is present in only one strain (e.g. due to a gene loss event), then a mutation can appear to be fixed within the remaining read alignments without actually being fixed in the population. To reliably filter such gene loss events, we restricted our attention to an even smaller subset of high quality samples where the noise in gene copy numbers was also minimal. Fig. S19 illustrates an example of this additional processing step for an example *B. vulgatus* population. We first computed the median read depth for each of the core genes (*D*_*i*_, where *i* is the gene index), and then calculated a moving average of 500 genes to estimate the large-scale depth variation due to bacterial growth dynamics [114] (Fig. S19 top). Next, we used this average variation (⟨*D*⟩_*i*_) to compute the relative copy number of each gene, defined as *c* = *D*_*i*_/⟨*D*⟩_*i*_.

If a gene is present in both strains, its copy number should remain close to one; if the gene is present in only one of the co-colonizing strains, its copy number should be close to the corresponding strain frequency (as inferred from the peak of the SFS, Fig. S19 middle). We computed the standard deviation of the copy number across all genes (*σ*_*c*_) to quantify the expected variation due to sequencing noise, and we used this estimate to compute a corresponding Z-score for each gene,

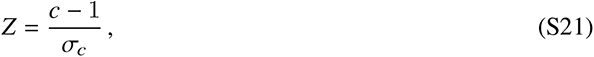

which quantifies the magnitude of the deviation from unit copy number. Genes with |*Z* | > 2 are filtered from our downstream analysis. In cases where the sequencing noise is large enough that a gene deletion event cannot possibly be distinguished from noise (1 − *f*_peak_ < 2*σ*_*c*_), we discard the entire sample from our downstream analysis. This procedure yields a set of co-colonized samples where we can confidently identify the set of genes that are present in both strains. We then classified the fixed differences and shared sites between the strains using the same frequency thresholds defined above.

Applying this procedure to our cohort yielded a total of 8 species (from 4 bacterial families) with ≥ 5 high-quality dual-colonization examples (Table S4). The two species with the largest sample sizes (*B. vulgatus* and *E. rectale*) are highlighted in Fig. 5.

#### 4.2 Identifying shared genomic regions among co-colonizing strains

The fixation of a successful recombination event will leave an extended run of zero fixed differences in the genomes of co-colonizing strains. To look for enrichment of such signals, we computed the lengths of all continuous stretches of 4D sites in the core genome that did not possess any fixed differences (a *homozygous run*). Filtered genes or other low coverage sites were omitted from this calculation. It is worth noting that any underlying recombination event could be longer than the measured run length if the transferred fragment also contained some accessory genes.

To interpret these measured run lengths, we first compared them to a null model in which the SNVs were randomly distributed across the high coverage sites in the genome. Let *d* be the pairwise divergence between a pair of strains. Then, the distance between two consecutive SNVs (*x*) follows a geometric distribution with mean (1 − *d*)/*d* ≈ 1/*d*, which can be approximated by a continuous exponential distribution when the distance is large. Thus, the probability of observing a homozygous run longer than ℓ under this null model is given by

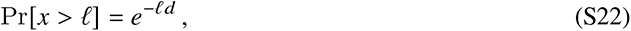

which was used to plot the null distributions in Figs. S20 and S23. This comparison revealed that co-colonizing strains can frequently share long homozygous runs that cannot be explained by the stochastic distribution of mutations along the genome (ℓ*d* > 25, or equivalently *P* ≲ 10^−11^). These shared genomic segments constitute potential candidates for within-host recombination events.

#### 4.3 Distinguishing within-host recombination from pre-existing sharing

To determine whether the shared genomic segments observed in co-colonizing strains were caused by previous within-host recombination events, we compared them to the baseline rates of sharing observed in random pairs of strains from unrelated hosts. We found that these random pairs can sometimes share segments that are just as long as those observed in co-colonizing strains (Fig. S20; see S1 Text 5.1 for a theoretical explanation). This suggests that it will be difficult to determine whether any *particular* shared segment was transferred before or after colonization. We therefore sought to quantify the global signatures of within-host recombination by examining the statistical enrichment of sharing among co-colonizing strains.

To perform this comparison, we took advantage of our large collection of QP strains and generated an empirical null distribution of runs of shared sequence between unrelated strains. We kept only one sample per host/household to approximate a random set of strains from the broader population. Since we are interested in recent recombination events between these strains, we want to minimize runs of shared sequence that arise through clonal inheritance of the entire genome (Fig. 1A). We therefore only analyzed “typically diverged” pairs with < 10% identical blocks in Fig. 1B. These pairs share a common ancestor long enough ago that imported fragments should have overwritten most of the clonal fraction, and any long sharing regions should be mostly due to subsequent recombination events between the ancestors of the pair. We then sampled a random subset of 5,000 between-host pairs for each species and computed the homozygous run lengths for each pair as described above; in cases where a species had fewer than 5,000 qualified pairs, we used all available pairs. There were two species that required additional consideration:

1. Anticipating the potential impact of the strong clade structure in *B. vulgatus*, we analyzed between-clade pairs separately from within-clade comparisons. To do so, we first clustered all the strains into two clades as described in S1 Text 2, and obtained the resulting within- vs between-clade strain combinations. Out of the 372 strains of *B. vulgatus* in our cohort, we identified 12,935 between-clade pairs and 16,711 within-clade pairs with “typical” levels of divergence.
2. *Eubacterium rectale* has a geographically structured population, which suggests that co-colonizing strains are sampled from a different distribution than the worldwide population. To account for this issue, we also need to control for geographical structure when selecting the between-host pairs for our null distribution. In our cohort, there are 15 co-colonized samples collected in the United States, and 13 samples collected in the United Kingdom. To reflect this geographical distribution, we sampled a subset of pairs from the US pool of *E. rectale* strains and another subset of pairs from the UK pool. We combined these subsets in a 15:13 ratio, yielding a total of 3000 strain pairs that reflect the geographical distribution of our co-colonized samples.

To compare these empirical null distributions with the observed within-host data, we used a test statistic based on the length of the longest homozygous region for each pair. The reverse cumulative distributions of these “max run” statistics for *B. vulgatus* and *E. rectale* are shown in Fig. 5D in the main text, and analogous distributions for the other species are shown in Fig. S22. We calculated P-values for the deviation between within-host and between-host distributions using a one-sided Kolmogorov-Smirnov test using the ks_2samp function in the SciPy package [103].

We also analyzed a complementary metric that sums all the runs longer than a given threshold ℓ^∗^. This metric is more suitable to detect the enrichmen<u>t</u> of *multiple* tr<u>an</u>sfer events between a given pair of strains. We choose the length threshold ℓ^∗^ such that ℓ^∗^ · *d* = 30, where *d* is the average pairwise divergence between the typically diverged pairs of that species (e.g. equivalent to ∼ 20kb in the case of *B. vulgatus*). This ensures that the sum is dominated by runs that likely reflect recent transfer events, rather than the random spacing of SNVs along the genome. Within- vs between-host comparisons of this new metric yielded results that were similar to the “max run” statistic above, with only *E. rectale* showing an enrichment of long runs in co-colonizing strains Fig. S21.

### 5 Identifying signatures of selection from the global distribution of recent transfers

#### 5.1 Neutral expectation for the probability of observing long shared fragments

Using the results in Ref. 72, one can derive an approximate formula for the expected number of long shared fragments in a simple neutral model with a constant population size. Consider a genome comparison between a random pair of strains. Let *H*(*l*) be the probability that a randomly chosen site will be a SNV and followed on the right by *l* identical sites. In general, *H*(*l*) can not be computed analytically because we need to consider all possible recombination histories within this shared segment. However, since recombination typically brings in SNVs that break a run of identical sites, we can expand *H*(*l*) in terms of the number of recombination break points it contains, under the assumption that contributions from shared segments spanning multiple ancestral recombination events become increasingly small [72]. For simplicity, here we only consider the contribution from the lowest order term, in which the shared segment contains zero ancestral recombination events (Ref. 72 computed this quantity for up to two crossover recombination events). In this case, the *H*(*l*) function can be approximated by

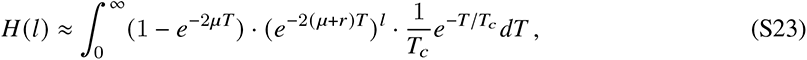

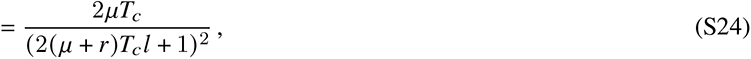

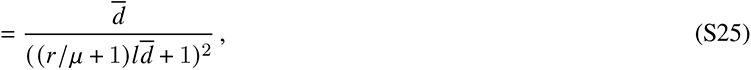

where *μ* and *r* are the per site, per generation mutation and recombination rates, *T*_*c*_ is the average coalescence time in the populatio<u>n</u>, and *d* = 2*μT*_*c*_is the average pairwise divergence. Since we are interested in long shared fragments (*l* · *d* ≫ 1), we can further approximate the above formula as

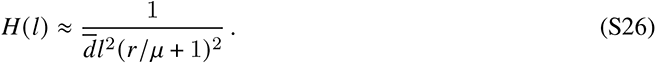

Using this formula, the expected number of shared fragments with length exactly *l* in a genome of length *L* is given by

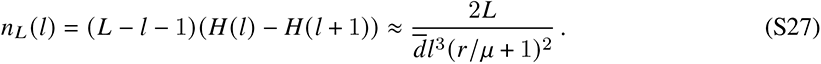

Finally, the total probability of observing a shared fragment longer than *l*^∗^ is given by

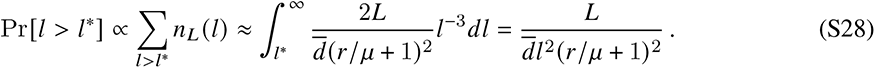

Note that this distribution has a heavier tail that the exponential distribution of run lengths expected from random mutations alone (Eq. S22). This shows that it is not surprising to observe long shared fragments between random pairs of strains in neutral, recombining populations. We verified this intuition using computer simulations (Fig. S23).

#### 5.2 Quantifying parallelism in recent transfers

While neutral models like Eq. (S28) predict that a given pair of genomes can share several long segments, they place strong constraints on the probability that a particular fragment will be shared across multiple independent pairs (see S1 Text 5.3 below). We quantified this signature of parallelism using the haplotype sharing metric illustrated in Fig. 6A. For each 4D site in the core genome, we recorded the fraction of pairs in which that site was contained within a run of length ≳ ℓ^∗^. W e call this normalized quantity the “sharing probability.” As above, we chose the threshold such that ℓ^∗^ · *d* = 15, in order to distinguish likely recombination events from random fluctuations in SNV spacing.

The resulting “sharing landscapes” in Fig. 6 were calculated using the same set of between-host pairs that were previously used as the null distribution in Figs. 5D and S22 (S1 Text 4.3). In particular, we (1) used only typically diverged pairs (< 10% identical blocks), (2) analyzed between- and within-clade pairs of *B. vulgatus* separately, and (3) controlled the geographical structure of *E. rectale* as before.

Compared to neutral expectations (Fig. S2), a greater fraction of closely related pairs is observed in some species (e.g. *Bacteroides caccae*, Fig. S3), potentially reflecting certain sampling biases in the dataset. To minimize the impact of the over-representation of these closely related strains, we clustered strains with fraction of identical blocks > 95% and keep only one representative from each cluster.

We used a similar approach to quantify parallel transfers in pairs of co-colonizing strains as well (Fig. 6C), using the homozygous runs identified in S1 Text 4.2. For hosts with multiple time points (e.g. the within-host sweep example in Fig. 5C), we only included the comparison from the last time point in order to avoid counting the same pair twice.

##### 5.2.1 Connections to selection tests based on haplotype homozygosity

Our sharing probability metric is closely analogous to statistics based on haplotype homozygosity [99, 115], with a few key differences. For bacteria, two random genomes can share a haplotype either through clonal inheritance or a recent recombination event containing the haplotype. Haplotype homozygosity (and its close relatives) measures the total probability of these two modes of sharing. In contrast, our sharing probability statistic focuses on the second mode of sharing by only examining pairs with minimal clonal regions (i.e. typically diverged pairs as in Fig. 5B, rather than close pairs as in Fig. 2B). When none of the genome pairs share clonal regions, we expect the sharing probability to be equivalent to haplotype homozygosity (e.g. Fig. 6A), where the length threshold above, ℓ^∗^, serves the role of the window size in identifying the local haplotype spectra. However, if some of the pairs share large fractions of clonal ancestry (>10%), then the sharing probability will generally differ from the local haplotype spectrum. We therefore expect our pairwise sharing probability statistic to be better suited for detecting recent transfers than haplotype homozygosity alone.

To better illustrate the connection with existing statistics, we compared the within-clade sharing landscape of *B. vulgatus* with genome-wide scans of haplotype homozygosity (Fig. S27). As expected, the variations of haplotype homozygosity closely follow the sharing landscape along the genome, and all statistics are strongly elevated at regions with a single, abundant haplotype (Fig. S27D,E). However, the sharing landscape also contains peaks at regions where multiple haplotypes are circulating at lower frequencies (Fig. S27F), which are harder to detect by simple versions of haplotype homozygosity statistics (Fig. S27B). It is worth noting that by itself, the sharing probability statistic does not distinguish elevated values caused by one haplotype or multiple. Explicit analysis of haplotypes, or alternatively, statistics that account for the contribution of the most dominant haplotype (Fig. S27B), can help distinguish these scenarios.

#### 5.3 Comparison to simulated data from simple neutral models

To calculate the expected sharing landscapes of a neutral population, we performed simulations using the simulator FastSimBac [116]. FastSimBac simulates a recombining population and is capable of generating haplotype sequences for a large number of genomes. This allows us to compute statistics involving multiple strains (e.g. how frequent a region is shared by multiple pairs), which cannot be obtained by our simpler pairwise simulations in S1 Text 3.2. The simulations in Fig. 6B were run with parameters that were chosen to match the observed data from *B. vulgatus*. We set the genome length to be 2.8 × 10^5^ (which corresponds to the synonymous core genome length of *B. vulgatus*) and simulated a sample size of 200 genomes. We set the scaled mutation rate *θ*, to be equal to the within-clade diversity of *B. vulgatus* (*d*_*within*_ ≈ 0.008). We chose the recombination length to be approximately equal to the average recombination length inferred for *B. vulgatus* in Fig. 2 (*λ* = 2000), and scanned over a range of scaled recombination rates (*ρ*/*θ* ∈ {0, 0.1, 0.2, 0.5, 1, 2, 4}). Each parameter was simulated for 100 replicate populations with a constant population size. Each of these simulation runs generated a sample of 200 genomes, which we analyzed in exactly the same way as the between-host analysis above. In particular, we only used typically diverged pairs (i.e. those with an identical fraction < 10%) when calculating the resulting sharing landscapes. This filter is important because some parameter combinations tended to produce a much larger fraction of pairs that inherited large clonal regions.

We computed the sharing landscape of these simulated populations for a range of threshold lengths (ℓ^∗^ · *d* = 10, 12, . . . 34). For each sharing landscape, we computed the median sharing probability as well as the coefficient of variation (CV) in sharing probability (Fig. S28). We found that adjusting the threshold length has roughly equivalent effects as adjusting *ρ*/*θ*: increasing both parameters would decrease the median sharing probability while leaving the CV almost unchanged. Since higher values of *ρ*/*θ* were more computationally expensive to simulate, we used this empirical scaling relation to cover a wider range of median sharing probabilities than would be feasible to simulate directly.

To compare the sharing distribution in simulations with that of *B. vulgatus* (Fig. 6C), we found the combination of effective ℓ^∗^ and *ρ*/*θ* values that matched the median sharing probability of the observed data (Fig. S28A). The resulting sharing landscape were shown in Fig. 6B.

To test the effects of a locally elevated recombination rate, we used the -R option of FastSimBac to specify regions of modified recombination rates. Figure S29 shows that elevated recombination rates actually *lower* the local sharing probability, which is consistent with the intuition that faster recombination generates more unique haplotypes. Conversely, Fig. S29 shows that lower recombination rates slightly increase the sharing probability, but the overall levels still remain much lower than the “hotspots” observed in *B. vulgatus*. Curiously, we also observe a non-monotonic increase of the local sharing probability as we lower the local recombination rates: a small decrease in recombination rate initially increases the local sharing probability, but larger decreases eventually eliminate this pattern. Understanding the origins of this non-monotonic behavior would be an interesting topic for future work.

#### 5.4 Quantifying the differences in the sharing landscapes of co-colonizing strains

The within-host sharing landscape of *E. rectale* deviates from its between-host counterpart at multiple locations along the genome (Fig. 6C). To quantify the significance of this trend, we compared the observed data to a null model in which the within- and between-host labels were randomly permuted across pairs. For each of these bootstrapped datasets, we computed the sharing landscape as above and recorded the *excess sharing* (defined as the difference between the sharing probabilities of within vs between host pairs) at each location along the core genome. By comparing the observed data with the null distribution of *n* = 5000 bootstrapped datasets, we obtained an array of (uncorrected) P-values for each location.

To adjust these P-values for multiple comparisons, we performed a second level of bootstrapping. For each of the permuted datasets, we computed an analogous array of P-values using the other *n* − 1 permutations as the null distribution. To obtain a global metric of the deviation from this null model, we computed the total number of sites with *P* < 0.01 in both the observed and simulated datasets. We found that the total number of sites with excess sharing was significantly longer in the observed data than in any of the bootstrapped datasets (*P* < 0.001).

This second level of bootstrapping also allowed us to identify local deviations from the null model. To do so, we took the observed and simulated P-value arrays from our analysis above and created smoothed versions using a sliding window of 1000 sites. We then recorded the *minimum* smoothed P-value across the genome for each simulated dataset. We selected the 5th percentile of this distribution as a threshold for assessing the smoothed P-values observed in the data: a smoothed P-value below this threshold is unlikely to have occurred anywhere in the genome under the null model (*P* < 0.05). This local metric enabled us to identify many individual regions along the genome where the within-host sharing probabilities were significantly larger than the corresponding between host distribution (*P* < 0.05; Fig. S31). Some enriched regions coincided with regions of moderate between-host sharing (e.g. near location 150,000), while others were entirely new (e.g. near location 90,000).

To further validate our approach, we repeated the above process using an additional permutated dataset that was not in the *n* simulated datasets above. As expected, this negative control showed no region in the genome that had significantly more excess sharing than the simulated datasets (Fig. S31). This result confirms that our two-level bootstrapping approach provides an adequate correction for multiple-hypothesis testing.

#### 5.5 Enrichment analysis of functional classes of genes

To test if certain classes of genes were enriched for frequent sharing across hosts, we compared the observed data to a null model in which the gene annotations were randomly permuted across genes. To preserve potential co-occurrence of gene clusters, we permuted annotations in groups of 10 consecutive genes. For each permutation, we recorded the genes in regions of frequent sharing (sharing probability > 10%), and computed the number of genes in the functional class of interest. We repeated this procedure for 10^5^ bootstrapped datasets, and defined a corresponding P-value based on the fraction of bootstrapped datasets that exceeded the observed value.

We applied this test to two candidate functional classes, glycosyltransferases and ribosomal proteins, which frequently appeared in the sharing hotspots of *B. vulgatus*. We found that for *B. vulgatus*, the glycosyltransferases were enriched for within-clade sharing (*p* < 10^−2^) while the ribosomal proteins were enriched for between-clade sharing (*p* ≈ 10^−3^). The spatial distribution of genes belonging to these two classes coincide with some of the most prominent sharing hotspots in *B. vulgatus*, suggesting that selection on these genes could play a major role in shaping the overall patterns of the sharing landscape. This observation is consistent with previous work in *B. fragilis*, which found that glycosyltransferases were frequent targets of selection within individual hosts [71]. Interestingly, ribosomal proteins have previously been shown to be associated with recombination “cold spots” in a number of other bacterial species [74]. This observation could be consistent with the scenario of recent local sweeps, which leads to a shorter coalescence time and less time to recombine. Further analysis is needed to establish the differences between the sharing hotspots examined here and the recombination “hot/cold regions” inferred by existing chromosome painting algorithms [73, 117]. Combining the information encoded in these different statistics could be an interesting avenue for future work.

